# Biophysics-based protein language models for protein engineering

**DOI:** 10.1101/2024.03.15.585128

**Authors:** Sam Gelman, Bryce Johnson, Chase Freschlin, Arnav Sharma, Sameer D’Costa, John Peters, Anthony Gitter, Philip A. Romero

## Abstract

Protein language models trained on evolutionary data have emerged as powerful tools for predictive problems involving protein sequence, structure, and function. However, these models overlook decades of research into biophysical factors governing protein function. We propose Mutational Effect Transfer Learning (METL), a protein language model framework that unites advanced machine learning and biophysical modeling. Using the METL framework, we pretrain transformer-based neural networks on biophysical simulation data to capture fundamental relationships between protein sequence, structure, and energetics. We finetune METL on experimental sequence-function data to harness these biophysical signals and apply them when predicting protein properties like thermostability, catalytic activity, and fluorescence. METL excels in challenging protein engineering tasks like generalizing from small training sets and position extrapolation, although existing methods that train on evolutionary signals remain powerful for many types of experimental assays. We demonstrate METL’s ability to design functional green fluorescent protein variants when trained on only 64 examples, showcasing the potential of biophysics-based protein language models for protein engineering.

## Introduction

Just as words combine to form sentences that convey meaning in human languages, the specific arrangement of amino acids in proteins can be viewed as an information-rich language describing molecular structure and behavior. Protein language models (PLMs) harness advances in natural language processing to decode intricate patterns and relationships within protein sequences [1]. These models learn meaningful, low-dimensional representations that capture the semantic organization of protein space and have broad utility in protein engineering [2]. PLMs can be adapted to specific protein properties like enzyme activity or stability with limited training examples [3, 4], and they can be used in predictive or generative settings to design custom-made proteins with desired characteristics [5, 6].

PLMs such as UniRep [7] and Evolutionary Scale Modeling (ESM) [8] are trained on vast repositories of natural protein sequences distributed across the evolutionary tree. The training process typically involves self-supervised autoregressive next token prediction or masked token prediction [1, 9]. Through this process, PLMs learn context-aware representations of amino acids within proteins. Training on examples of natural proteins produces PLMs that implicitly capture protein structure, biological function, and other evolutionary pressures. While these models are powerful, they do not take advantage of the extensive knowledge of protein biophysics and molecular mechanisms acquired over the last century, and thus, they are largely unaware of the underlying physical principles governing protein function.

We introduce Mutational Effect Transfer Learning (METL), a PLM that integrates biophysical knowledge during pretraining before being finetuned with experimental data for protein engineering applications. Unlike evolutionary-based PLMs, METL is pretrained on biophysical data generated through molecular simulations across diverse protein sequences and structural folds. METL captures biophysical relationships inherent in these molecular simulations and learns a biophysically-grounded protein representation. This biophysics-informed approach allows METL to understand and predict protein function based on underlying biophysical mechanisms, offering insights that can complement traditional evolutionary-based models.

Following pretraining, we finetune METL using experimental sequence-function data, producing biophysics-aware models that can predict specific protein properties. Experimental data plays a critical role in protein engineering by providing direct, empirical relationships between sequence variations and observed functional outcomes. In contrast to zero-shot models that rely solely on pretrained knowledge or *de novo* models that build completely new proteins from scratch, METL uses experimental data to explicitly predict how sequence changes influence protein function. METL excels in protein engineering tasks like generalizing from small experimental training sets and extrapolating to mutations not observed in the training data. We demonstrate METL’s ability to design functional green fluorescent protein (GFP) variants when trained on only 64 sequence-function examples. METL establishes a general framework for incorporating biophysical knowledge into PLMs and will become increasingly powerful with advances in molecular modeling and simulation methods.

## Results

### Pretraining protein language models with synthetic data

Deep neural networks and language models are revolutionizing protein modeling and design, but these models struggle in low data settings and when generalizing beyond their training data. Although neural networks have proven capable in learning complex sequence-structure-function relationships, they largely ignore the vast accumulated knowledge of protein biophysics. This limits their ability to perform the strong generalization needed for protein engineering, which is the process of modifying a protein to improve its properties [10]. We introduce a framework that incorporates synthetic data from molecular simulations as a means to augment experimental data with biophysical information (Fig. 1). Molecular modeling can generate large datasets revealing mappings from amino acid sequences to protein structure and energetic attributes. Pretraining on this data imparts fundamental biophysical knowledge that can be connected with experimental observations.

**Figure 1.**
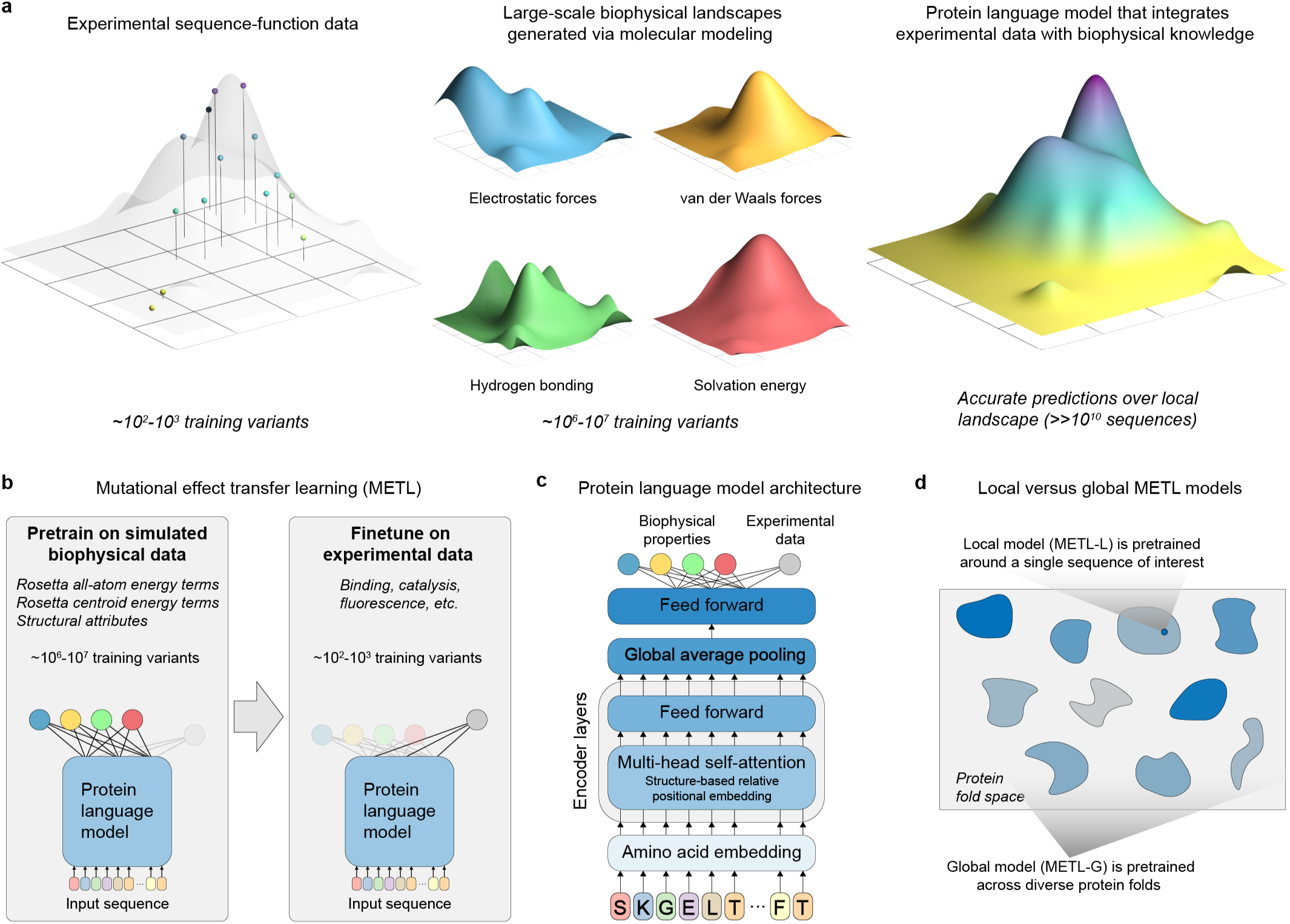
Mutational Effect Transfer Learning (METL). **(a)** METL combines sparse experimental protein sequence-function data with dense biophysical simulation data to learn biophysics-informed sequence-function landscapes. **(b)** The pretraining phase involves generating millions of protein sequence variants and computing biophysical attributes for them with Rosetta, which are then used to pretrain a protein language model. The model is subsequently finetuned with experimental sequence-function data to predict protein properties such as binding, enzyme activity, thermostability, and expression. **(c)** The METL architecture consists of a transformer encoder with a structure-based relative position embedding. **(d)** METL-Local and METL-Global differ in the sequences included in the pretraining data. METL-Local trains on the local sequence space around a protein of interest, learning a representation specific to that protein. METL-Global trains on diverse sequences across protein fold space, learning a general-purpose protein representation.

We introduce the METL framework for learning protein sequence-function relationships. METL operates in three steps: synthetic data generation, synthetic data pretraining, and experimental data finetuning. First, we generate synthetic pretraining data via molecular modeling with Rosetta [11] to model the structures of millions of protein sequence variants. For each modeled structure, we extract 55 biophysical attributes including molecular surface areas, solvation energies, van der Waals interactions, and hydrogen bonding (Table S1). Second, we pretrain a transformer encoder [12] to learn relationships between amino acid sequences and these biophysical attributes and to form an internal representation of protein sequences based on their underlying biophysics. The transformer uses a protein structure-based relative positional embedding [13] that considers the three-dimensional distances between residues. Finally, we finetune the pretrained transformer encoder on experimental sequence-function data to produce a model that integrates prior biophysical knowledge with experimental data. The finetuned models input new sequences and predict the particular property learned from the sequence-function data.

We implement two pretraining strategies, METL-Local and METL-Global, that specialize across different scales of protein sequence space (Fig. 1d). METL-Local learns a protein representation targeted to a specific protein of interest. We start with the protein of interest, generate 20M sequence variants with up to 5 random amino acid substitutions, model the variants’ structures using Rosetta, compute the biophysical attributes, and train a transformer encoder to predict the biophysical attributes from sequence. METL-Local demonstrates strong predictive performance on these attributes (Fig. S1a), achieving a mean Spearman correlation of 0.91 for Rosetta’s *total score* energy term across the eight METL-Local source models we trained. Although METL-Local accurately recapitulates the biophysical attributes, the primary purpose of pretraining is to learn an information-rich protein representation that can be finetuned on experimental data.

METL-Global extends the pretraining to encapsulate a broader protein sequence space, learning a general protein representation applicable to any protein of interest. We select 148 diverse base proteins [14] (Table S2) and generate 200k sequence variants with up to 5 random amino acid substitutions for each. We then model the approximately 30M resulting structures with Rosetta, extract biophysical attributes, and train a transformer encoder, following a similar methodology to METL-Local. With METL-Global, we observed a substantial difference in predictive ability for in-distribution structures (those included in the METL-Global pretraining data, mean Rosetta *total score* Spearman correlation of 0.85) and out-of-distribution structures (those not included, mean Rosetta *total score* Spearman correlation of 0.16) (Fig. S1b), indicating METL-Global overfits to the 148 base proteins present in the pretraining data. However, we find it still captures biologically relevant amino acid embeddings (Fig. S2) that are informative for protein engineering tasks even on the out-of-distribution proteins.

### Generalization of biophysics-based protein language models

Generalizing to new data is challenging for neural networks trained with small or biased datasets. This issue is crucial in protein engineering because experimental datasets often have few training examples and/or skewed mutation distributions. These factors impact the accuracy and utility of learned models when using them to design new protein variants.

We rigorously evaluated the predictive generalization performance of METL on 11 experimental datasets, representing proteins of varying sizes, folds, and functions: GFP, DLG4-Abundance (DLG4-A), DLG4-Binding (DLG4-B), GB1, GRB2-Abundance (GRB2-A), GRB2-Binding (GRB2-B), Pab1, PTEN-Abundance (PTEN-A), PTEN-Activity (PTEN-E), TEM-1, and Ube4b (Table S3). The METL-Global pretraining data contains proteins with sequence and structural similarity to DLG4, GRB2, and TEM-1 (Table S4), although their sequence identities are all below 40%. We observed no meaningful performance advantage for these proteins compared to others when using METL-Global to predict Rosetta scores (pre-finetuning) or experimental function (post-finetuning).

We compared METL to established baseline methods that provide zero-shot or standalone predictions, including Rosetta’s *total score*, the evolutionary model of variant effect (EVE) [15], and Rapid Stability Prediction (RaSP) [16]. We also evaluated supervised learning and finetuning methods, including linear regression with a one hot amino acid sequence encoding (Linear), an augmented EVE model that includes the EVE score as an input feature to linear regression in combination with the amino acid sequence (Linear-EVE) [17], a non-parametric transformer for proteins (ProteinNPT) [18], and the ESM-2 [19] PLM finetuned on experimental sequence-function data. We created comprehensive train, validation, and test splits, encompassing small training set sizes and difficult extrapolation tasks, and we tested multiple split replicates to account for variation in the selection of training examples.

We evaluated the models’ ability to learn from limited data by sampling reduced training sets and evaluating performance as a function of training set size (Fig. 2). The protein-specific models METL-Local, Linear-EVE, and ProteinNPT consistently outperformed the general protein representation models METL-Global and ESM-2 on small training sets. Among the protein-specific approaches, the best-performing method on small training sets tended to be either METL-Local or Linear-EVE, with METL-Local demonstrating particularly strong performance on GFP and GB1. While ProteinNPT sometimes surpassed METL-Local on small training sets, ProteinNPT was still generally outperformed by Linear-EVE in those instances. The relative merits of METL-Local versus Linear-EVE partly depend on the respective correlations of Rosetta *total score* and EVE with the experimental data. However, as the number of training examples increases, the METL-Local performance becomes dominated by dataset-specific effects rather than Rosetta *total score* relevance (Fig. S3). For the general protein models, METL-Global and ESM-2 remained competitive with each other for small to mid-size training sets, with ESM-2 typically gaining an advantage as training set size increased.

**Figure 2.**
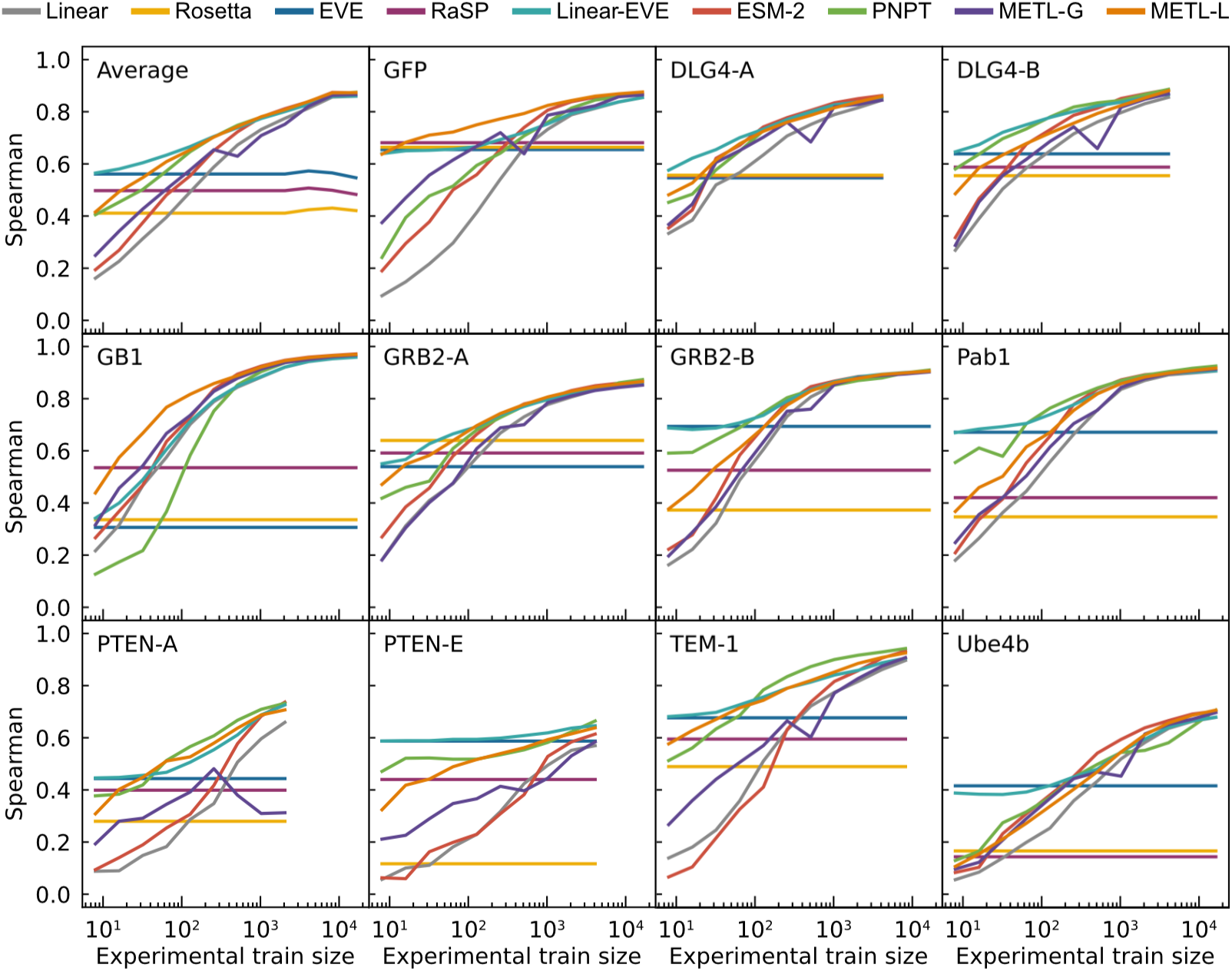
Comparative performance of Linear, Rosetta *total score*, EVE, RaSP, Linear-EVE, ESM-2, ProteinNPT, METL-Global, and METL-Local across different training set sizes. Learning curves for 11 datasets showing the test set Spearman correlation between true and predicted protein function scores across a number of training set sizes ranging from 8 to 16,384 examples. We tested multiple replicates for each training set size, starting with 101 replicates for the smallest train set size and decreasing to 3 replicates for the largest size. We show the median Spearman correlation across these replicates. The top left panel (“Average”) shows the mean of the learning curves across the 11 datasets.

We implemented four challenging extrapolation tasks — mutation, position, regime, and score extrapolation — to simulate realistic protein engineering scenarios, such as datasets lacking mutations at certain positions, having biased score distributions with predominantly low-scoring variants, and consisting of solely single-substitution variants (Fig. 3). Mutation extrapolation evaluates a model’s ability to generalize across the 20 amino acids and make predictions for specific amino acid substitutions not present in the training data [20] (Fig. 3a). The model observes some amino acid types at a given position and must infer the effects of unobserved amino acids. We found ProteinNPT, ESM-2, METL-Local, Linear-EVE, and METL-Global all performed well at this task, achieving average Spearman correlations across datasets ranging from ∼0.70 to ∼0.78. Position extrapolation evaluates a model’s ability to generalize across sequence positions and make predictions for amino acid substitutions at sites that do not vary in the training data [20–22] (Fig. 3b). This task is more challenging than mutation extrapolation and requires the model to possess substantial prior knowledge or a structural understanding of the protein [23]. ProteinNPT and METL-Local displayed the strongest average position extrapolation performance with Spearman correlations of 0.65 and 0.59, respectively. METL-Local’s success in mutation and position extrapolation relative to METL-Global is likely the result of the local pretraining data, which includes all mutations at all positions, providing the model with comprehensive prior knowledge of the local landscape.

**Figure 3.**
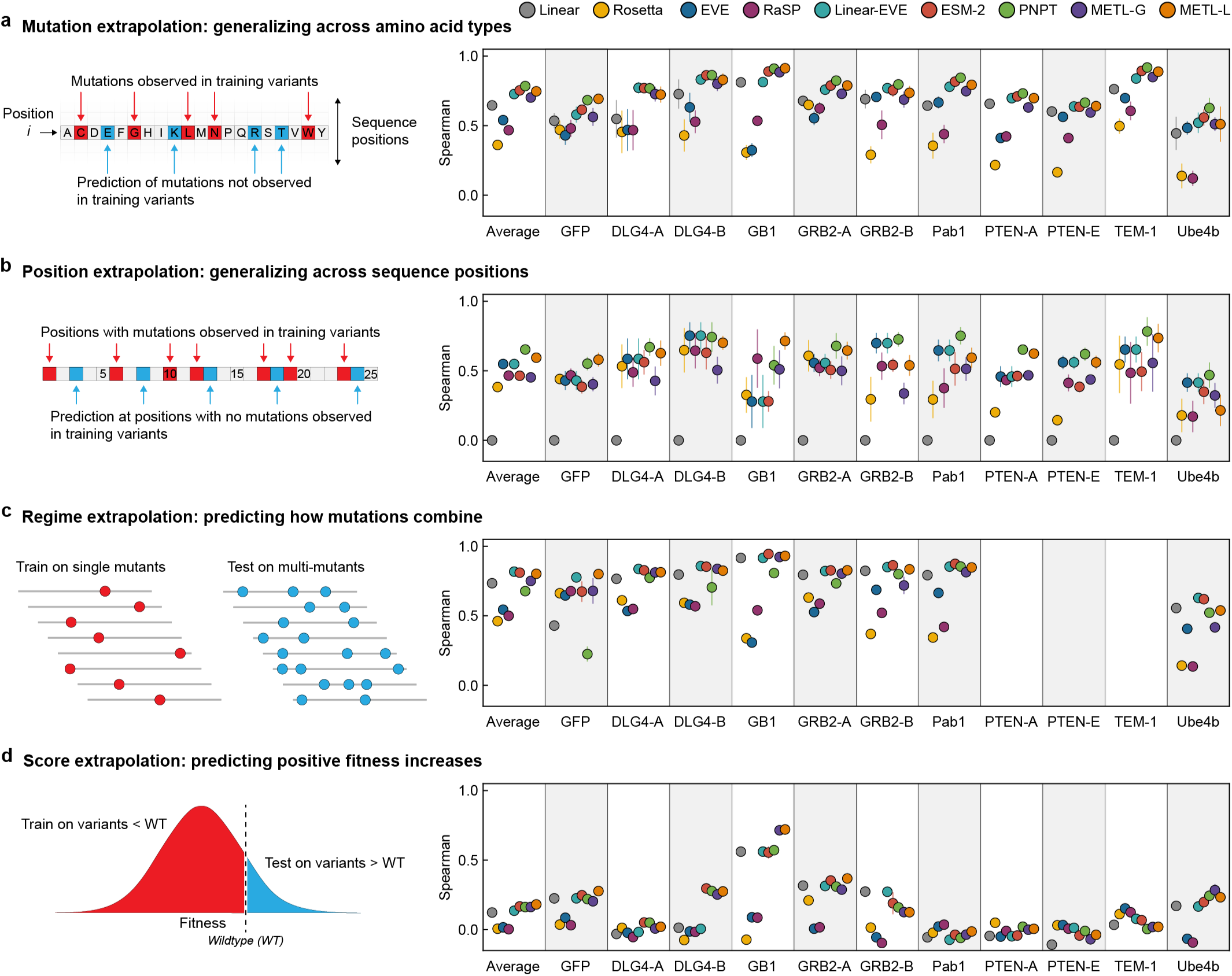
Comparative performance across extrapolation tasks. Correlation performance of Linear, Rosetta *total score*, EVE, RaSP, Linear-EVE, ESM-2, ProteinNPT, METL-Global, and METL-Local on (a) mutation, (b) position, (c) regime, and (d) score extrapolation. We tested 9 replicates for each type of extrapolation and show the median. Error bars indicate one standard deviation.

Regime extrapolation tests a model’s ability to predict how mutations combine by training on single amino acid substitutions and predicting the effects of multiple substitutions [21, 22, 24, 25] (Figs. 3c and S4). The supervised models generally performed well at regime extrapolation, achieving average Spearman correlations above 0.75. The strong performance of linear regression, which relies on additive assumptions, suggests the sampled functional landscape is dominated by additive effects. ProteinNPT performed slightly worse than the other supervised models, with an average Spearman correlation of 0.67, partly driven by lower performance on the GFP dataset. Score extrapolation tests a model’s ability to train on variants with lower-than-wild-type scores and predict variants with higher-than-wild-type scores [25] (Fig. 3d). This proves to be a challenging extrapolation task, with all models achieving a Spearman correlation less than 0.3 for most datasets. The GB1 dataset is an exception for which all supervised models achieved Spearman correlations of at least 0.55, and both METL-Local and METL-Global displayed correlations above 0.7. The difficulty of score extrapolation might be attributed to the fact that the mechanisms to break a protein are distinctly different than those to enhance its activity. It is notable that Rosetta *total score* and EVE, which are not trained on experimental data, performed worse at score extrapolation than they did at the other extrapolation tasks. This suggests these methods are largely capturing whether a sequence is active or inactive, rather than the finer details of protein activity.

We performed the above prediction and extrapolation tasks with several additional baselines, including METL-Local with random initialization (Fig. S5), augmented linear regression with Rosetta’s *total score* as an input feature (Fig. S6), and sequence convolutional networks and fully-connected networks (Fig. S7). METL-Local outperformed these additional baselines on nearly every prediction task for every dataset or provided much better scalability. We evaluated the recall of the top 100 test variants as an alternative metric (Fig. S8), which showed that strong Spearman correlation does not necessarily imply strong recall performance. Further, we conducted a systematic evaluation of the METL architecture to investigate one-dimensional (sequence-based) versus three-dimensional (structure-based) relative position embeddings (Fig. S9), feature extraction versus finetuning (Fig. S10), global model sizes (Figs. S11 and S12), and the extent of overfitting to the pretraining biophysical data (Fig. S13).

### Information value of simulated versus experimental data

METL models are trained on both simulated and experimental data. Generating simulated data is orders of magnitude faster and less expensive than experimental data. We wanted to understand how these two sources of data interact and if simulated data can partially compensate for a lack of experimental data. To quantify the relative information value of simulated versus experimental data, we measured the performance of the GB1 METL-Local model pretrained on varying amounts of simulated data and finetuned with varying amounts of experimental data (Fig. 4). Increasing both data sources improves model performance, and there are eventually diminishing returns for adding additional simulated and experimental data. The shaded regions of Fig. 4 define iso-performance lines with simulated and experimental data combinations that perform similarly. For instance, a METL-Local model pretrained on 1,000 simulated data points and finetuned on 320 experimental data points performs similarly to one pretrained on 8,000 simulated data points and finetuned on only 80 experimental data points. In this example, adding 7,000 simulated data points is equivalent to adding 240 experimental data points, and thus ∼29 simulated data points give the same performance boost as a single experimental data point.

**Figure 4.**
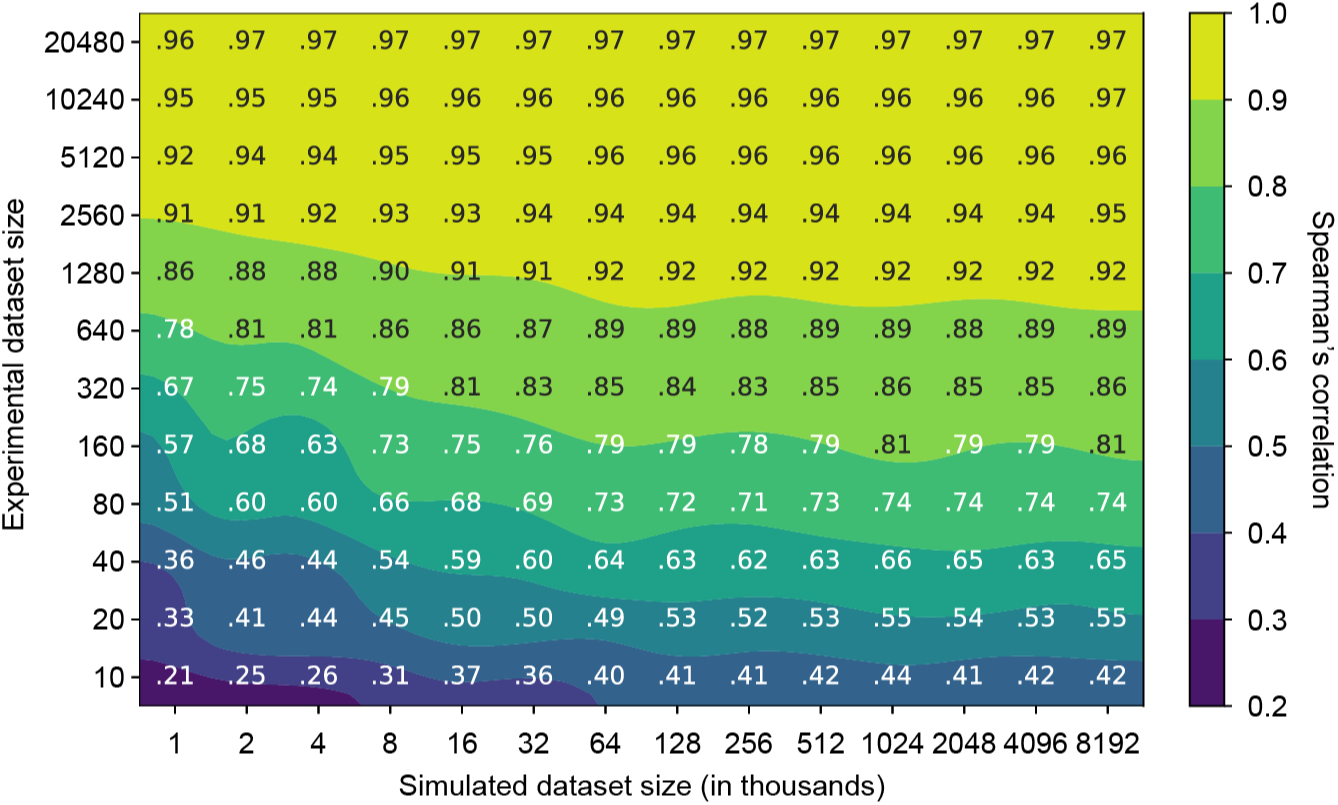
Relationship between experimental and simulated data quantities for GB1. The contour plot illustrates the test set Spearman’s correlation resulting from training METL-Local with varying amounts of simulated (pretraining) and experimental (finetuning) data. The plot displays a grid of Spearman’s correlation values corresponding to discrete combinations of experimental and simulated dataset sizes. The model benefits from larger quantities of experimental and simulated data, with the latter producing diminishing returns after approximately 128k examples.

We observe distinct patterns in how different proteins respond to increasing amounts of simulated pretraining data (Fig. S14). For larger proteins like GFP (237 residues), TEM-1 (286 residues), and PTEN (403 residues), we see a threshold effect wherein performance for a given experimental dataset size remains relatively flat until reaching a critical mass of simulated examples, at which point there is a sharp improvement in downstream performance. In contrast, smaller proteins like GB1 (56 residues), GRB2 (56 residues), and Pab1 (75 residues) show a more gradual response to increased simulated data over the tested dataset sizes. The performance gains are more modest, particularly when experimental data is abundant, but occur more consistently across the range of pretraining data sizes, until hitting a point of diminishing returns. A number of factors could influence this information gain phenomenon, including the protein’s size, the protein’s structural and functional properties, the experimental assay characteristics, and Rosetta’s modeling accuracy. Finally, we observe diminishing returns and saturated performance starting with simulated dataset sizes as small as ∼16K examples, depending on the protein and number of experimental examples. The point of diminishing returns occurs at a substantially smaller number of simulated examples than the ∼20M used for our main results, suggesting that less simulated data could be used to train METL-Local in practice.

### Synthetic data pretraining imparts biophysical knowledge

The purpose of METL’s pretraining is to learn a useful biophysics-informed protein representation. To further probe METL’s pretraining and gain insights into what the PLM has learned, we examined attention maps and residue representations for the GB1 METL-Local model after pretraining on molecular simulations but before finetuning on experimental data (Fig. 5). Our METL PLMs with 3D relative position embeddings start with a strong inductive bias and include the wild-type protein structure as input. After pretraining, the METL attention map for the wild-type GB1 sequence closely resembles the residue distance matrix of the wild-type GB1 structure (Fig. 5ab). In contrast, an alternative METL model with 1D relative position embeddings that does not use the GB1 structure while training fails to learn an attention map that resembles the GB1 contacts (Fig. 5c). The 3D relative position embedding and pretraining successfully allows METL to focus attention on residue pairs that are close in 3D space and may be functionally important.

**Figure 5.**
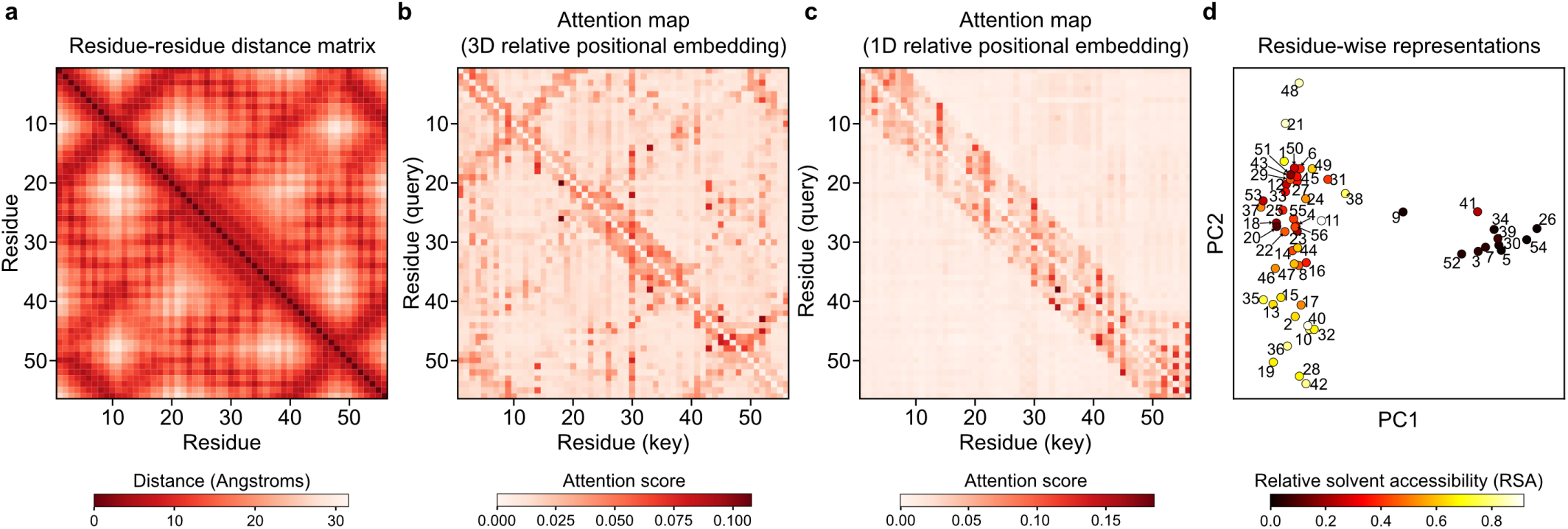
METL attention maps and residue representations relate to structure and biophysical properties. **(a)** The residue distance matrix shows C*β* distances between residues for the wild-type GB1 structure. **(b-c)** The attention maps show the mean attention across layers and attention heads for the wild-type GB1 sequence when it is fed as input to the pretrained GB1 METL-Local model. The 3D structure-based relative position embeddings (RPEs) enable the network to focus attention on residues that are close in 3D space, effectively capturing GB1’s structural contacts. The 1D sequence-based RPEs do not. **(d)** Principal component analysis (PCA) of the residue representations output by the pretrained GB1 METL-Local model, averaged across the 20 possible amino acids at each sequence position. Points are colored according to relative solvent accessibility (RSA) computed from the wild-type GB1 structure.

We further explored the information encoded in the pretrained GB1 METL model by visualizing residue-level representations at each sequence position, averaged across amino acid types (Fig. 5d). These residue-level representations show strong clustering based on a residue’s relative solvent accessibility (RSA) and weaker organization based on a residue’s location in the three-dimensional structure, as observed through visual inspection and qualitative cross-checking with residue–residue distance patterns. Analysis of the additional datasets in our study reaffirmed these findings: models with 3D relative position embeddings consistently focused attention on spatially proximate residues, and residue representations showed RSA-based clustering patterns across all datasets (Figs. S15 and S16). This suggests the pretrained METL models have an underlying understanding of protein structure and important factors like residue burial, even before they have seen any experimental data.

To test whether METL pretraining learns underlying epistatic interactions, we evaluated GB1 variants with well-characterized epistatic effects [26]. The pretrained METL-Local model successfully identifies known interacting positions in GB1’s dynamic *β* 1-*β* 2 loop region, with pairwise combinations of positions 7, 9, and 11 all ranking in the top 10% of predicted positional epistasis. The pretrained model also captures strong negative epistasis in the G41L,V54G double mutant (top 0.5% of predicted epistasis), consistent with the known compensatory exchange of small-to-large and large-to-small residues. However, METL underestimates the disulfide-driven positive epistasis in the Y3C,A26C variant, likely due to Rosetta’s lack of automatic disulfide bond modeling while generating pretraining data. Overall, these findings demonstrate that METL’s pretrained representations capture biologically-relevant structural information driving epistasis, while also highlighting a potential limitation of Rosetta-based pretraining.

### Function-specific simulations improve METL representations

METL models are pretrained on general structural and biophysical attributes but are not tailored to any particular protein property such as ligand binding, enzyme activity, or fluorescence. There is a great body of research using molecular simulations to model protein conformational dynamics, small molecule ligand and protein docking, enzyme transition state stabilization, and other function-specific characteristics [27–31]. These function-specific simulations can be used to generate METL pretraining data that is more closely aligned with target functions and experimental measurements. Similarity between pretraining and target tasks is important to achieve strong performance and avoid detrimental effects in transfer learning [32].

To demonstrate how function-specific simulations can improve the initial pretrained METL model and its performance after finetuning, we customized the GB1 simulations to more closely match the experimental conditions. The GB1 experimental data measured the binding interaction between GB1 variants and Immunoglobulin G (IgG) [26]. To match this experimentally characterized function, we expanded our Rosetta pipeline to model the GB1-IgG complex and compute 17 attributes related to energy changes upon binding (Table S5). These function-specific attributes are more correlated with the experimental data than the general biophysical attributes (Fig. S17), suggesting they could provide a valuable signal for model pretraining.

We pretrained a METL PLM that incorporates the IgG binding attributes into its pretraining data and refer to it as METL-Bind (Fig. 6a). METL-Bind is a variant of METL-Local and is specific to GB1. METL-Bind outperformed a standard METL-Local PLM, pretrained only with GB1 biophysical attributes, when finetuned on limited experimental data (Fig. 6b-c and Fig. S18). We calculated the predictive error for each residue position in the GB1 sequence to understand if the two models specialize on distinct structural regions (Fig. 6d-e). METL-Bind performed better across most residue positions and was notably better at predicting mutation effects at the GB1-IgG interface. The residue where METL-Bind showed the largest improvement was glutamate 27, an interface residue vital for the formation of a stable GB1-IgG complex [33].

**Figure 6.**
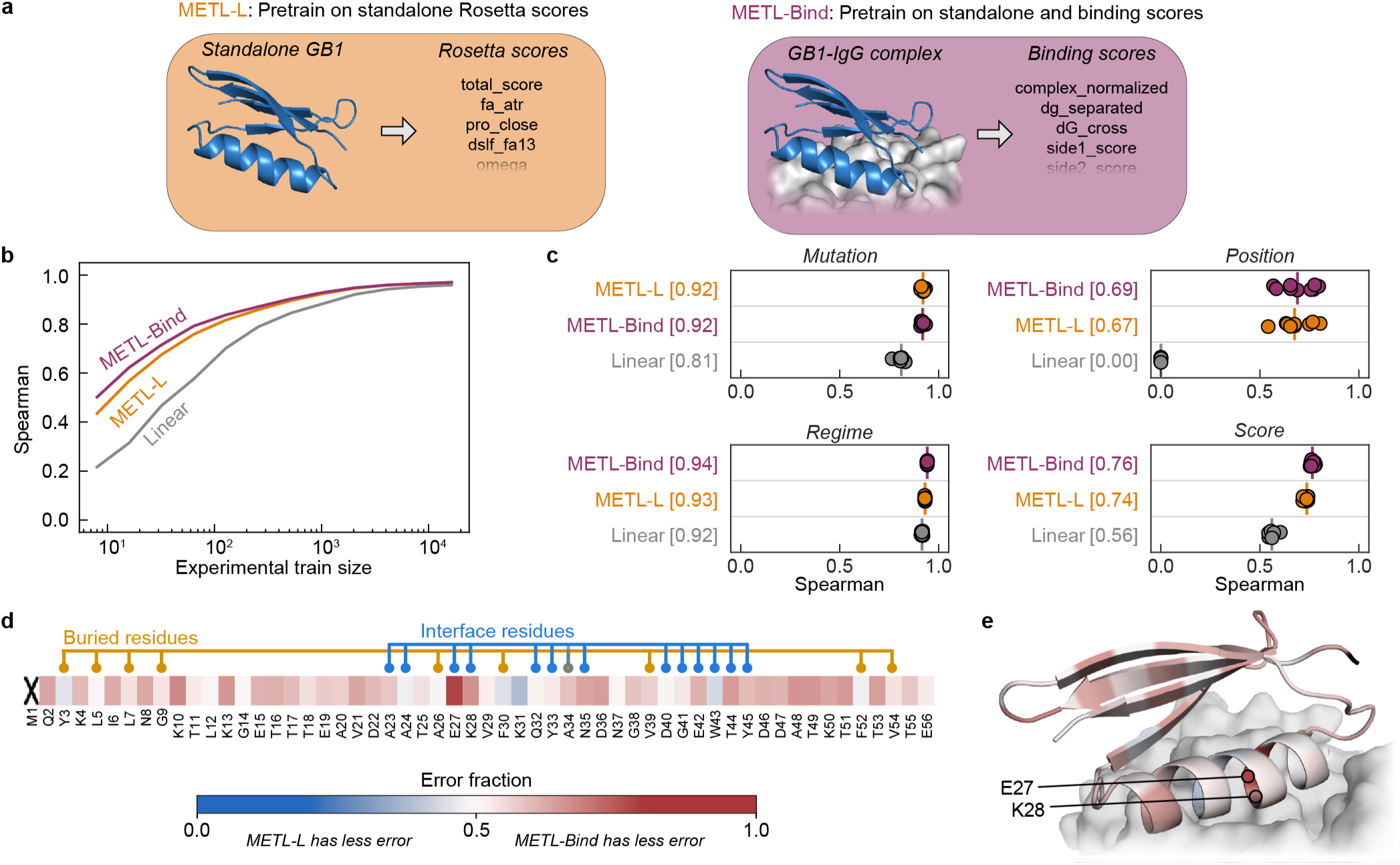
Function-specific simulations improve METL pretraining for GB1. **(a)** METL-Local (METL-L) pretrains on general Rosetta biophysical attributes from the standalone GB1 structure. METL-Bind pretrains on both general biophysical attributes from the standalone GB1 structure and binding-specific scores from the GB1-IgG complex structure. **(b-c)** Learning curves and extrapolation performance for Linear, METL-Local, and METL-Bind on the GB1 dataset. We pretrained METL-Local and METL-Bind on the same variants, differing only in the Rosetta score terms. We used the same finetuning dataset splits and replicates as in Figure 2. Each point represents 1 of 9 extrapolation replicates. The vertical bars denote the medians of the replicates, and the square brackets indicate the median Spearman correlations. **(d-e)** The heatmap shows the fraction of test set variants for which METL-Bind has lower error than METL-Local, broken down by sequence position. Results are shown for training set size 32 and averaged across replicates. Position 1 is marked with an “X” because the dataset does not contain variants with mutations in that position. METL-Bind has less error for 44 out of 55 sequence positions. The structure shows the GB1-IgG interface with the GB1 structure colored using same error fraction as the heatmap.

While both models converge to similar performance with abundant training data, METL-Bind’s superior performance with limited data shows that pretraining on the additional GB1-IgG complex attributes successfully improved the model’s learned representation. Many important protein properties can only be assayed accurately using low-throughput techniques. METL-Bind is a promising proof of concept for enhancing predictions when those properties can be approximated computationally.

Pretraining on function-specific simulations provides METL with an initial awareness of protein function that can be integrated with limited experimental data.

### METL generalization to design diverse GFP variants

Predictive models can guide searches over the sequence-function landscape to enhance natural proteins or design new proteins [6, 34, 35]. However, these models often face the challenge of making predictions based on limited training data or extrapolating to unexplored regions of sequence space. To demonstrate METL’s potential for real protein engineering applications, we tested METL-Local’s ability to prioritize fluorescent GFP variants in these challenging design scenarios. We used METL-Local to design 20 GFP sequences that were not part of the original dataset, and we experimentally validated the resulting variants to measure their fluorescence brightness (Fig. 7).

**Figure 7.**
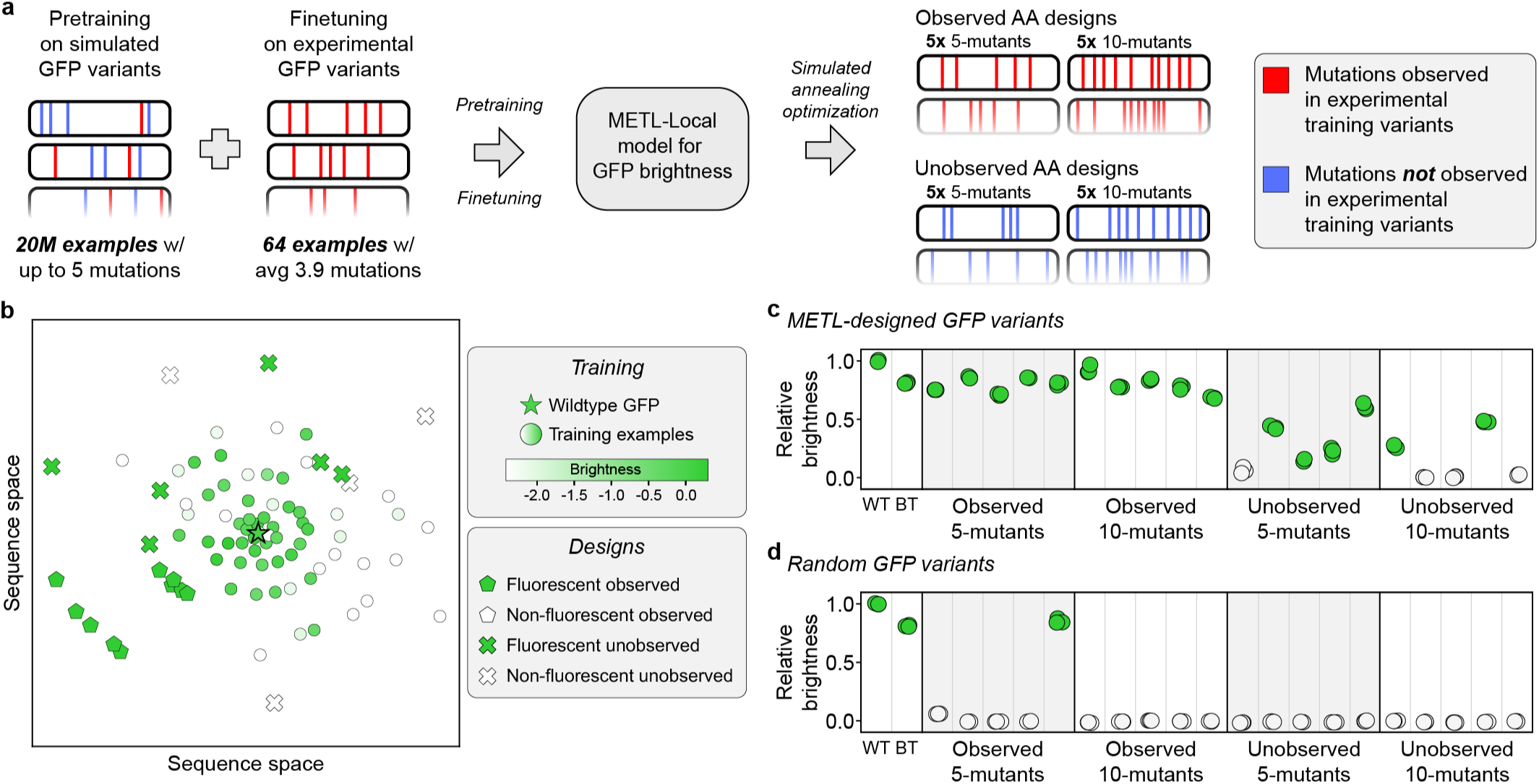
Low-N GFP Design. **(a)** Overview of the GFP design experiment. We used METL-Local to guide GFP design in a low-N setting with only *N* = 64 experimental training examples. We tested two different design constraints: *Observed AA*, where sequences contain only amino acid substitutions observed in the training set, and *Unobserved AA*, where sequences exclude any amino acid substitutions observed in the training set. **(b)** Multidimensional scaling (MDS) sequence space visualization of the wild-type GFP sequence, the 64 GFP training sequences, and the 20 designed proteins. The designed sequences contain either 5 or 10 amino acid substitutions from wild-type. Training set sequences are colored on a gradient according to their experimental brightness score. Designed sequences are colored according to whether they exhibited fluorescence, which we define as having at least 10% of wild-type GFP’s brightness. **(c)** Experimentally characterized brightness of the designed sequences, the best training set sequence (BT), and the wild-type sequence (WT). Each dot represents one distinct sample of the three replicates. **(d)** Experimentally characterized brightness of the random baselines. Evolutionary data, consisting of massive collections of naturally evolved protein sequences, captures information relevant to organismal fitness, including protein expression, folding, stability, and biological function. However, the precise selective pressures for each protein are different and largely unknown, and evolutionary patterns can be confounded by historical events, phylogenetic biases, and unequal sequence sampling [46]. In contrast, biophysical simulations allow precise control of the input sequence distribution, even sequences with non-natural amino acids [47, 48], and capture fundamental properties of protein structure and energetics. Yet, biophysical simulations are only imperfect approximations of the true physics.

We intentionally set up the design tasks to mimic real protein engineering settings with limited data and extrapolation. We finetuned a METL-Local PLM on only 64 GFP variants randomly sampled from the full dataset. The 64 sampled variants had an average of 3.9 amino acid substitutions and a fitness distribution similar to the full dataset (Figs. S19 and S20). We designed variants with either 5 or 10 amino acid substitutions, forcing the model to perform regime extrapolation. Furthermore, we tested two design scenarios, *Observed AA* and *Unobserved AA*, in which designed variants were constrained to either include or exclude amino acid substitutions observed in the training set, respectively. The Unobserved AA setting forces the model to perform mutation and/or position extrapolation. We designed five variants at each extrapolation distance (5- and 10-mutants) and design setting (Observed AA and Unobserved AA) (Fig. S21 and Table S6). We used simulated annealing to search sequence space for GFP designs that maximize METL-Local’s predicted fitness and clustered the designs to select diverse sequences. We also sampled random variants under the same scenarios as the METL designs to serve as baselines.

We had the genes for the 20 GFP METL designs and the 20 random baselines synthesized and cloned into an expression vector as a fusion protein with the fluorescent protein mKate2, emulating the conditions used to collect the training data [36]. The mKate2 is constant in each fusion protein, while the GFP sequence varies. The ratio of a GFP variant’s fluorescence to mKate2’s fluorescence provides an intrinsic measure of the GFP variant’s “relative brightness” that is independent of the absolute protein expression level [37]. Overall, METL was successful at designing functional GFP variants, with 16 of the 20 designs exhibiting measurable fluorescence (Fig. 7c). Each design setting had notable differences in the success rates and fluorescence characteristics of the designed GFP sequences. The Observed design setting was 100% successful at designing fluorescent five (5/5) and ten (5/5) mutants, demonstrating METL’s robust ability to learn from very limited data and extrapolate to higher mutational combinations. The more challenging Unobserved design setting had an 80% (4/5) hit rate with five mutants and a 40% (2/5) hit rate with ten mutants. The Unobserved designs were less bright than wild-type GFP and the Observed designs.

The random baselines provide context for evaluating the designed variants and METL-Local’s predictions (Fig. 7d). Across all design scenarios, the random baseline variants exhibited minimal or no fluorescence activity, with the exception of one of the Observed 5-mutant baselines, which fluoresced. METL-Local assigns a high predicted score to this variant, showing its ability to recognize functional sequences (Fig. S22). Conversely, METL-Local did not predict high scores for any of the other random baselines. This suggests that the functional METL-designed variants likely emerged from the model’s understanding of the GFP fluorescence landscape rather than random chance.

The mKate2 fluorescence signal provides additional insight into the designs (Fig. S23). The mKate2 protein is constant, so changes in its fluorescence signal are caused by changes in mKate2-GFP fusion protein concentration and thus provide an indirect readout of the GFP designs’ folding, stability, solubility, and aggregation. The Observed designs all exhibit higher mKate2 fluorescence than wild-type GFP, possibly indicating moderate stabilization, while the Unobserved designs mostly exhibit lower mKate2 fluorescence than wild-type GFP, suggesting destabilization.

### Accessing METL tools

In addition to making the METL code, models, and datasets available (Methods), we also made them accessible through multiple web interfaces. We provide a Hugging Face interface to download and use our METL models (https://huggingface.co/gitter-lab/METL) and a Hugging Face Spaces demo (https://huggingface.co/spaces/gitter-lab/METL_demo) [38]. The Gradio [39] web demo supports generating predictions with our pretrained METL models for a list of sequence variants and visualizes those variants on the protein structure [40]. We created two Colab notebooks to run METL workflows with GPU support, which are available from https://github.com/gitter-lab/metl. One notebook is for loading a pretrained METL model and finetuning it with user-specified protein sequence-function data. The other is for making predictions with pretrained METL models, the same functionality as the Hugging Face Spaces demo but better-suited for large datasets. These Colab notebooks are part of the Open Protein Modeling Consortium [41]. Finally, the METL GitHub repository also links to a Jupyter notebook to generate Rosetta pretraining data at scale in the Open Science Pool [42] for eligible researchers.

## Discussion

Motivated by decades of research into biophysics, molecular dynamics, and protein simulation [11, 27, 28, 31, 43], we present METL, which leverages synthetic data from molecular simulations to pretrain biophysics-aware PLMs. These biophysical pretraining signals are in contrast to existing PLMs or multiple sequence alignment-based methods that train on natural sequences and capture signals related to evolutionary selective pressures [2, 7, 8, 15, 44, 45]. By pretraining on large-scale molecular simulations, METL learns a biophysically-informed representation of protein space, which provides valuable context for understanding protein sequence-function relationships. Pretrained METL models can be finetuned on experimental data to produce models that integrate biophysical knowledge and are capable of predicting properties such as binding, thermostability, and expression. METL excels at challenging protein engineering tasks such as learning from limited data and extrapolating to mutations not observed in the training data, enabling the design of new proteins with desired properties.

Our results highlight important differences between evolutionary data and biophysical simulations, especially in terms of their effectiveness for pretraining PLMs to understand sequence-function relationships and predict experimental functions.

Generally, we found that the protein-specific models METL-Local, Linear-EVE, and ProteinNPT demonstrated superior performance compared to general protein representation models METL-Global and ESM-2. The relative performance of METL-Local and Linear-EVE was partly determined by a dataset’s correlation with Rosetta *total score* and EVE, respectively. Certain protein properties and experimental measurements more closely align with either biophysical or evolutionary signals [49–51], providing guidance on where different models may excel. One of METL’s key strengths is its ability to incorporate function-specific molecular modeling and simulations. For instance, pretraining on GB1-IgG binding data led to improved performance compared to our standard METL-Local model, which was pretrained only on GB1 structure-derived data. This opens the door to incorporating more sophisticated simulations, such as dynamic simulations of conformational transitions in allosteric regulation, quantum mechanics/molecular mechanics (QM/MM) studies of enzyme catalysis, coarse grained models of macromolecular machines, and small molecule docking to assess binding specificity. It would also be straightforward to extend METL to make multitask predictions, such as both GB1 thermostability and GB1-IgG binding affinity.

The current version of METL-Global represents an initial step toward a universal biophysics-based representation of all proteins. METL-Global provides a comparable or better representation than a similarly-sized ESM-2 model when finetuning with small training sets for all datasets except GRB2-A (Fig. 2). However, METL-Global overfits to the proteins it was pretrained on (Fig. S1), indicating there is room for improvement. We can greatly expand the number and diversity of protein structures used for pretraining METL-Global using the RCSB Protein Data Bank (PDB) [52] or AlphaFold Protein Structure Database [53]. Meta-learning strategies [54, 55] could alleviate overfitting to the pretraining structures and have helped with domain generalization in chemical screening [56]. In this study, we intentionally examined biophysical and evolutionary signals separately by training METL models from scratch and comparing them to evolutionary-based models. Future iterations of METL-Global could integrate these signals by leveraging evolutionary PLMs as a pretrained foundation, potentially enhancing generalization by combining complementary information from both domains. Sequence-based PLMs can learn about protein structure from evolutionary statistics [19, 57–59]. However, many recent PLMs directly incorporate structural information [60–64], and we envision METL-Global would continue to use this prior knowledge when it is available.

Prior studies have integrated biophysics and machine learning either by using biophysics-based features as input to machine learning models or approximating biophysical simulations with machine learning. Rosetta and FoldX stability, energy, and docking terms have been provided as features for an augmented linear regression model [17], random forests [65, 66], a 2D CNN [67], and on nodes and edges in a graph neural network [68] for antibody and protein property prediction. Function-value-prior augmented-Bayesian Neural Networks can incorporate Rosetta stability as a prior on protein function prediction in regions where a Bayesian Neural Network has high epistemic uncertainty [69]. Molecular dynamics-derived features have been included in supervised learning models of big potassium channels [70] and bovine enterokinase [71]. Wittmann et al. evaluate Triad ΔΔG predictions for selecting initial variants for machine learning-guided directed evolution [72]. Unlike a finetuned METL-Local model, all of these approaches must run the biophysics calculations for each sequence prediction, which could limit their scalability to search sequence space for protein design. Other related work uses machine learning to approximate molecular simulations [73], often with the goal of obtaining much faster approximate models [74]. This scenario is similar to METL’s pretraining stage. These methods include the Epistasis Neural Network that has been used to engineer xylanases [75] and GFP variants [76], molecular dynamics approximations to minimize energy and match a target structure [77], learning to predict Rosetta protein-ligand binding energy to speed up variant scoring [78], and sampling protein conformations [79]. ForceGen trains a protein language diffusion model on molecular dynamics simulations of mechanical unfolding responses [80]. METL’s pretraining on biophysical attributes for protein engineering is also related to the long-standing problem of predicting protein stability [81–92]. Finally, machine learning has been integrated into Rosetta to guide its sampling [93].

Machine learning-guided protein engineering is often data-limited due to experimental throughput constraints, with datasets sometimes containing as few as tens to hundreds of sequence-function examples [34, 94–99]. We demonstrated METL’s performance in realistic protein engineering settings with limited data (low-N) and extrapolation. PLMs are an important component in many existing methods for low-N protein engineering. They have been used to extract protein sequence representations [3, 100–103], for finetuning on the low-N function data [102, 104–106], to predict structures and derive features [107], and to generate auxiliary training data in more complex models [18, 105, 108]. Other computational strategies for addressing the low-N problem include Gaussian processes [101, 109, 110], augmenting regression models with sequence-based [17, 111] or structure-based [112] scores, custom protein representations that can produce pretraining data [113], representations of proteins’ 3D shape [114], active learning [115], few-shot learning [116], meta learning [117, 118], contrastive finetuning [119], and causal inference [120].

Our GFP design experiments showcased METL’s ability to learn from only 64 training examples and generalize to distant and unexplored regions of sequence space. METL’s success in the Unobserved AA design setting was especially remarkable because it requires the model to infer the effects of mutations it has not observed and predict how these mutations combine in 5- and 10-mutants. It is notable that none of the designed GFPs appeared brighter than wild-type GFP. We estimated brightness as the ratio of GFP fluorescence to mKate2 fluorescence. We noticed many of the designed variants exhibited absolute GFP and mKate2 fluorescence signals higher than wildtype, indicating that the mKate2-GFP fusion protein may have increased expression levels and improved stability in these variants. In limited data settings, METL-Local’s strong biophysical prior may indirectly improve designs through stabilizing effects rather than directly improving the brightness.

Examples across diverse scientific domains have demonstrated the power of combining simulations and machine learning [121], spanning topics such as gene regulatory network reconstruction [122], chemical foundation model pretraining [123], climate emulation [124], and quantum chemistry approximation [31, 125]. METL fits within this broader trend and represents a significant step toward effectively integrating biophysics insights with machine learning-based protein fitness prediction. The METL framework pretrains PLMs on molecular simulations to capture accumulated biophysical knowledge, and this pretraining strategy will benefit from continued advances in computation and molecular simulation. METL can pretrain on general structural and energetic terms or more focused function-specific terms, offering the potential to model completely non-natural protein functions with nonexistent evolutionary signals. PLMs fluent in fundamental biophysical dialect will push the boundaries of protein design to new realms of sequence-function space.

## Methods

### Generating Rosetta pretraining data

The Rosetta pretraining data consists of protein sequences and their corresponding score terms, computed by modeling the sequences with Rosetta. We refer to the METL models pretrained on the Rosetta biophysical attributes as source models. The data used for local and global source models differs in what sequences are included. Rosetta data for local source models contains protein variants within the local sequence space surrounding the protein of interest. Rosetta data for global source models contains protein variants from a diverse range of base sequences and structures.

We generated local Rosetta datasets for each target protein from the experimental datasets (Table S3). We acquired the target protein structures from RCSB PDB [52] and AlphaFold Protein Structure Database [53]. For cases where the acquired structure did not match the reference sequence of the target protein, we used Rosetta comparative modeling or truncated the acquired structure to match the reference sequence. For each local pretraining dataset, we generated ∼20M protein sequence variants with a maximum of 5 amino acid substitutions. See Table S7 for additional details regarding local Rosetta dataset structures and variants, including exceptions to the above.

We generated the global Rosetta dataset based on 150 diverse protein structures identified in Supplementary Table 1 of Kosciolek and Jones [14]. We downloaded the 150 structures from RCSB PDB [52]. Some structures contained modified or missing residues. We replaced modified residues with canonical amino acids and used the RosettaRemodel application to fill in the structure of missing residues. We were unable to remodel PDB IDs 1J3A and 1JBE, thus we excluded these structures from the final dataset. For each of the remaining 148 structures (Table S2), we generated ∼200K variants with a maximum of 5 amino acid substitutions, for a total of ∼30M variants.

To assess the similarity between the 148 proteins used for pretraining METL-Global and the 8 proteins used for evaluation, we clustered all proteins based on sequence and structure representations. We ran sequence-based clustering with MMseqs2 v15.6f452 [126] and structure-based clustering with Foldseek v9.427df8a [127]. We set a coverage threshold 0.5 for both sequence- and structure-based clustering and identified clusters that contained structures from the METL-Global pretraining collection and one of the 8 unique structures used for evaluation. At this threshold, both MMseqs2 and Foldseek identified structures in the METL-Global pretraining collection that are similar to the GRB2 and TEM-1 structures. Foldseek also detected a match for DLG4 and a second matching structure for GRB2. We aligned the pairs of co-clustered structures with the RCSB PDB pairwise structure alignment tool [128] and the TM-align alignment method [129].

We implemented a custom sub-variants sampling algorithm to generate the variants for both the local and global datasets. The algorithm iteratively samples a random variant with 5 amino acid substitutions from the wild-type sequence then generates all possible 4-, 3-, 2- and 1-substitution sub-variants with the same amino acid substitutions as the 5-substitution variant. Duplicate variants generated through this process are discarded. The iterations terminate when the target number of variants is reached. For the global dataset, we used the sub-variants sampling algorithm to generate all of the ∼200K variants per base sequence. For the local datasets, we first generated all possible 1-substitution or 1- and 2-substitution variants, and then we used the sub-variants sampling algorithm to generate the remainder of the ∼20M variants per target protein (Table S7).

Once sequence variants were generated, we used Rosetta to compute biophysical attributes for each variant sequence. We first prepared each base PDB file for use with Rosetta by following the recommendation in the Rosetta documentation. We ran Rosetta’s `clean_pdb.py` and relaxed the structure with all-heavy-atom constraints. We generated 10 structures and selected the lowest energy structure to serve as the base structure for subsequent steps.

We used Rosetta v3.13 [11] to compute full-atom energy terms (ref2015 score function), centroid-atom energy terms (score3 score function), and custom filter terms based on Rocklin et al. [130]. For each variant, we introduced the variant’s mutations to the corresponding base structure using a Rosetta resfile. Then, to generate the full-atom energy terms, we used FastRelax to relax the mutated structure using the ref2015 score function, only repacking residues within 10Å of the mutated residues, with 1 repeat. To generate the centroid-atom energy terms, we used score_jd2 to score the resulting structure using the score3 score function. Finally, to generate the remainder of the score terms used in the standard version of METL, we used a RosettaScript to compute custom filter terms on the relaxed structure. To calculate additional binding scores for METL-Bind, we used the Rosetta InterfaceAnalyzer protocol. See Tables S1 and S5 for a list and description of each term. We designed a computing workflow based on HTCondor [131] to orchestrate the Rosetta scoring on the Open Science Pool [42]. Rosetta simulation times scale with protein sequence length. Average runtimes per variant ranged from ∼37-50 seconds for smaller proteins (56-102 residues, e.g., GB1, GRB2, Pab1, Ube4b) to 135-215 seconds for larger proteins (237-403 residues, e.g., GFP, PTEN, TEM-1).

### Preprocessing Rosetta pretraining data

Prior to training neural networks, we preprocessed the raw Rosetta data by dropping variants with NaN values for any of the biophysical attributes (343 in the global dataset and 645 in the GB1 dataset, corresponding to 0.0012% and 0.0051% of each dataset, respectively). No variants with NaN values were present in the other datasets. We also removed duplicates by randomly selecting one of the duplicates to keep and filtered out variants with outlier *total*_*score* values. We grouped variants by base PDB and removed outliers independently for each group using a modified z-score method, which uses the median and median absolute deviation instead of the mean and standard deviation. For each data point *i*, we calculated the modified z-score using the following equation:

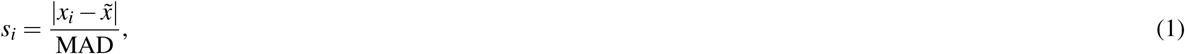

where *s_i_* is the modified z-score, *x_i_* is the Rosetta *total*_*score*, 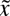 is the median *total*_*score* of the group, and MAD is the Median Absolute Deviation, defined as 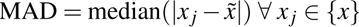, or the median of the absolute deviations of all data points from the median of the group. We removed variants with *s_i_ >* 6.5 from the dataset.

Additionally, we standardized the Rosetta scores to equalize the contribution of each score term to the model’s loss function and to ensure score terms are comparable across different base PDBs in the global dataset. Once again, we grouped variants by base PDB, and then we standardized each group and score term independently by subtracting the mean and dividing by the standard deviation. We calculated the mean and standard deviation using only the training set data. This process scales the score terms to have zero mean and a standard deviation of one.

We excluded the following score terms from the final dataset because the values were zero for a large portion of base PDBs: dslf_fa13 (from ref2015 score function), linear_chainbreak and overlap_chainbreak (from score3 score function), and filter_total_score (custom filter term). We also discarded res_count_all (custom filter term that counts the residues in the protein) because it did not vary among variants of an individual protein. After these removals, 55 score terms remained (Table S1).

### METL source model architecture

The METL source model architecture accepts amino acid sequences as input and outputs predictions for each of the 55 Rosetta score terms. The main component of the source model architecture is a transformer encoder based on the original transformer architecture [12], with the notable differences being the use of a relative positional embedding [13] instead of a sinusoidal positional encoding and pre-layer normalization instead of post-layer normalization [132]. METL-Local source models total ∼2.5M parameters and have transformer encoders consisting of a 256 embedding size, 3 encoder layers, 4 attention heads, a 1024 feed forward hidden size, and 0.1 dropout. METL-Global source models total ∼20M parameters and have transformer encoders consisting of a 512 embedding size, 6 encoder layers, 8 attention heads, a 2048 feed forward hidden size, and 0.1 dropout. We also evaluated a METL-Global source model with ∼50M parameters, consisting of a similar architecture as the 20M parameter METL-Global source model but with 16 encoder layers instead of 6 encoder layers. After the transformer encoder, source models implement an additional layer normalization layer, a global average pooling layer, a nonlinear fully-connected layer, and a linear output layer with 55 output nodes corresponding to the 55 Rosetta score terms. The global average pooling layer computes the mean of the per-residue encodings, which are output from the encoder, to produce a sequence-level representation of the same size as the embedding dimension. This sequence-level encoding is fed into a fully-connected layer with 256 hidden nodes for the local model and 512 hidden nodes for the global model. We used the rectified linear unit (ReLU) activation function for the transformer encoder and final fully-connected layer.

We implemented relative position embeddings as described by Shaw et al. [13]. In contrast to the absolute position encoding used in the original transformer architecture [12], the relative position embedding enables the network to consider positional representations of the inputs in terms of distances between sequence positions. We consider two distinct ways to encode relative distances, generating what we refer to as 1D positional embeddings and 3D positional embeddings. In the 1D approach, relative distances are based on the protein amino acid sequence alone. This approach is identical to the implementation of relative position embeddings described by Shaw et al. In the 3D approach, relative distances are based on the 3D protein structure.

In the 1D approach, we calculate relative distances by determining the offset between each pair of sequence positions (*i*, *j*) in the input. The relative distance is defined as *d* = *j* − *i*, representing how far sequence position *j* is relative to position *i*. A negative value signifies that *j* precedes *i* in the sequence, and a positive value signifies that *j* succeeds *i*. We map each of the possible relative distances to a pair of learnable embedding vectors, corresponding to attention keys and values. When calculating attention between sequence positions *i* and *j*, we add the key and value positional embedding vectors to the keys and values, respectively. As was hypothesized by Shaw et al., precise relative position information might not be useful beyond a certain distance. Thus, we clipped the possible relative distances to ±8.

In the 3D approach, we calculate relative distances using the protein 3D structure instead of the amino acid sequence. When using 3D relative position embeddings, the model requires a protein structure in the form of a PDB file, corresponding to the base protein that the input variant sequence is based on. We first represent the protein structure as an undirected graph, where each node corresponds to a residue. We place an edge between any pair of nodes if the beta carbon atoms (C*β*) of the residues are within 8Å of each other in the 3D space. We define the relative distance between residues (*i*, *j*) as the minimum path length from node *i* to node *j* in the graph. Unlike the 1D approach, relative distances computed using the 3D approach cannot be negative values. We clip the 3D relative distances at 3, effectively transforming distances greater than 3 into a relative distance of 3. A relative distance of 0 represents a node with itself, 1 signifies direct neighbors, 2 signifies second degree neighbors, and 3 encapsulates any other node not covered by the previous categories. As in the 1D approach, each possible relative distance in the 3D approach is mapped to a pair of embedding vectors corresponding to keys and values. These vectors are learned during training and are added to keys and values during the attention calculation.

### METL source model training

We split the Rosetta source data into randomly sampled train, validation, test, and withheld sets. For each dataset, we first withheld 5% of the data to be used for final evaluations. We split the remaining data into 80% train, 10% validation, and 10% test sets.

We trained source models for 30 epochs using the AdamW optimizer [133] with a learning rate of 0.001. We applied a linear warm-up learning rate scheduler, with a warm-up period of 2% of the total training steps. Additional AdamW hyperparameters were *weight*_*decay* = 0.01, *β*_1_ = 0.9, *β*_2_ = 0.999, and *ε* = 1*e* − 8. We computed mean squared error loss independently for each of the 55 prediction tasks (corresponding to the 55 Rosetta biophysical attributes) and took the sum to compute the final loss for the network. We applied gradient norm clipping with a max norm of 0.5. We employed distributed data parallel (DDP) training with 4 GPUs using PyTorch Lightning [134, 135]. We trained local source models with an effective batch size of 2048 (512 x 4 GPUs) and global source models with an effective batch size of 1024 (256 x 4 GPUs). For the METL-Bind experiment, we trained both standard METL-Local and METL-Bind using the same process, except using 2 GPUs instead of 4 and a batch size of 1024 instead of 512, which yielded an effective batch size 2048, identical to the source models trained for the main experiment. METL-Bind was trained on 17 additional binding scores, for a total of 55 + 17 = 72 tasks, but was otherwise identical to the standard METL-Local model.

The global source data contains variants of 148 base sequences, with most having different sequence lengths. This complicates the process of encoding data into a single fixed-length batch. Padding is a commonly employed approach in such scenarios. However, incorporating different sequence lengths and base structures in a single batch would negatively impact efficiency of computing attention with our implementation of relative position embeddings. Thus, we implemented a PDB-based data sampler that ensures each batch only contains variants from a single base PDB structure. Due to the use of DDP training with 4 GPUs, each aggregated training batch effectively contains variants from 4 base PDBs.

Pretraining times depend on the model size, protein size, amount of simulated data, and computational resources. For a 2M parameter METL-Local model with a simulated data size of ∼20M examples, running on 4x NVIDIA A100 or A30 GPUs, pretraining times ranged from 6-13 hours for 30 epochs (12 to 26 minutes per epoch) for most proteins, with the exception of PTEN, which took ∼33 hours for 30 epochs (1.1 hours per epoch). With a smaller training size of ∼1M examples and just a single GPU, training times ranged from 6-26 hours for 100 epochs for most proteins (4 to 16 minutes per epoch). Pretraining METL-Global with 20M parameters took ∼50 hours on 4x A100s and ∼142 hours with 50M parameters.

### Experimental datasets for target model training

The METL target model architecture accepts amino acid sequences as input and outputs predictions for one specific protein function. We evaluated METL on experimental datasets representing proteins of varying sizes, folds, and functions: GFP [36], DLG4-2021 [136], DLG4-Abundance [137], DLG4-Binding [137], GB1 [26], GRB2-Abundance [137], GRB2-Binding [137], Pab1 [138], PTEN-Abundance [139], PTEN-Activity [140], TEM-1 [141], and Ube4b [142] (Table S3). We acquired raw datasets from published manuscript supplements, MaveDB [143], and NCBI GEO [144]. We transformed raw data into a standardized format, making sure that functional scores were log-transformed, normalized so that the wild-type score is 0, and rounded to 7 decimal places. We removed variants with mutations to stop codons and converted variant indexing to be 0-based. For DLG4-2021 and GB1, we filtered variants to ensure a minimum number of reads. See Table S8 for additional details about dataset transformations. We opted to use the DLG4 dataset instead of the DLG4-2021 dataset in our main analysis due to weak correlation between the two datasets (Fig. S24) and because linear regression yielded better results on the DLG4 dataset, suggesting a cleaner signal.

We used GB1 as an exploratory dataset during method development to make modeling decisions such as at what size validation set to enable model selection, where to place the prediction head on the source model, whether to use a linear or nonlinear prediction head, and others. Due to this, there is potential we overfit to GB1 and that our final results are optimistic for GB1. That said, we took precautions to limit the potential impact of using GB1 as our development dataset. The results presented for the small training set size experiment use an evaluation dataset that was completely held out, even during method development. The randomly sampled train and validation sets used to generate the final results are also different splits than the ones we used during method development. Additionally, the results presented for the extrapolation experiments use different splits than the ones we used to test extrapolation during method development.

We adjusted the GFP dataset preprocessing after seeing early small training set size results. Performance was lower than expected, which led us to realize that the dataset scores were not normalized so wild-type is 0. We modified the GFP dataset to normalize the scores and set wild-type to 0 by subtracting the wild-type score from all the scores. All our other datasets were already normalized so wild-type is 0.

### METL target model architecture

METL target models are made up of a backbone and a head. The backbone contains network layers from the METL source model, pretrained to predict Rosetta biophysical attributes. The head is a new, randomly-initialized linear layer placed on top of the backbone to predict experimental functional scores. We also added a dropout layer with dropout rate 0.5 between the backbone and the head. For METL-Local source models, we attach the head immediately after the final fully-connected layer. For METL-Global source models, we attach the head immediately after the global pooling layer. METL target models have a single output node corresponding to the experimental functional score prediction.

### METL target model training

We implemented two training strategies for PLM target models: feature extraction and finetuning. Feature extraction is a training strategy where only the head is trained, and the backbone weights are not updated during the training process. In contrast, finetuning is a training strategy where both the backbone and head weights are updated during training. For feature extraction, we trained the head using scikit-learn [145] ridge regression with *alpha* = 1.0 and the cholesky solver. This provides a closed-form solution for the ridge regression weights.

For finetuning, we implemented a dual-phase finetuning strategy [146]. In the first phase, we froze the backbone and trained only the head for 250 epochs. In the second phase, we trained both the backbone and the head for an additional 250 epochs at a reduced learning rate. We used the AdamW optimizer with a learning rate of 0.001 in the first phase and 0.0001 in the second phase. We applied a learning rate scheduler with linear warm-up and cosine decay to each phase, with a warm-up period of 1% of the total training steps. Additional AdamW hyperparameters were set as follows: *weight*_*decay* = 0.1, *β*_1_ = 0.9, *β*_2_ = 0.999, and *ε* = 1*e* − 8. We used a batch size of 128 and mean squared error loss. We applied gradient norm clipping with a max norm of 0.5.

After the full training period, we selected the model from the epoch with the lowest validation set loss. We only performed model selection if the validation set size was ≥ 32 for METL-Local and ≥ 128 for METL-Global and ESM-2. We found the optimization was more stable for METL-Local than METL-Global and ESM-2, thus smaller validation sets were still reliable. For validation sets smaller than those thresholds, we did not perform model selection. Instead, we used the model from the last epoch of training. We determined these thresholds using the GB1 dataset, which we designated as our development dataset, by selecting the dataset size along the learning curve where using model selection started to outperform not using model selection. In retrospect, these thresholds were too low for other datasets, leading to the dips in METL-Global correlations observed in Figure 2.

Finetuning METL-Local was relatively quick, with training times scaling with the experimental dataset size. For a dataset size of 320 examples, finetuning typically took ∼2-5 minutes; for 20,480 examples, it took ∼20-42 minutes. Finetuning METL-Global (20M parameters) took between 7-45 minutes for small datasets (320 examples) and 40-150 minutes for large datasets (20,480 examples).

### Target model dataset splits

We created comprehensive train, validation, and test splits to evaluate performance with small training set sizes and a range of extrapolation tasks, including position, mutation, regime, and score extrapolation. For small training set sizes, we first sampled a random 10% test set from each full dataset. Then, from the remaining data, we sampled datasets of sizes 10, 20, 40, 80, 160, 320, 640, 1280, 2560, 5120, 10240, and 20480. To account for especially easy or difficult training sets that may be sampled by chance, we generated multiple replicates for each dataset size. The number of replicates decreases as the dataset size increases: 101 replicates for the smallest dataset size, followed by 23, 11, 11, 11, 11, 7, 7, 5, 5, 3, and 3 replicates for the largest dataset size. We split the sampled datasets into 80% train and 20% validation sets. We used the same test set across all dataset sizes and replicates. We report median performance metrics across replicates.

Whereas the small dataset splits are sampled randomly, the extrapolation splits are specially designed to assess the models’ ability to generalize to more challenging test sets. For position, mutation, and score extrapolation, we randomly resampled any datasets with *>* 50000 variants down to 50000 variants before generating the extrapolation splits. To account for random effects, we generated 9 replicate splits for each extrapolation type. We report the median across the 9 replicates.

Position extrapolation tests the ability of a model to generalize to sequence positions not present in the training data. To generate position extrapolation splits, we first randomly designated 80% of sequence positions as train and the other 20% as test. Then, we divided all variants (single- and multi-mutant) into training and testing pools depending on whether the variants contain mutations only in positions designated as train or only in positions designated as test. If a multi-mutant variant had mutations in both train and test positions, we discarded it. To create the final train, validation, and test sets, we split the train pool into randomly sampled 90% train and 10% validation sets. We used the entire test pool as the test set.

Mutation extrapolation tests the ability of a model to generalize to mutations not present in the training data. To generate mutation extrapolation splits, we followed a similar procedure as position extrapolation, except with mutations instead of sequence positions. We randomly designated 80% of mutations present in the dataset as train and the other 20% as test. We divided all variants (single- and multi-mutant) into training and testing pools depending on whether the variants contain only mutations designated as train or only designated as test. If a multi-mutant variant had mutations that were designated as train and test, we discarded it. We split the train pool into randomly sampled 90% train and 10% validation sets and used the entire test pool as the test set.

Regime extrapolation tests the ability of the model to generalize from lower numbers of amino acid substitutions to higher numbers of amino acid substitutions. For datasets with single and double substitution variants, we divided the variants into a train pool comprising of the single substitution variants and a test pool comprising of the double substitution variants. We split the train pool into into an 80% train and a 20% validation set. We sampled a 10% test set from the test pool. For datasets containing greater than double substitution variants, we also implemented another regime extrapolation split where the train pool was comprised of single and double substitution variants and the test pool was comprised of variants with three or more substitutions.

Score extrapolation tests the ability of a model to generalize from low-scoring variants to high-scoring variants. We divided variants into train and test pools depending on whether the variant had a score less than wild-type (train pool) or greater than wild-type (test pool). We split the train pool into a 90% train and a 10% validation set and used the entire test pool as the test set.

### Baseline models

We implemented and evaluated additional baseline models: Linear, a fully-connected neural network (FCN), a sequence convolutional neural network (CNN), METL-Local with random initialization, Rosetta’s *total score* as a standalone prediction, and linear regression with Rosetta *total score* (Linear-RTS).

Linear is a linear regression model that uses one hot encoded sequences as inputs. One hot encoding captures the specific amino acid at each sequence position. It consists of a length 21 vector where each position represents one of the possible amino acids or the stop codon. All positions are zero except the position of the amino acid being encoded, which is set to a value of one. Note that we removed variants containing mutations to the stop codon during dataset preprocessing, so this feature was not used in our analysis. We implemented linear regression using scikit-learn’s ridge regression class, which incorporates L2 regularization. We set the solver to *cholesky* to calculate a closed-form solution for the ridge regression weights. We set *alpha*, the constant that controls regularization strength, to the default value of 1.0. We set all other parameters to the default scikit-learn values.

For baseline neural networks, we tested an FCN, a CNN, and a transformer encoder with a similar architecture as METL-Local, but with a random initialization. The FCN and CNN used one hot encoded sequences as input. The FCN consisted of 1 hidden layer with 1024 nodes followed by a dropout layer with a dropout rate of 0.2. The CNN consisted of 1 convolutional layer with kernel size 7, 128 filters, and zero-padding to ensure the output has the same shape as the input (padding mode “same” in PyTorch’s Conv2d class). Following the convolutional layer, we placed a fully-connected layer with 256 nodes and a dropout layer with a dropout rate of 0.2. We used the ReLU activation function for both models. In addition to the FCN and CNN, we tested a randomly initialized transformer encoder neural network with a similar architecture as METL-Local. Unlike METL-Local, this randomly initialized version was set up with a single output node corresponding to the experimental functional score instead of multiple output nodes corresponding to Rosetta scores.

We trained the FCN, CNN, and randomly initialized METL-Local for 500 epochs using the AdamW optimizer with a base learning rate of 0.001. We applied a learning rate scheduler with linear warm-up and cosine decay, with a warm-up period of 2% of the total training steps. Additional AdamW hyperparameters were set as follows: *weight*_*decay* = 0.1, *β*_1_ = 0.9, *β*_2_ = 0.999, and *ε* = 1*e* − 8. We used a batch size of 128 and mean squared error loss. We applied gradient norm clipping with a max norm of 0.5. Similar to METL-Local finetuning, we selected the model from the epoch with the lowest validation loss when the validation set size was ≥ 32. Otherwise, we used the model from the last epoch of training.

We evaluated Rosetta’s *total score* as a standalone, unsupervised prediction, as well as an additional input feature for linear regression, which we refer to as Linear-RTS. By default, the lower Rosetta’s *total score*, the more stable the structure is predicted to be. Thus, when using Rosetta’s *total score* as an unsupervised prediction, we multiplied it by −1 before computing correlation with the experimental functional score. We also tested Rosetta’s *total score* as part of a supervised learning framework. Linear-RTS is identical to Linear, but it uses Rosetta *total score* as an additional input feature in combination with the one hot encoded sequence in an augmented regression setting [17]. We standardized the *total score* for use as an input feature by first calculating its mean and standard deviation in the train set. Then, we subtracted the mean and divided by the standard deviation.

### Comparison to ESM-2

We used the ESM-2 [19] 35M parameter model with identifier `esm2_t12_35M_UR50D` as our default ESM model so that the comparisons with the 20M parameter METL-Global model would primarily emphasize their different pretraining strategies rather than model size. We incorporated several additional layers to match the METL architecture, including a global mean pooling layer, a dropout layer with dropout rate 0.5, and a linear prediction head. We attached these additional layers immediately after layer 12. We trained the ESM-2 models using the same training procedures we used for the METL models. We also explored feature extraction with larger 150M and 650M parameter ESM-2 models (identifiers `esm2_t30_150M_UR50D` and `esm2_t33_650M_UR50D`). For these larger models, we attached the additional layers after layers 30 and 33, respectively.

### Comparison to RaSP

We compared METL to RaSP [87] using the pre-trained model weights for both the cavity model and downstream models shared by the authors on their GitHub repository (https://github.com/KULL-Centre/_2022_ML-ddG-Blaabjerg). RaSP is a relevant comparison to METL since it is trained on ΔΔ*G* values predicted using the Rosetta `cartesian_ddg` protocol [87]. Since RaSP is a point mutation stability predictor, it is not designed to handle mutants with multiple mutations (multi-mutants). After consulting with the authors, we adapted RaSP to handle multi-mutants by assuming an additive effect for each mutation in multi-mutants. As a result, we scored multi-mutants by scoring each point mutation individually and adding their scores. We used default parameters to extract the atomic environment for each mutant. After extracting the atomic environment, we used RaSP’s cavity model to get a vectorized representation of the atomic environment followed by the ensemble of downstream models to predict the stability effect of a variant. We used RaSP in a zero-shot setting, i.e. we did not finetune the model on examples from our target experimental dataset. Lastly, the authors note that RaSP is neither trained nor evaluated on disulfide-bonded cystine residues since they cannot be predicted using the Rosetta protocol used to generate RaSP’s training data [16]. Since the goal of our evaluations was to test how well models predict protein functions measured by the various assays, we do not filter our test data based on this criteria.

### Comparison to EVE

We obtained multiple sequence alignments (MSAs) for GB1, Ube4b, GFP, and Pab1 from the EVcouplings web server [147] in March 2023. We obtained MSAs for TEM-1, GRB2, and DLG4 in July 2023 and for PTEN in September 2024. We queried the UniRef100 database with search parameters consistent with those in EVMutation [148]: a position sequence filter of 70 percent, a sequence fragment filter of 50 percent, 100 percent removal of similar sequences, and 80 percent down weighting of similar sequences. We started our bitscore value at 0.5 bits per residue. If we did not have 80 percent sequence coverage, we increased the threshold by 0.05 bits per residue until the constraint was satisfied. If the number of effective sequences in the alignment was less than 10 times the length of the protein, we decreased the bits per residue until the requirement was satisfied. We prioritized the number of effective sequences objective if the two were in conflict. We trained EVE using the default training parameters of 40,000 training iterations, sampling 20,000 evolutionary indices, and a default theta reweighting value of 0.2 to preprocess the MSA. We made mutation effect predictions for every position in the sequence by capitalizing all amino acids in the MSA.

In addition to using EVE as a standalone zero-shot method, we incorporated the EVE score into a supervised learning model. We selected EVE for augmented regression instead of the models evaluated by Hsu et al. [17] because EVE outperforms them in ProteinGym’s zero-shot substitution deep mutational scanning evaluation [51], therefore providing a stronger baseline. The augmented regression model Linear-EVE is identical to the Linear model described above, but it uses the EVE score as an additional input feature in combination with the one hot encoded protein sequence. We standardized the EVE score for use as an input feature by first calculating its mean and standard deviation in the train set. Then, we subtracted the mean and divided by the standard deviation.

### Comparison to ProteinNPT

We ran the full ProteinNPT [18] pipeline, including the optional step of computing zero-shot fitness predictions with MSA Transformer [149] and incorporating them as auxiliary labels. The authors state this optional step helps improve performance, especially for position extrapolation. We followed the instructions from the ProteinNPT GitHub repository (https://github.com/OATML-Markslab/ProteinNPT) and used the model configuration defined in `PNPT_final.json`. This configuration specifies the MSA Transformer model with identifier `esm_msa1b_t12_100M_UR50S` as the sequence embedding model and 10,000 total training steps. We used the same MSAs obtained from the EVCouplings web server that we used for EVE (see above).

### Calculating predicted epistasis

For the GB1 epistasis analysis, we computed predicted epistasis using the pretrained METL-Local GB1 model. Let score(*S*) denote the model-predicted Rosetta *total_score* for variant *S*, and let wt_score denote the predicted *total_score* for the wild-type sequence. For each possible single and double variant, we first computed its effect relative to wild type:

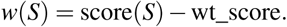

Then, we computed epistasis as:

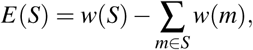

where *m* represents each single mutation in variant *S*. To compute pairwise positional epistasis, we calculated the mean absolute epistasis across all variants with mutations in the given pair of positions.

### GFP sequence design

We finetuned a pretrained METL-Local model on 64 randomly sampled variants from the GFP dataset. The selected variants had 1 to 11 mutations, and their experimental score distribution was bimodal (Fig. S19), similar to the distribution of the full GFP dataset. We refer to the finetuned METL-Local GFP model in this low-N setting as METL-L-GFP. We inspected the extrapolation behaviors of the METL-L-GFP model for increasing number of mutations. For increasing numbers of mutations selected with simulated annealing, the predicted brightness approximately stabilized at a positive value (Fig. S25), in contrast to what has been observed in convolutional neural networks [150]. Conversely, for increasing numbers of randomly selected mutations, the predicted brightness stabilized at a negative value (Fig. S26). That the predicted scores did not continue to increase positively or negatively with the number of mutations was a basic verification of the METL-L-GFP model’s extrapolation properties.

We performed in silico optimization with METL-L-GFP to design a total of 20 variants distributed evenly across 4 different design criteria. These criteria are the product of 2 primary design categories: the number of mutations (either 5 or 10) and the constraints on mutation selection (either Observed or Unobserved). In the Observed constraint, the designed sequences contain only amino acid substitutions found in the 64-variant training set. Conversely, in the Unobserved constraint, the designed sequences exclude any amino acid substitutions found in the 64-variant training set. The combinations of these categories resulted in the 4 design criteria: Observed 5-mutant, Unobserved 5-mutant, Observed 10-mutant, and Unobserved 10-mutant. We designed 5 sequences for each criterion, resulting in a total of 20 designed sequences.

To perform the in silico optimization, we ran simulated annealing 10,000 times for each design criterion. For each simulated annealing run, we changed the random seed and executed the Monte Carlo optimization for 10,000 steps. Each step consisted of suggesting a mutation for the currently sampled variant and deciding whether to accept the new variant according to the Metropolis-Hastings criteria. We decreased the optimization temperature according to a logarithmic gradient beginning at 10^1^ and ending at 10^−2^. The initial temperature was chosen by randomly sampling 10,000 variants, predicting their brightness with METL-L-GFP, and calculating the absolute value of the difference between the lowest and highest predicted brightness, rounded to the nearest power of 10. The final temperature was determined by calculating the absolute value of the smallest difference in predicted brightness between any two variants in the 64 variant training set, rounded to the nearest power to 10. The initial temperature encouraged acceptance of all variants, while the final temperature meant that only variants better than the current ones would be accepted.

The simulation began by randomly selecting a variant with the necessary number of mutations depending on the design criterion. We determined how many mutations to change at each step by sampling from a Poisson distribution. To generate a new variant from an existing one, we first determined the difference between the number of mutations to change and the maximum allowable mutations, which indicated the number of current mutations to keep, *m*. We randomly sampled which *m* mutations to keep, and reset the other mutations to wild type. Subsequently, we compiled all feasible single mutations of the sequence with the *m* existing mutations and randomly sampled new mutations without replacement until the variant mutation count reached the maximum allowable mutations.

The optimization process described above yielded 10,000 designs for each criterion, which we downsampled to 5 designs for each criterion via clustering. Our downsampling approach prioritized diversity and was predicated on the idea that repeated convergence to similar sequences may correlate with higher true fitness values, as these regions of the fitness landscape would have broader peaks and allow more room for error in the model predictions or optimization process. We clustered the 10,000 designs using scikit-learn’s agglomerative clustering with complete linkage and a BLOSUM62-based distance metric. Because selecting 10, 20, or 50 clusters did not substantially impact the diversity of the selected mutations, we chose 20 clusters. We then removed clusters that contained less than 100 sequences, which represented 1% of the simulated annealing solutions.

To select 5 (or 10) clusters from those remaining, we employed an iterative, greedy approach. We identified a representative sequence for each cluster, choosing the one with the lowest average BLOSUM62-based distance to all other sequences within the same cluster. To initialize, we selected the largest cluster. We then proceeded iteratively, selecting additional clusters one at a time. In each iteration, we calculated the distances between the representative sequences of the already selected clusters and the remaining unselected clusters. We selected the cluster with the largest mean distance to the already selected clusters to promote sequence diversity. The GFP sequence designs were the representative sequences from the selected clusters.

To generate the baseline random GFP variants, we used two different random sampling algorithms corresponding to the different design criteria. For the Observed random variants, we randomly sampled individual mutations without replacement from the 209 unique mutations in the training set. For the Unobserved random variants, we randomly sampled individual mutations without replacement from all other all possible mutations excluding those 209 in the training set.

### Cloning and experimental validation of GFP variants

We modeled our expression system on that used in Sarkysian et al. [36], which uses a pQE-30 vector (Qiagen) to express GFP as a fusion protein with mKate2. To generate the expression construct, we used the vector backbone from a related pQE-30 system that expresses KillerOrange (Addgene 74748) and ordered a gene encoding the mKate2-GFP fusion protein from Twist Biosciences. We first removed a BsaI restriction site in the *ampR* gene from the backbone using site directed mutagenesis (NEB M0554S) and then used Golden Gate cloning to replace the *killerorange* gene with the fusion protein. We incubated (1 hr, 37 C) the backbone and insert with BsaI (15 U, NEB: R3733), T4 Ligase (1,000 U, NEB M0202), and T4 Ligase Buffer (NEB B0202) to assemble the vector. The assembly was cleaned up with a PCR Clean and Concentrate column (Zymogen D4003) and transformed into in-house DH5a cells. Plasmid stock was purified from an overnight culture starting from a single colony using a Qiagen Miniprep kit (Qiagen 27104), and the vector was on-boarded with Twist Biosciences. All GFP variants were ordered as clonal genes from Twist Biosciences wherein the wild-type *gfp* gene sequence was replaced with the variant sequence. For each variant, the nucleotide sequence was kept the same as the wild-type sequence except at mutated residues. We selected new codons for mutated residues based on an *E. coli* codon usage index [151] to mitigate poor expression due to rare codons.

Clonal genes ordered from Twist Biosciences were transformed into NEBExpress Iq Competent *E. coli* (NEB C3037I) cells and plated on Luria Broth (LB) plates with carbenecillin selection (0.1 mg/mL). Proteins were expressed as previously described in Sarkysian et al. [36]. Briefly, freshly plated transformants were incubated overnight at 37 ^◦^C and then moved to 4 °C the following day. After 24 hours, plates were washed with 4 mL LB and normalized to 1 OD. This wash was used to create triplicate expression cultures where protein expression was induced for 2 hours with 1 mM IPTG at 23 ^◦^C. An empty pQE-30 vector was used as a negative expression control.

To prepare cultures for fluorescence measurement, expression cultures were pelleted (3,000xg, 5 mins) and re-suspended in sterile 1X PBS to a concentration of 1 OD. Cells were diluted 2-fold into 96-well plates to measure fluorescence and culture density. Measurements were taken with either the Tecan Spark 10M or the Tecan Infinite M1000 Pro. Measurements for GFP (ex. 405 nm, em. 510 nm), mKate2 (ex. 561 nm, em. 670 nm), and OD600 (abs. 600 nm) were collected.

Relative brightness was reported as the ratio of GFP fluorescence to mKate2 fluorescence averaged across replicates. First, raw fluorescent measurements were normalized to cell density by dividing by the sample’s OD600 value. The background fluorescence signal was subtracted out of each sample. The background fluorescence signals for GFP and mKate2 were measured from negative control cells containing no fluorescent protein. A sample’s relative brightness was calculated for each sample by dividing the normalized background-subtracted GFP fluorescence by the normalized background-subtracted mKate2 fluorescence. All fluorescent values were normalized to wildtype avGFP.

### Visualizations

We used FreeSASA [152] to compute relative solvent accessibility (RSA), which was used to color the points in Figures 5d, S15, and S16. We used Multidimensional Scaling (MDS) from scikit-learn to visualize GFP designs in Figure 7b. MDS is a dimensionality reduction technique that preserves relative distances between observations [153]. We used Hamming distance between sequences, which had the effect of making variants show up in concentric circles around the wild-type sequence based on the number of mutations from wild-type.

## Data availability

Pretrained METL models are available at doi:10.5281/zenodo.11051644 [154]. Rosetta simulation datasets are available at doi:10.5281/zenodo.10967412 [155]. Additional data is available in the GitHub repository https://github.com/gitter-lab/metl-pub, which is archived at doi:10.5281/zenodo.10819536 [156]. The PDB structure identifiers used to train METL-Global are listed in Table S2. The PDB and AlphaFold DB structure identifiers used for METL-Local are listed in Table S7. Experimental datasets used for model evaluation are listed in Table S8 with references and corresponding filenames.

## Code availability

We provide a collection of METL software repositories to reproduce the results of this manuscript and run METL on new data.

- Code for pretraining and finetuning METL PLMs is available at github.com/gitter-lab/metl. An archived version is available at doi:10.5281/zenodo.10819483 [157].
- Code for generating biophysical attributes with Rosetta is available at github.com/gitter-lab/metl-sim. An archived version is available at doi:10.5281/zenodo.10819523 [158].
- Code for making predictions with pretrained METL PLMs is available at github.com/gitter-lab/metl-pretrained. An archived version is available at doi:10.5281/zenodo.10819499 [159].
- Supporting code and data to reproduce this study is available at github.com/gitter-lab/metl-pub. An archived version is available at doi:10.5281/zenodo.10819536 [156].
- A Hugging Face wrapper and demo for METL models is available at huggingface.co/gitter-lab/METL. All code is available under the MIT license.

## Unique biological materials

All unique biological materials are available upon request from the authors.

## Acknowledgments

This research was supported by National Science Foundation awards 2226383 (P.A.R.) and 2226451 (A.G.), National Institutes of Health awards R01GM135631 (A.G.) and R35GM119854 (P.A.R.), the John W. and Jeanne M. Rowe Center for Research in Virology at the Morgridge Institute for Research, generous support from Jeanne M. Rowe, and the University of Wisconsin–Madison Office of the Vice Chancellor for Research and Graduate Education with funding from the Wisconsin Alumni Research Foundation. The funders had no role in study design, data collection and analysis, decision to publish or preparation of the manuscript. We thank Ben Gelman for insightful discussions regarding the transformer architecture, attention mechanism, and effects of data normalization; Jerry Duan for the initial molecular simulations implementation; Brian Aydemir for testing and feedback on the molecular simulation workflow; and Kaustubh Amritkar for collecting and processing PDB files for METL-Global. The research was performed using the computing resources and assistance of the University of Wisconsin-Madison Center for High Throughput Computing [160] and services provided by the OSG Consortium [42, 161, 162], which is supported by National Science Foundation awards 2030508 and 1836650.

## Author Contributions

S.G., A.G., and P.A.R. conceived the original idea for the study. S.G. developed the METL framework and conducted the computational experiments and analyses. S.G., A.G., P.A.R., B.J., and A.S. contributed to discussions on computational experiment design and results interpretation. B.J. implemented the EVE baseline, generated GFP variant designs, and created an interactive Jupyter notebook to run simulations. A.S. implemented the RaSP baseline. S.D. acquired and prepared PDB structures for METL-Global. J.P. developed the Hugging Face integration, interactive demo, and Google Colab notebooks. C.F. designed and performed the wet lab experiments. A.G. and P.A.R. supervised the research. S.G., A.G., P.A.R., C.F., B.J., and A.S. wrote the manuscript. All authors reviewed and edited the manuscript.

## Supplementary Information

**Figure S1.**
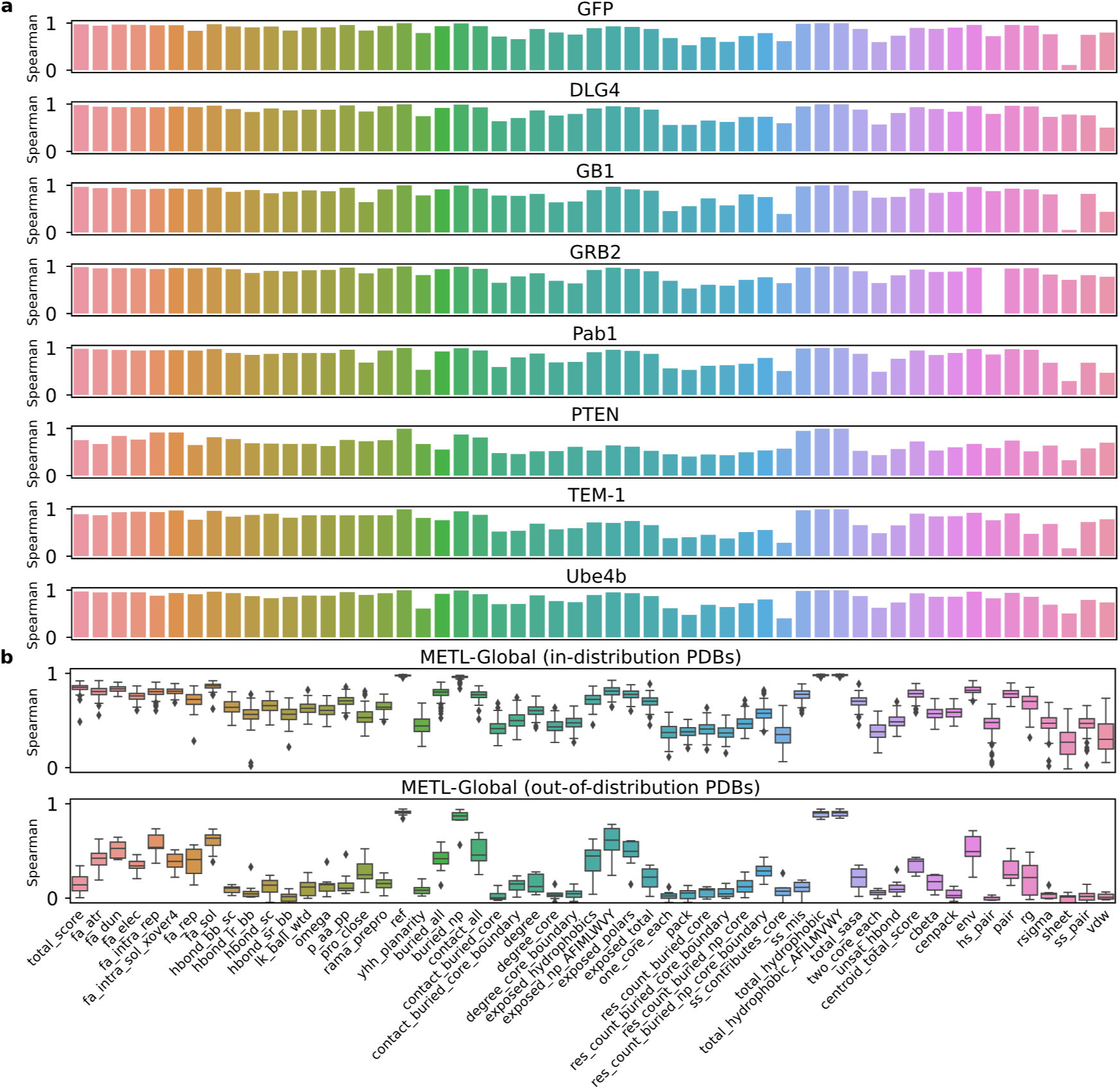
Performance of pretrained METL source models in predicting Rosetta scores. This figure shows Spearman correlations between true and predicted Rosetta scores for each of the 55 Rosetta score terms. (a) Correlation of METL-Local models predicting Rosetta biophysical attributes for the protein-specific pretraining data test sets. (b) Correlation of METL-Global in predicting Rosetta scores for protein variants originating from in-distribution base PDBs (those included in METL-Global pretraining, *n* = 148) and out-of-distribution base PDBs (those not included, *n* = 8). To evaluate in-distribution PDBs, we used variants in the pretraining data test set. To evaluate out-of-distribution PDBs, we used variants from the deep mutational scanning datasets included in this study. METL-Global performs substantially better for in-distribution PDBs, suggesting there is overfitting to the PDBs present in the training data.

**Figure S2.**
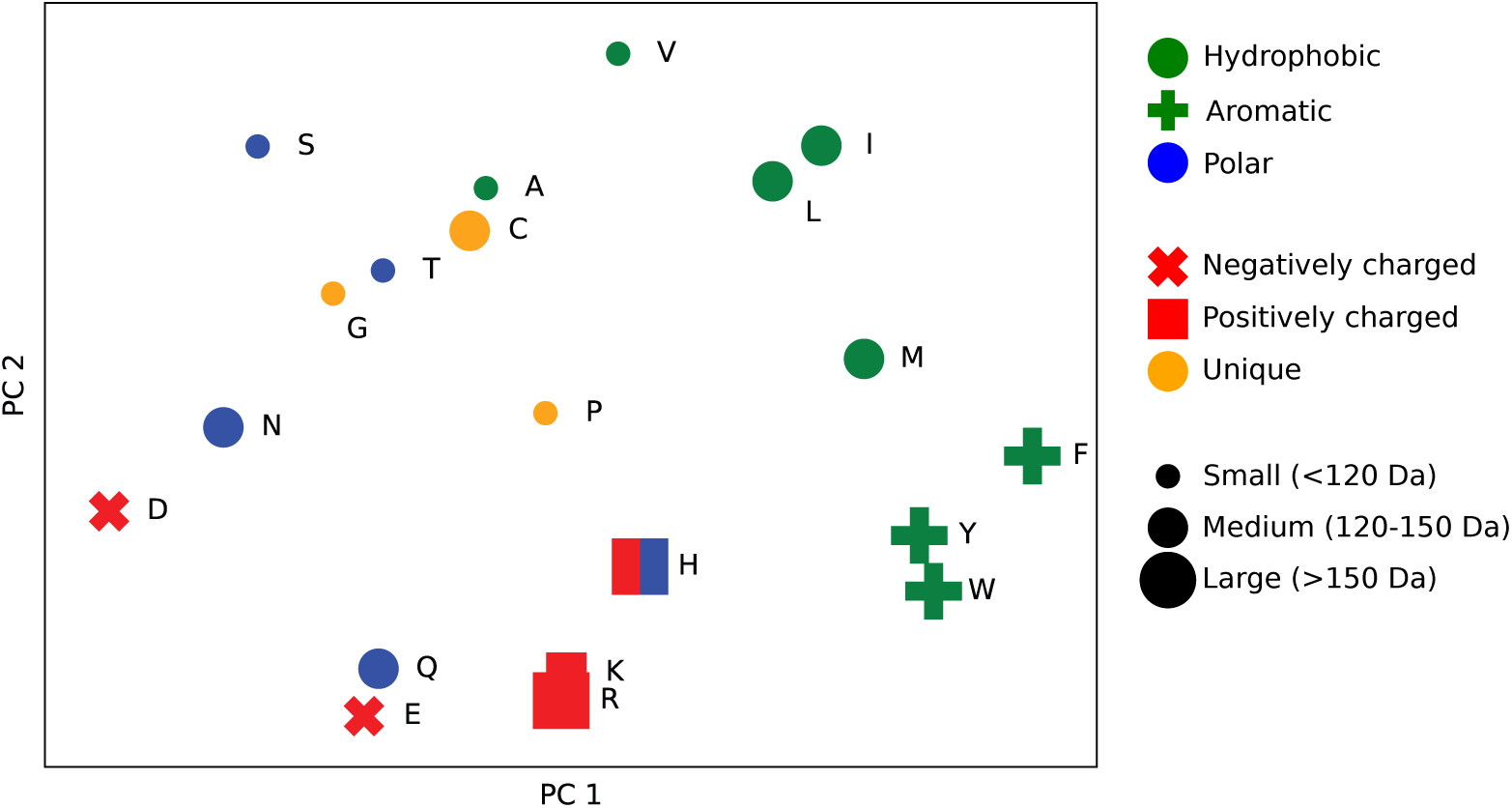
METL-Global amino acid embeddings. We applied principal component analysis (PCA) to reduce the METL-Global length 512 amino acid embeddings down to 2 dimensions, capturing 33% of the variance in data. This scatter plot of the 2-dimensional amino acid embeddings is annotated with amino acid properties. METL-Global groups amino acids with similar biochemical properties in the embedding space, like protein language models (PLMs) trained on millions of natural protein sequences [1].

**Figure S3.**
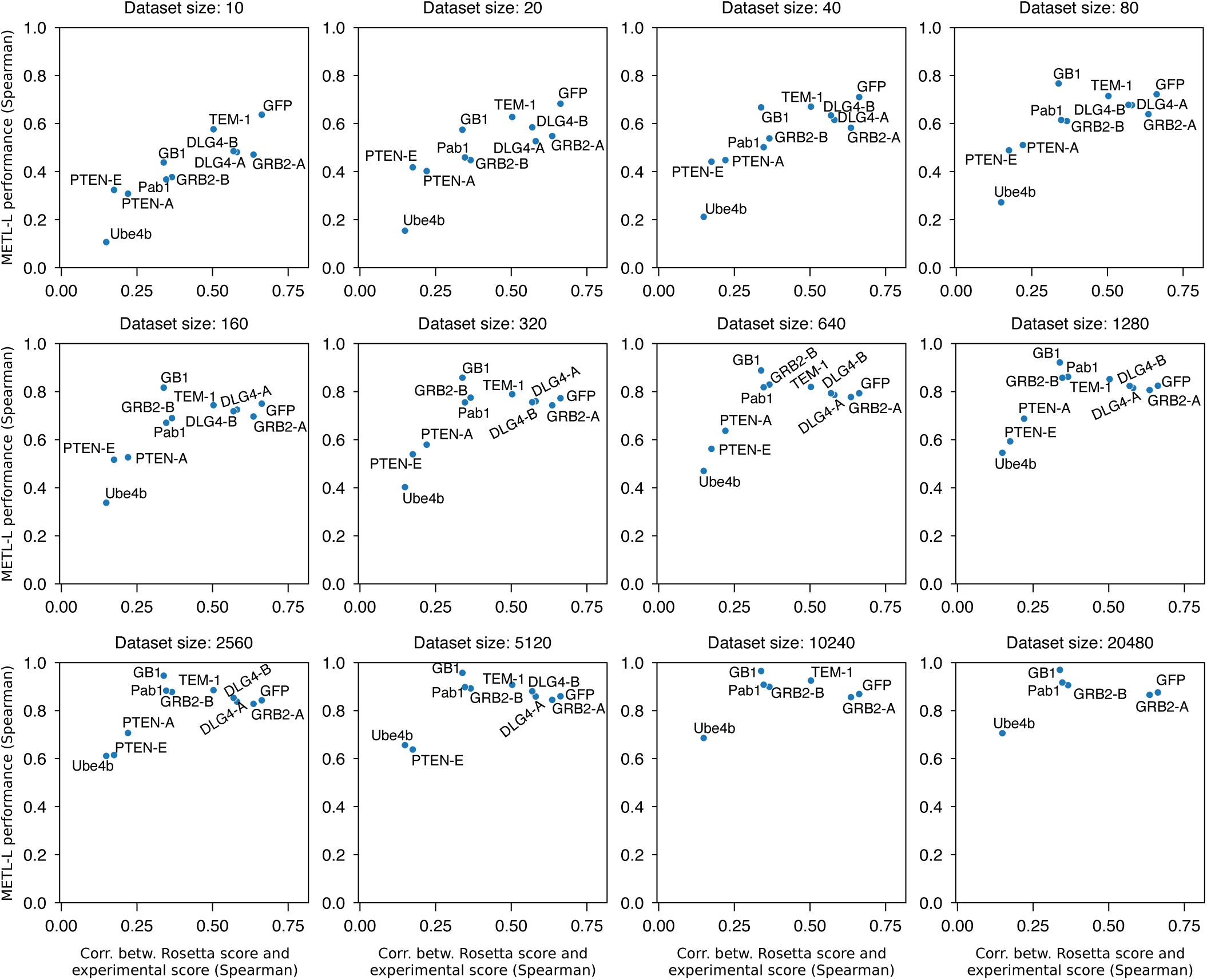
Relationship between METL-Local performance and the relatedness of Rosetta and experimental scores. The figure displays a series of scatterplots showing the relationship between METL-Local performance and the relatedness of Rosetta and experimental scores, across multiple experimental datasets and training set sizes. The x-axis shows the Spearman correlation between Rosetta total score and the experimental functional score for the entire dataset, representing the similarity between the Rosetta total score and the experimental functional score. The y-axis shows the METL-Local performance for the respective training set size, as determined by the Spearman correlation on the test set. As the similarity between Rosetta total score and the experimental functional score increases, so does the METL-Local performance, at least for small training set sizes. However, with increasing experimental training set sizes, the similarity between Rosetta total score and experimental functional score becomes less important to the METL-Local performance, suggesting a shift in METL-Local away from the Rosetta pretraining data and more toward the experimental finetuning data.

**Figure S4.**
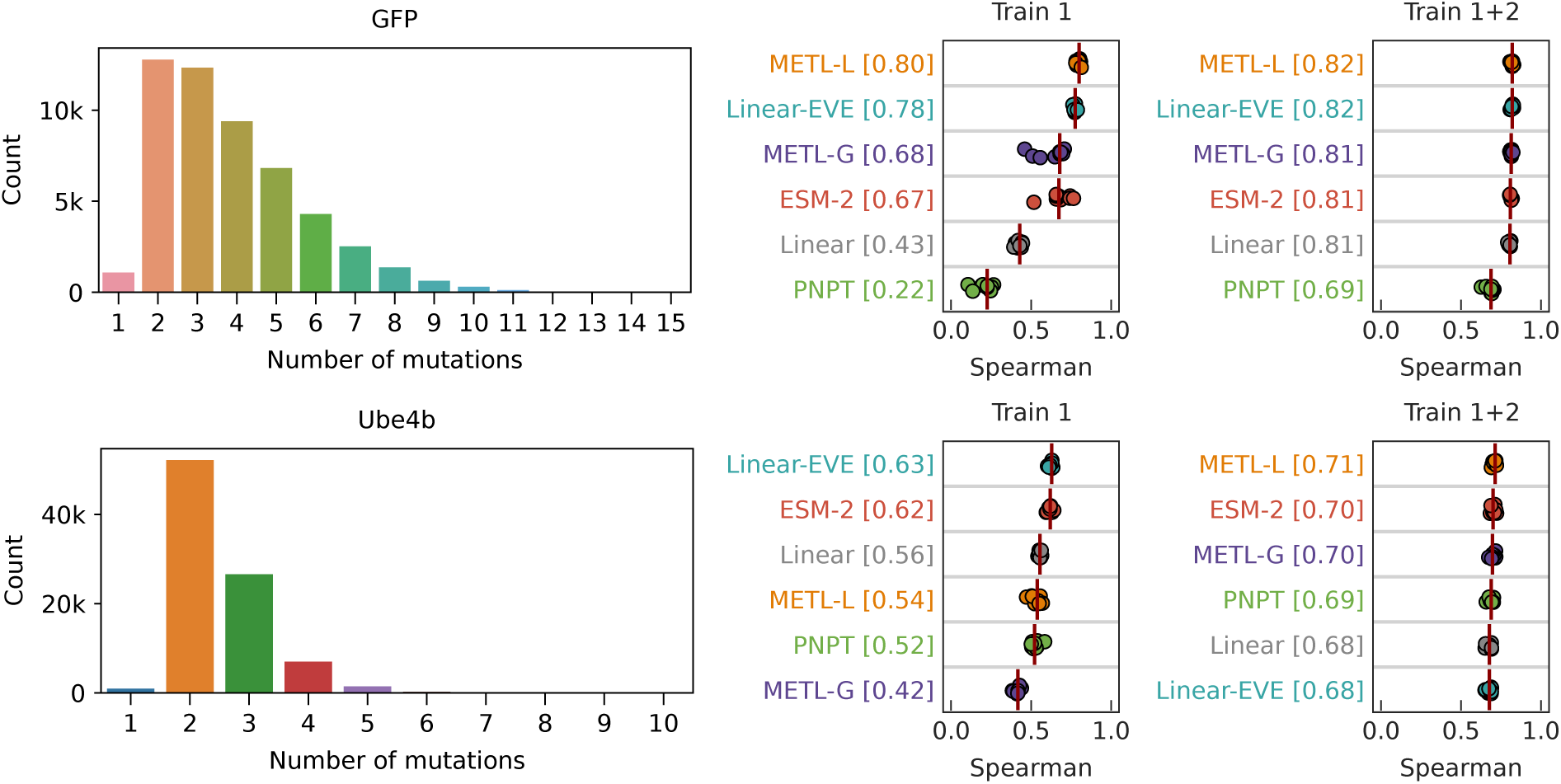
Regime extrapolation for GFP and Ube4b datasets. The GFP and Ube4b datasets contain variants with higher order mutations, enabling us to test two types of regime extrapolation: Train 1 and Train 1+2. The bar plots (left) show the counts of variants with the specified number of mutations for each dataset. The strip plots (right) show the performance of regime extrapolation for Train 1, where we train on single substitution variants and evaluate on variants with 2+ substitutions, and Train 1+2, where we train on variants with single or double substitutions, and evaluate on variants with 3+ substitutions. The strip plots show the performance of 9 test set replicates, and the red vertical line denotes the median.

**Figure S5.**
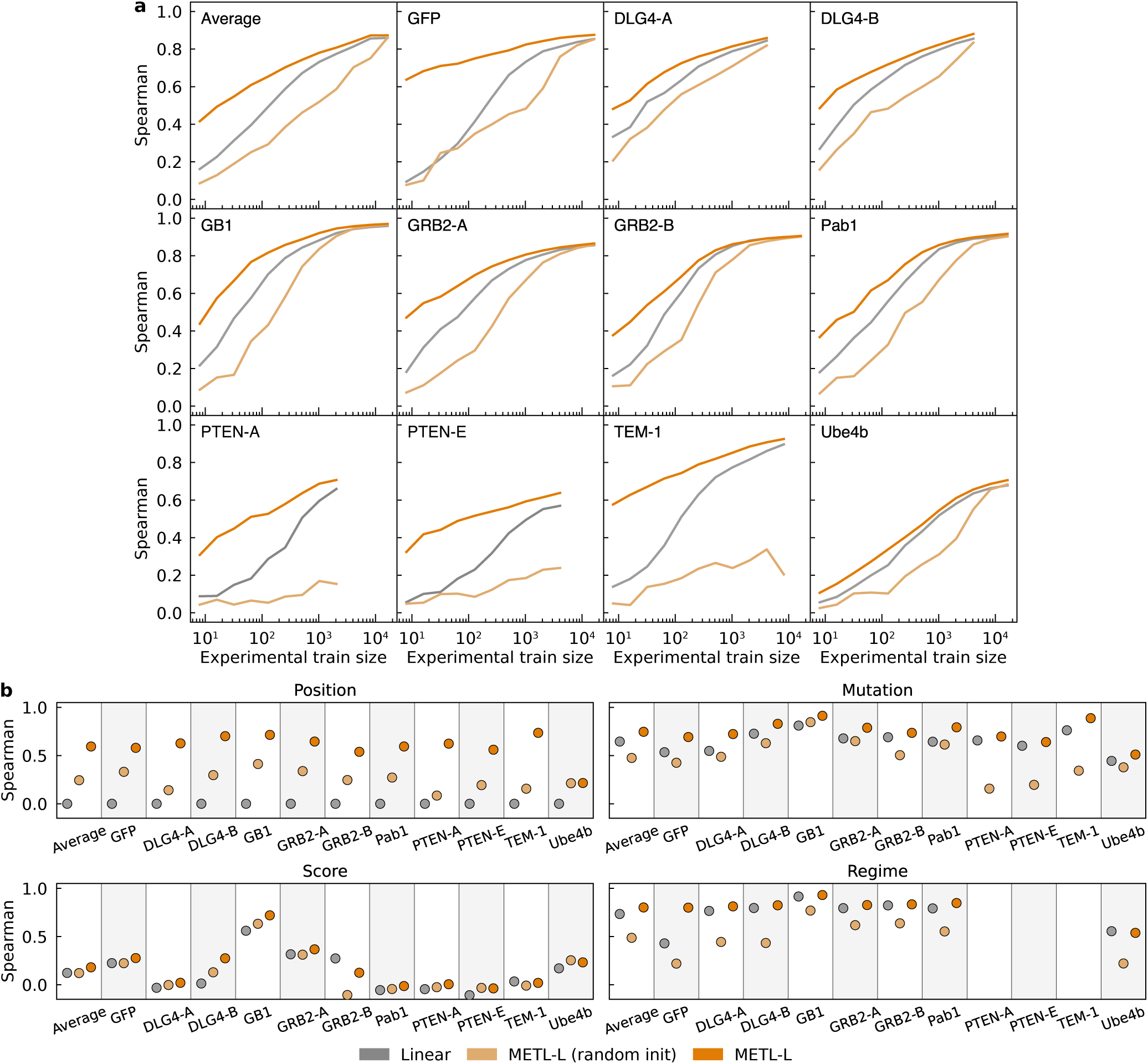
Performance of METL-Local with and without pretraining. These plots show the correlation performance of Linear, METL-Local (random init), and METL-Local. METL-Local (random init) is a model with the same architecture as METL-Local but without pretraining on Rosetta scores. (a) The learning curves show that METL-Local (random init) substantially underperforms both Linear and pretrained METL-Local, emphasizing the impact pretraining on Rosetta scores has on this transformer-based architecture. Given enough experimental training data, METL-Local (random init) converges to the performance of the other models for most datasets. (b) METL-Local (random init) outperforms Linear for position extrapolation due to the fact that Linear is not able to perform position extrapolation but is substantially worse than METL-Local. For the other types of extrapolation, METL-Local (random init) performs about the same or worse than Linear.

**Figure S6.**
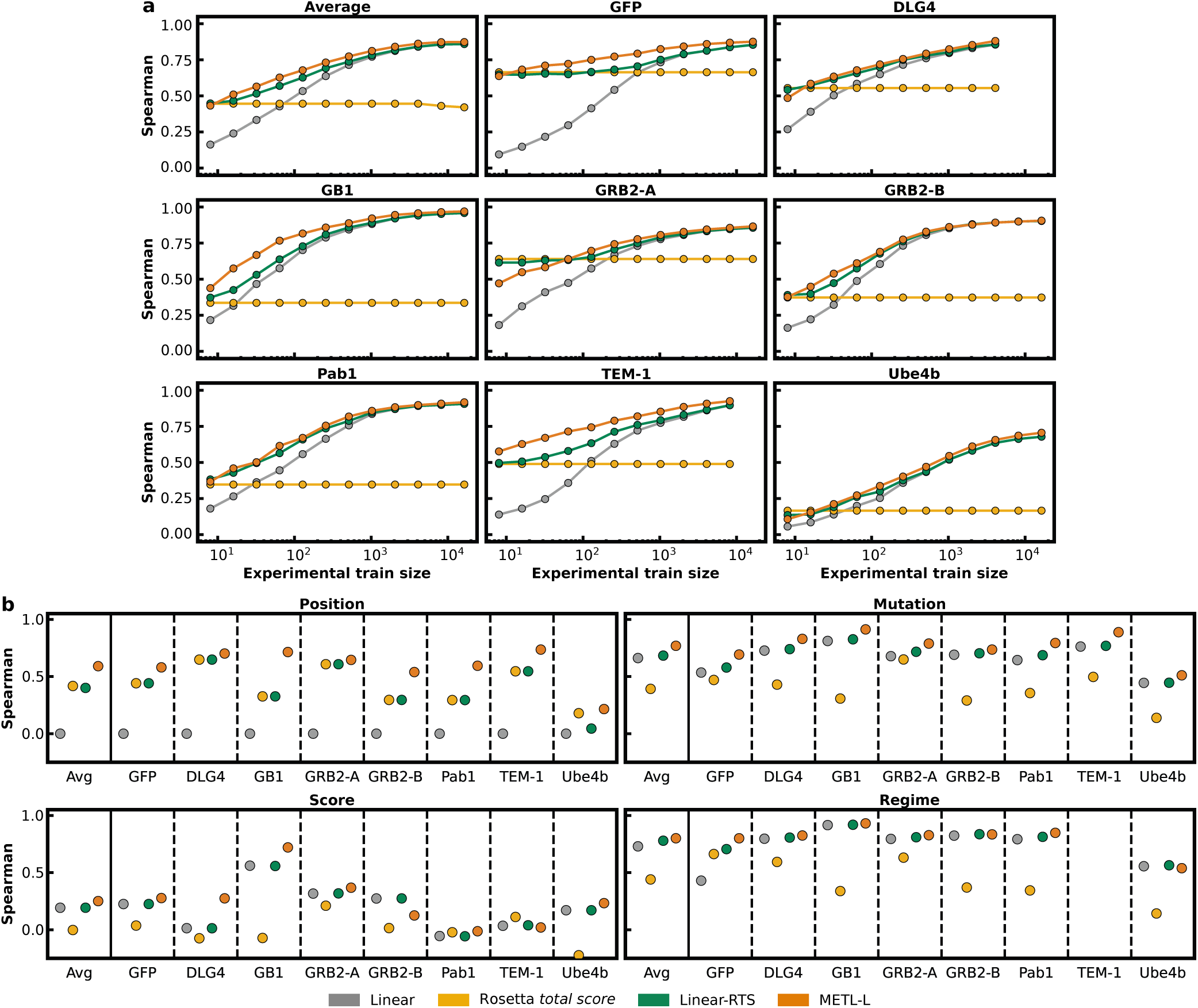
Performance of baseline models directly using Rosetta total score. Rosetta *total score* is the score term from Rosetta with no supervised training on experimental data. Linear-RTS is a linear ridge regression model trained on experimental data with one hot encoding features augmented with the Rosetta *total score* as an additional input feature. Both of these models require running Rosetta to compute the *total score* for every variant, even during inference. For comparison, this figure also shows the performance of Linear and METL-Local. (a) For small training set sizes, incorporating Rosetta *total score* as an additional input feature for ridge regression greatly improved performance over solely using one hot encoding features, as demonstrated by the difference in performance between Linear and Linear-RTS. While Linear-RTS sometimes matched METL-Local’s performance and even exceeded it on the GRB2-A dataset, METL-Local still outperformed Linear-RTS on average by a small amount and is much faster. (b) METL-Local outperformed Linear-RTS across most datasets and extrapolation tasks. The performance differences were sometimes substantial, such as for position extrapolation with GB1. In other cases, the performance differences were much smaller, such as for regime extrapolation. This analysis does not use all of the deep mutational scanning datasets.

**Figure S7.**
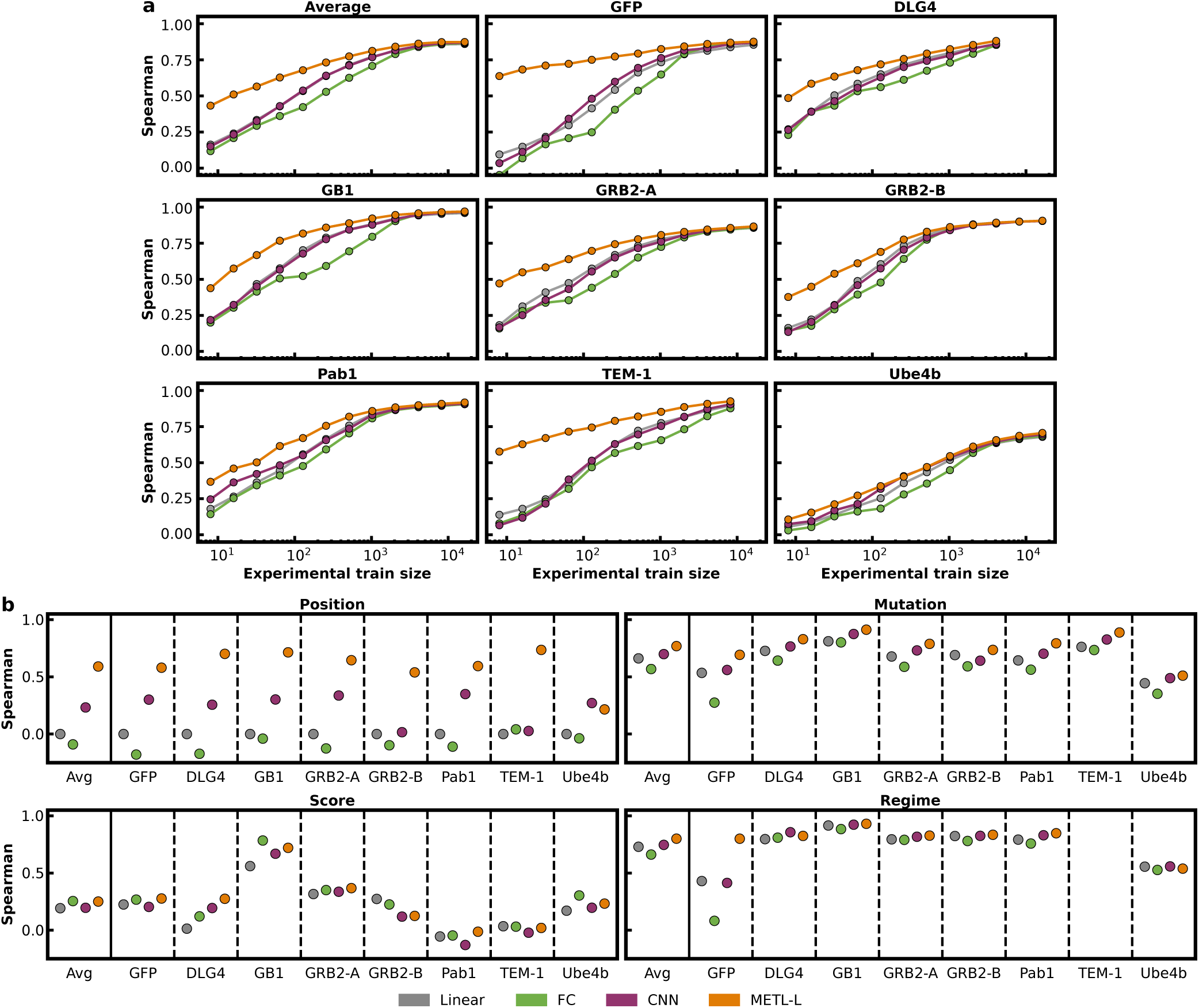
Performance of additional baseline models. Correlation performance of Linear, fully-connected networks (FC), sequence convolutional networks (CNN), and METL-Local. (a) METL-Local has strong advantages over the fully-connected network and CNN on nearly every dataset. The CNN performed about the same as Linear across different sized training sets. The fully-connected network typically performed about the same or worse than Linear, especially for mid-size training sets. (b) METL-Local exhibits much better position extrapolation capabilities than all three baseline models as well as substantially better GFP regime extrapolation. This analysis does not use all of the deep mutational scanning datasets.

**Figure S8.**
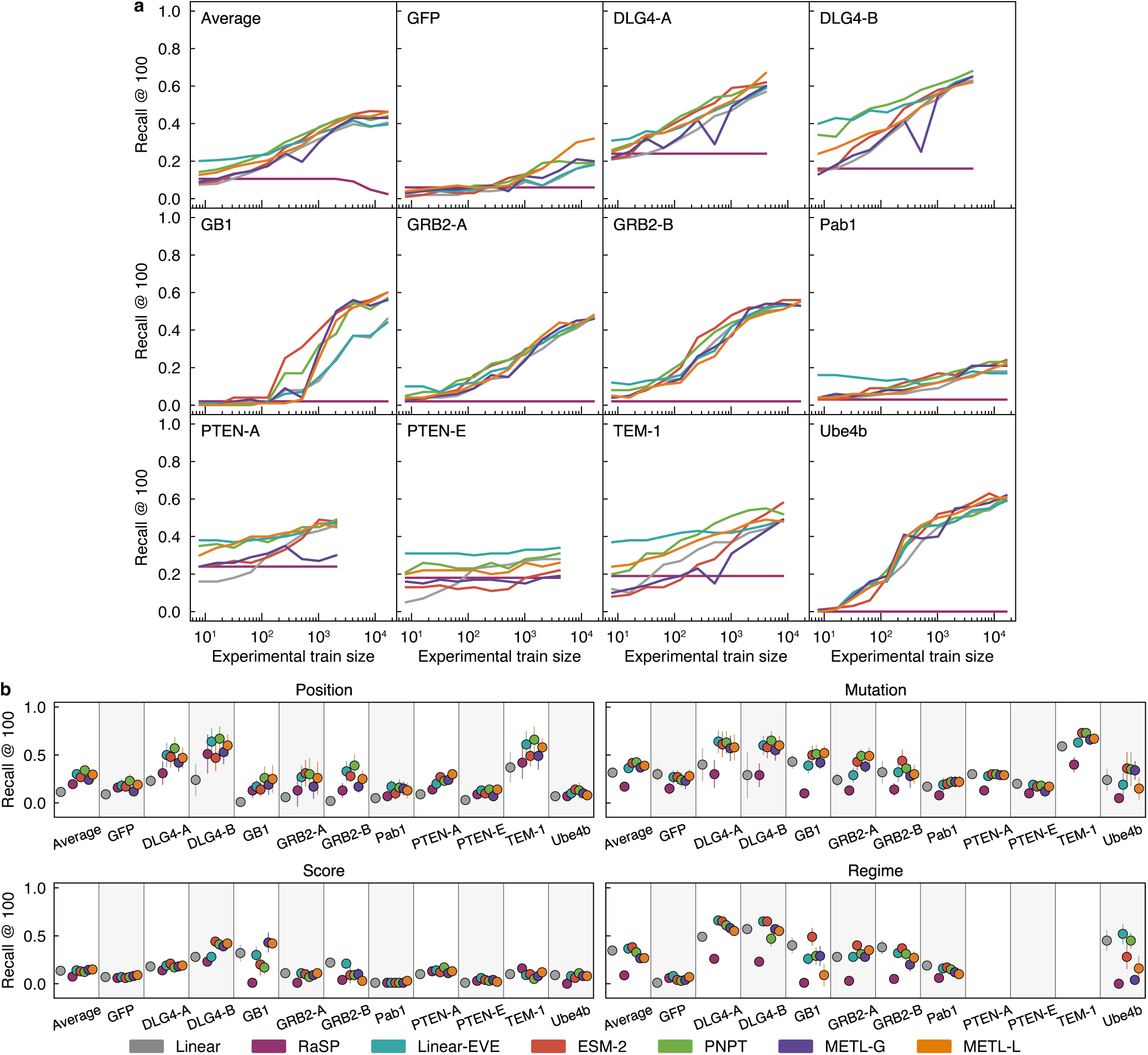
Performance using recall metric. These plots show the fraction of the true top 100 test set variants present within each model’s top 100 predicted variants. (a) Recall performance across training set sizes. Models with strong low-N Spearman correlation do necessarily achieve strong low-N recall. (b) Recall performance for extrapolation tasks. This analysis does not include Rosetta *total score* or EVE. The recall threshold has not been optimized, and using a different recall threshold may show different performance patterns.

**Figure S9.**
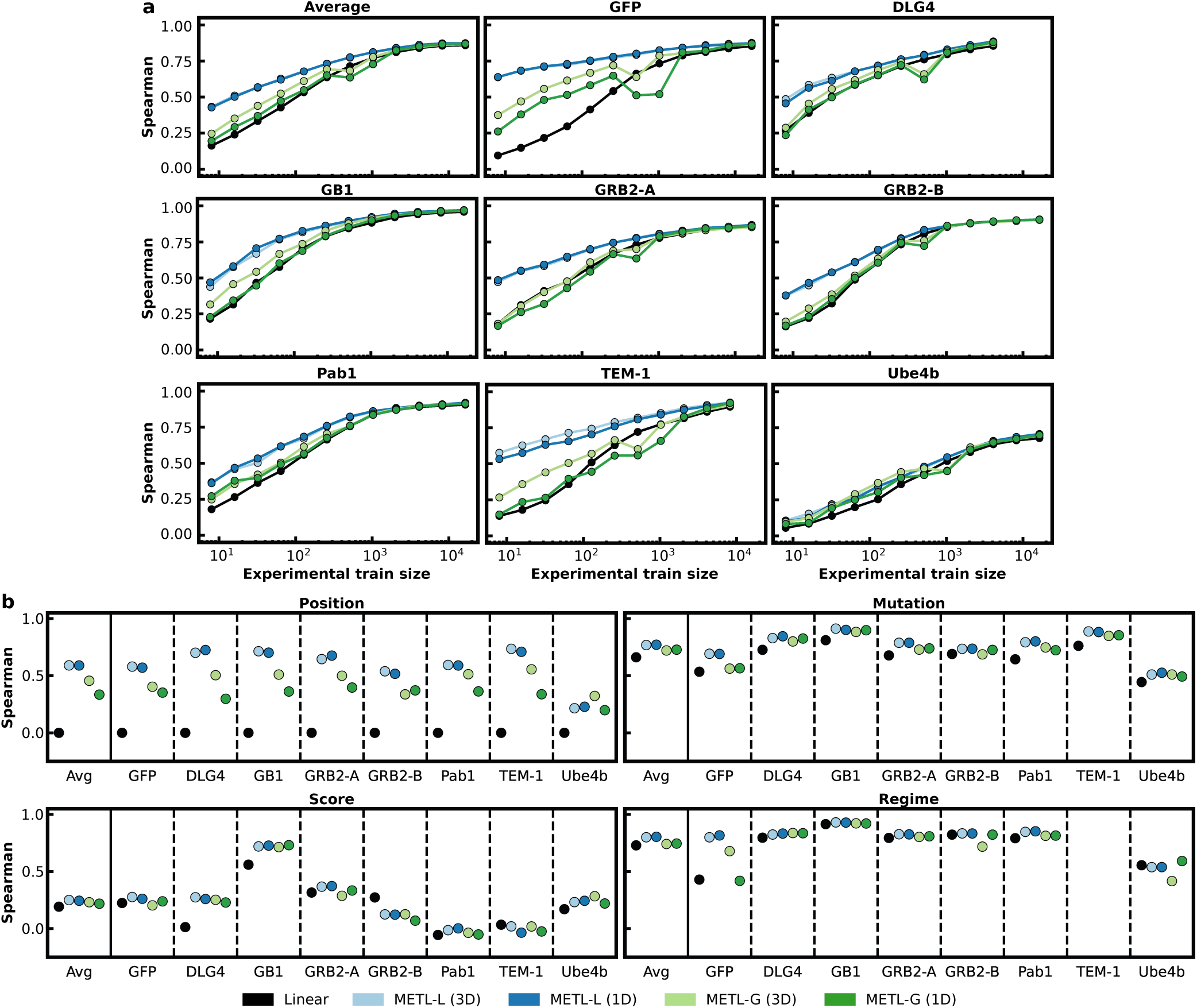
Performance of one-dimensional and three-dimensional relative position embeddings. This figure shows the performance of METL-Local and METL-Global with one-dimensional (1D) sequence-based and three-dimensional (3D) structure-based relative position embeddings. (a) Learning curves showing Spearman correlation between true and predicted scores across a range of training set sizes. (b) Spearman correlation between true and predicted scores for position, mutation, score, and regime extrapolation. Overall, METL-Local does not benefit much from three-dimensional embeddings over one-dimensional, while METL-Global shows consistent improvement with the three-dimensional embeddings. This analysis does not use all of the deep mutational scanning datasets.

**Figure S10.**
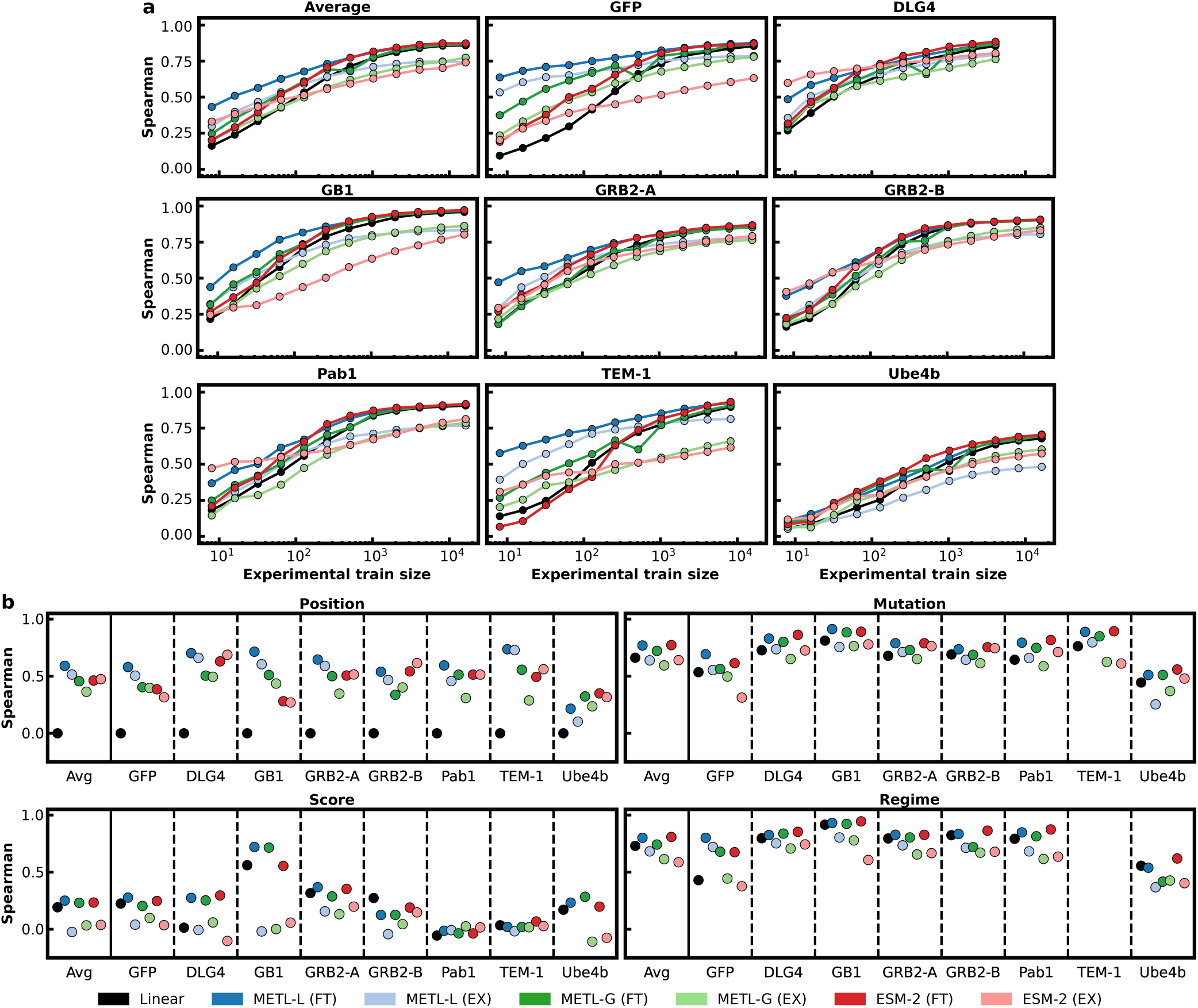
Performance of PLM finetuning and feature extraction. This figure shows the performance of METL-Local, METL-Global, and ESM-2 with both finetuning (FT) and feature extraction (EX). To perform feature extraction, we saved outputs from the appropriate internal layer of each model and then used those features as inputs to train linear ridge regression. Finetuning consistently outperformed feature extraction for METL-Local and METL-Global across (a) different training set sizes and (b) extrapolation tasks. For ESM-2, there were several instances where feature extraction substantially outperformed finetuning when applied to (a) small training set sizes, namely for the DLG4, GRB2-B, Pab1, and TEM-1 datasets. Notably, the performance of ESM-2 feature extraction exceeded the performance METL-Local finetuning for DLG4 and Pab1 with the smallest training set sizes. For (b) extrapolation tasks, ESM-2 finetuning generally performed better than feature extraction. This analysis does not use all of the deep mutational scanning datasets.

**Figure S11.**
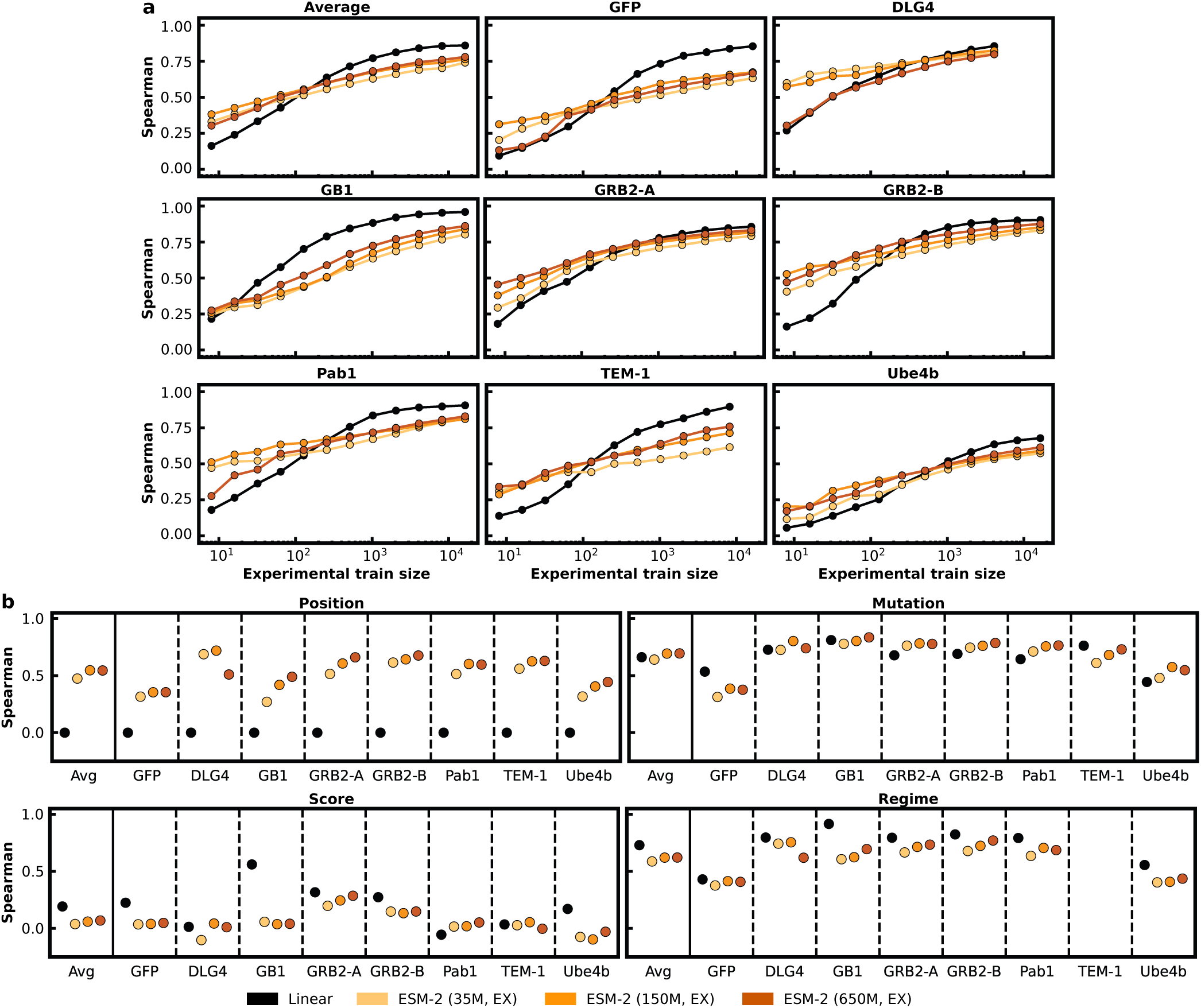
Feature extraction performance of ESM-2 models with 35M, 150M, and 650M parameters. (a) Across the range of training set sizes, the 150M parameter model consistently outperformed the 35M parameter model, with the exception of the DLG4 dataset, where the 35M parameter model performed better. Surprisingly, for small training set sizes, the 650M parameter model performed worse than both the 35M and 150M parameter models with the GFP, DLG4, and Pab1 datasets. For larger training set sizes, the 650M parameter model offered some improvement over the 35M and 150M parameter models with the GB1, GRB2-A, and GRB2-B datasets. However, in all cases Linear was the best model with larger datasets. (b) Across extrapolation tasks, the 35M parameter model tended to perform worse than the 150M and 650M parameter models. The 650M parameter model often performed the best, but not in all instances, and the differences between the models were minor in some cases. The Linear baseline was better than the feature extraction ESM-2 models on average for score and regime extrapolation. This analysis does not use all of the deep mutational scanning datasets.

**Figure S12.**
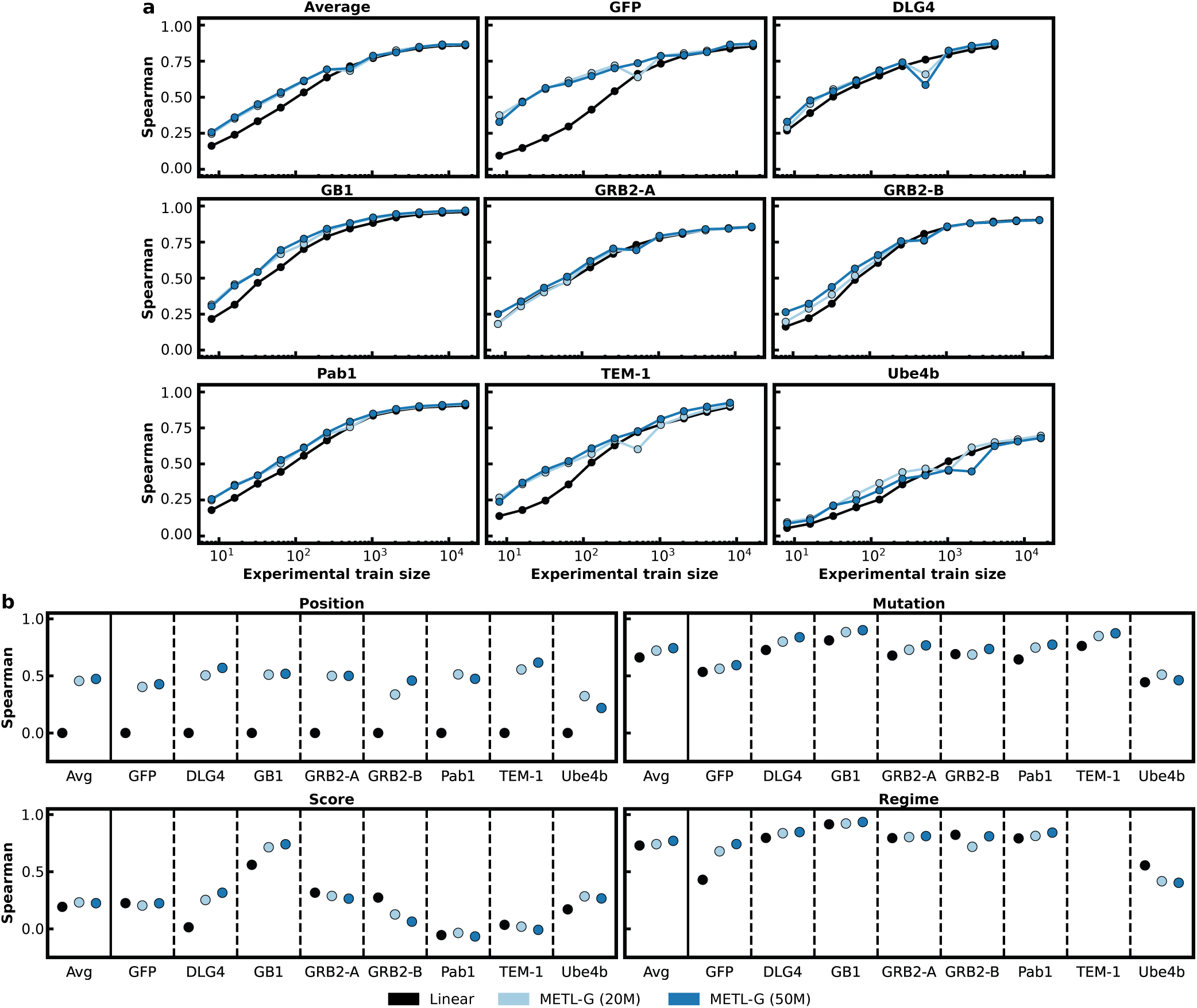
Performance of METL-Global with 20M and 50M parameters. (a) Across different training set sizes, the 50M parameter model performed similarly to the 20M parameter model on average. (b) For position, mutation, and regime extrapolation, the 50M parameter model performed slightly better on average than the 20M parameter model. For score extrapolation, the two models performed similarly on average. This analysis does not use all of the deep mutational scanning datasets.

**Figure S13.**
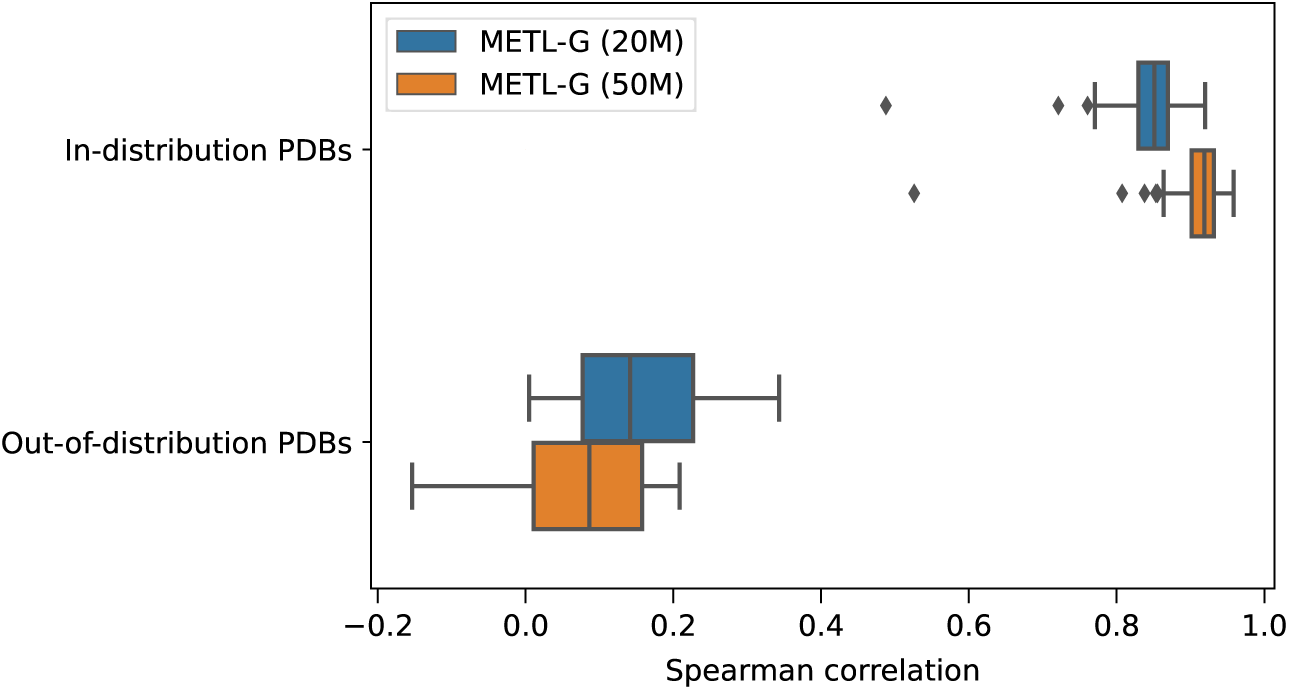
Performance of METL-Global source models predicting Rosetta *total score*. This figure shows the performance of 20M and 50M parameter METL-Global source models on predicting Rosetta *total score* for both in-distribution and out-of-distribution PDBs. In-distribution PDBs are the *n* = 148 PDBs that were used as part of the METL-Global pretraining data, while out-of-distribution PDBs consist of the *n* = 8 experimental dataset PDBs, which were not used for METL-Global pretraining. The 50M parameter METL-Global model overfits more than the 20M parameter model when predicting Rosetta *total score* on in-distribution PDBs, and it generalizes worse to out-of-distribution PDBs.

**Figure S14.**
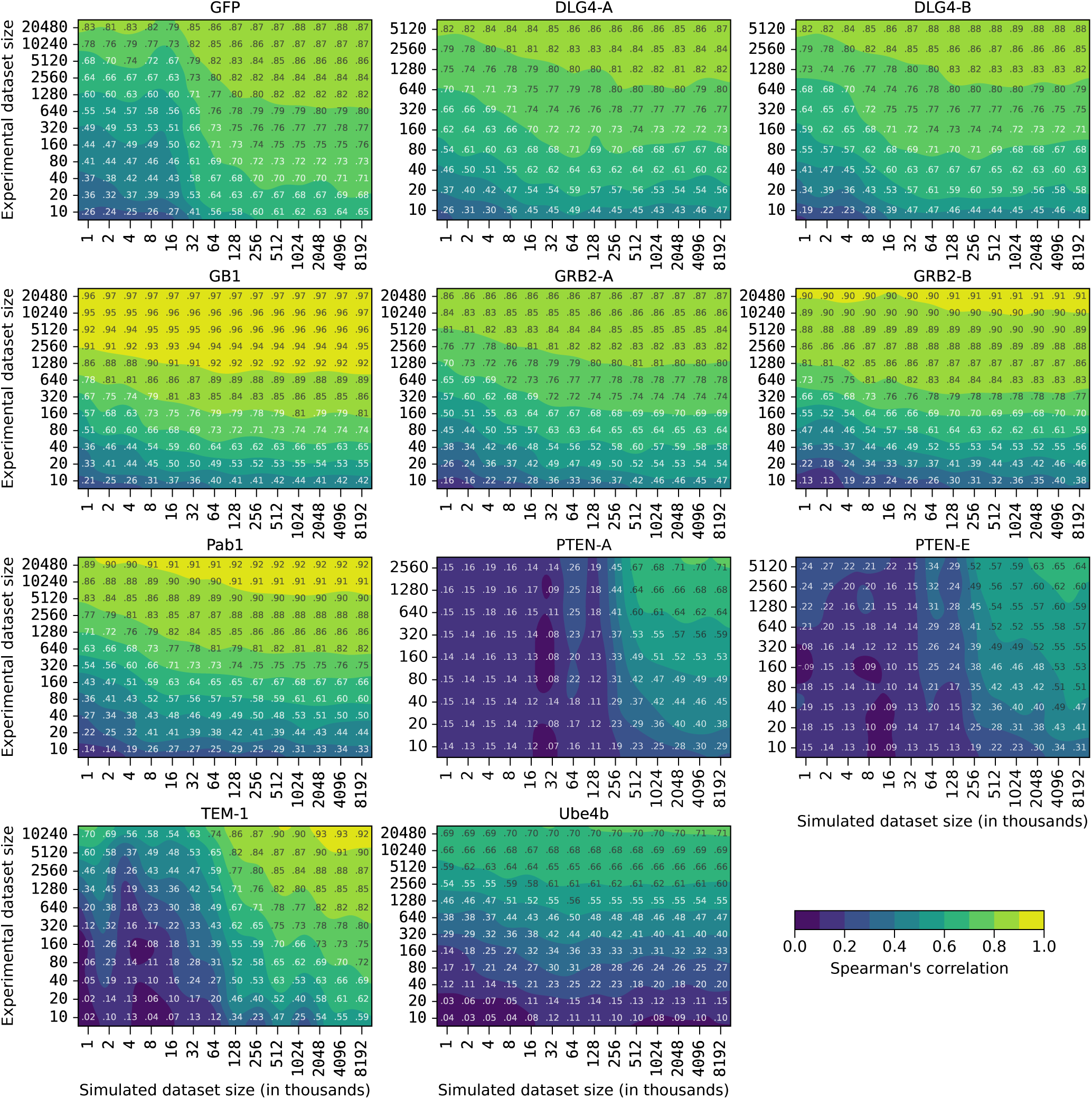
Relationships between experimental and simulated data quantities. These contour plots illustrate the test set Spearman’s correlation resulting from training METL-Local with varying amounts of simulated (pretraining) and experimental (finetuning) data. The plots display a grid of Spearman’s correlation values corresponding to discrete combinations of experimental and simulated dataset sizes.

**Figure S15.**
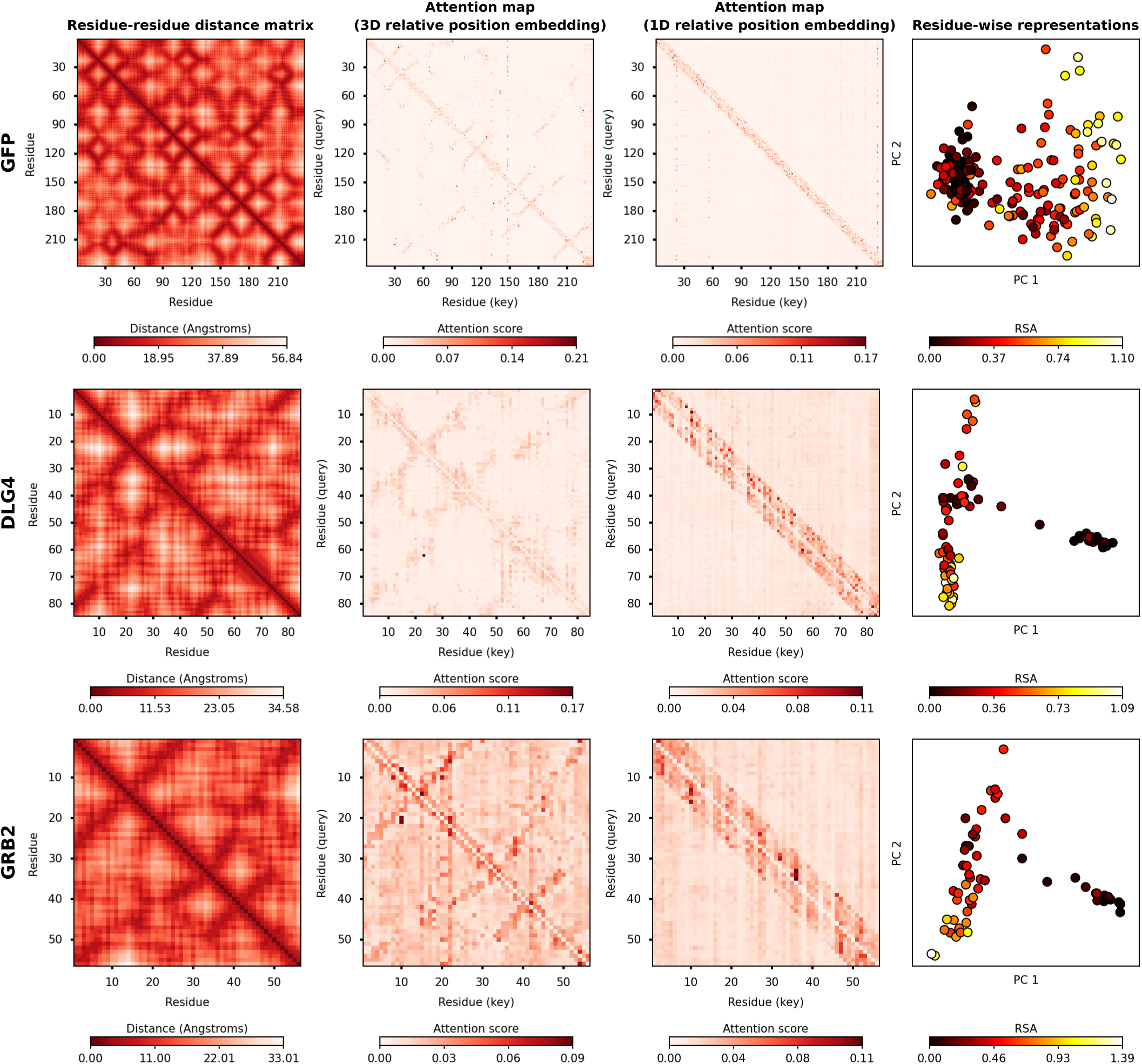
METL attention maps and residue representations for GFP, DLG4, and GRB2. The residue distance matrix shows C*β* distances between residues for the wild-type structure. The attention maps show the mean attention across layers and attention heads for the wild-type sequence when it is fed as input to the pretrained METL-Local model. The residue-wise representations show the principal component analysis (PCA) of the residue representations output by the pretrained METL-Local model, averaged across the 20 possible amino acids at each sequence position. Points are colored according to relative solvent accessibility (RSA) computed from the wild-type structure.

**Figure S16.**
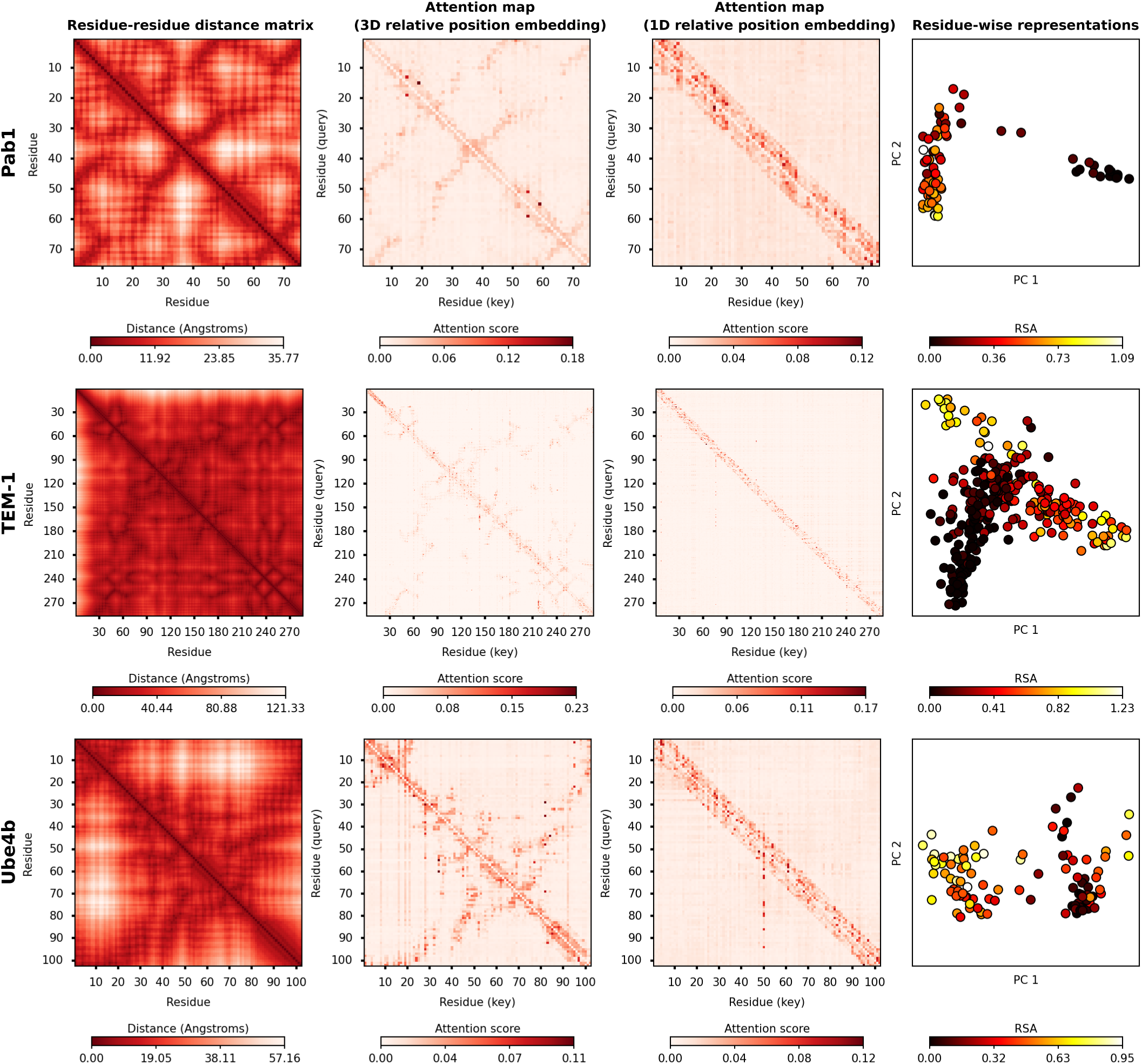
METL attention maps and residue representations for Pab1, TEM-1, and Ube4b. The residue distance matrix shows C*β* distances between residues for the wild-type structure. The attention maps show the mean attention across layers and attention heads for the wild-type sequence when it is fed as input to the pretrained METL-Local model. The residue-wise representations show the principal component analysis (PCA) of the residue representations output by the pretrained METL-Local model, averaged across the 20 possible amino acids at each sequence position. Points are colored according to relative solvent accessibility (RSA) computed from the wild-type structure.

**Figure S17.**
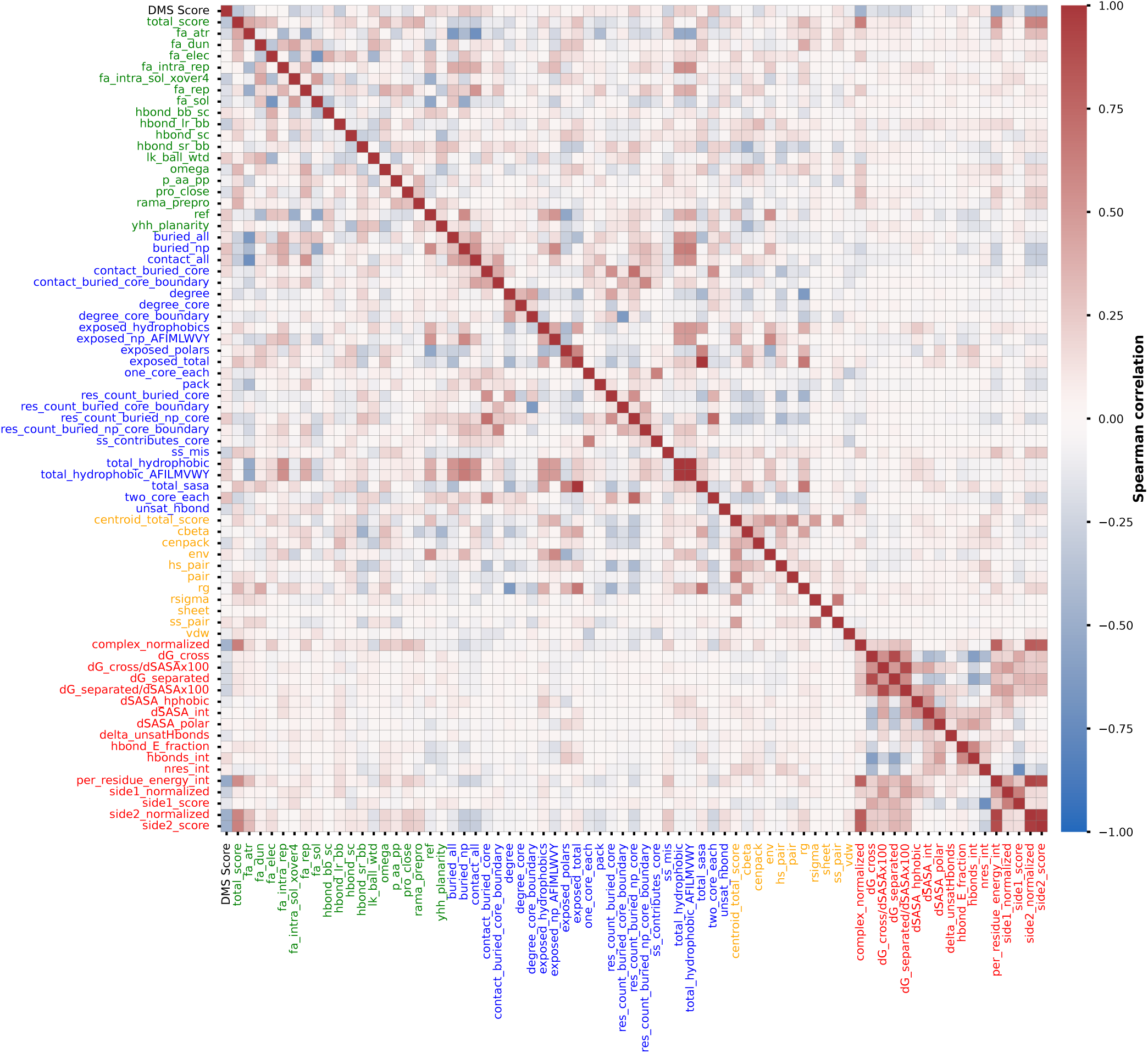
Pairwise correlations between GB1 DMS score and Rosetta scores. Heatmap showing pairwise Spearman correlations between the GB1 experimental functional score (DMS Score) and Rosetta score terms. Rosetta scores are color coded, with green representing all-atom REF15 scores, blue representing filter scores, orange representing centroid score3 scores, and red representing InterfaceAnalyzer binding scores. Correlations were computed using the GB1 DMS variants.

**Figure S18.**
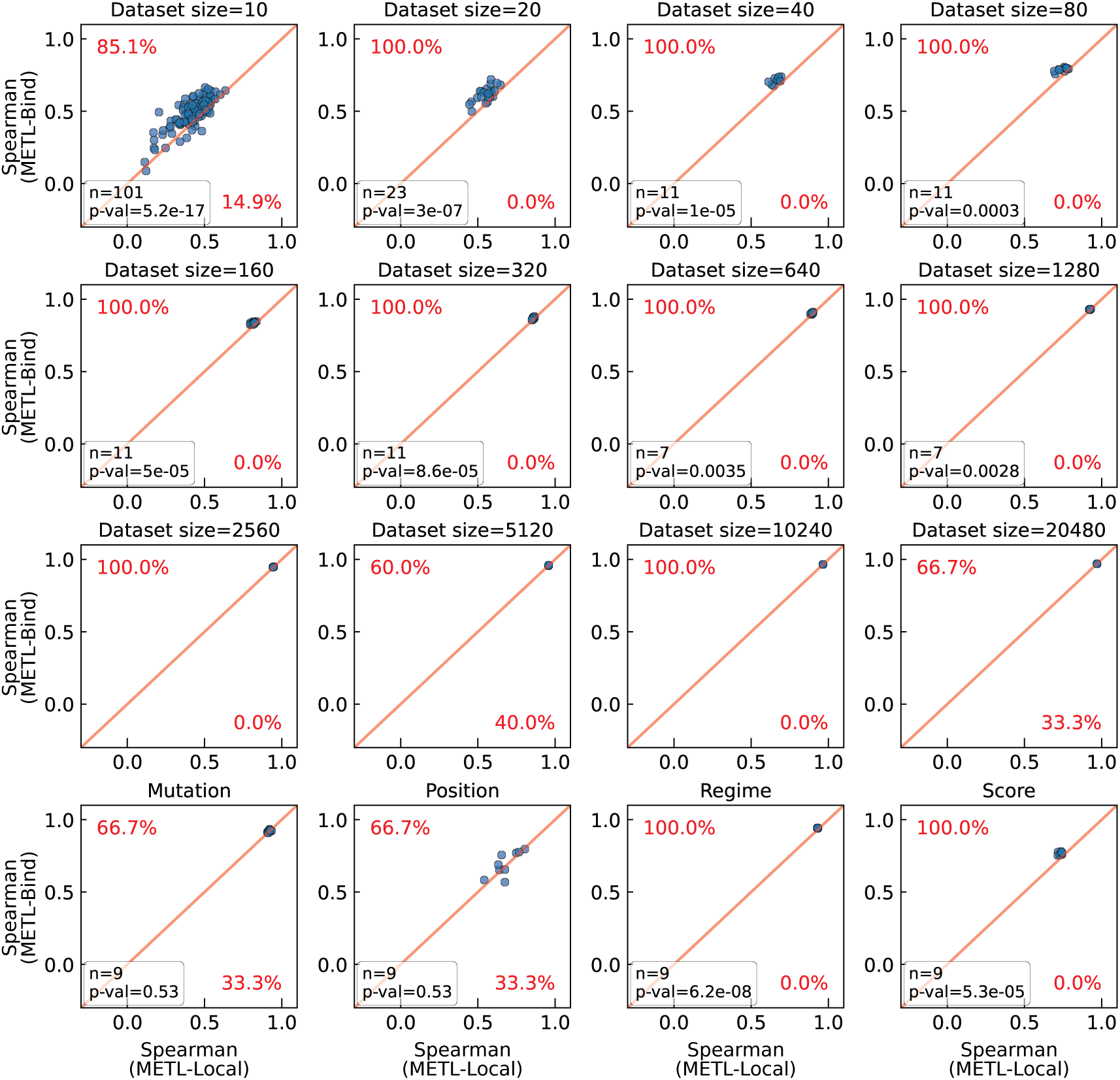
METL-Local and METL-Bind performance for individual training replicates. These scatterplots show the METL-Local and METL-Bind Spearman correlation performance for individual training replicates across different experimental dataset sizes and extrapolation tasks. The text indicates the percentage of replicates where METL-Bind performs better (upper left) or METL-Local performs better (lower right). For dataset sizes with at least 7 replicates (*n* ≥ 7), the plots are annotated with the p-value from a paired t-test, which evaluates whether the observed differences in mean performance between the two methods are statistically significant. These statistical tests confirm that METL-Bind’s improvements over METL are statistically significant (*p* ≤ 0.01) across all training set sizes with *n* ≥ 7 and for regime and score extrapolation even though the effect size can be small. Paired t-test results are summarized as: dataset size/task: *t*(df), Δ [95% CI]. Here, Δ denotes the mean difference in Spearman correlation (METL-Bind – METL-Local), and the bracketed values indicate the 95% confidence interval. Results: 10: *t*(100) = 10.13, Δ = 0.068 [0.055, 0.081]; 20: *t*(22) = 7.23, Δ = 0.064 [0.046, 0.082]; 40: *t*(10) = 8.15, Δ = 0.050 [0.036, 0.063]; 80: *t*(10) = 5.40, Δ = 0.041 [0.024, 0.058]; 160: *t*(10) = 6.76, Δ = 0.020 [0.013, 0.027]; 320: *t*(10) = 6.33, Δ = 0.009 [0.006, 0.013]; 640: *t*(6) = 4.64, Δ = 0.008 [0.004, 0.013]; 1280: *t*(6) = 4.86, Δ = 0.005 [0.003, 0.008]; mutation: *t*(8) = 0.65, Δ = 0.002 [–0.004, 0.007]; position: *t*(8) = 0.66, Δ = 0.013 [–0.032, 0.057]; score: *t*(8) = 7.78, Δ = 0.033 [0.023, 0.043]; regime: *t*(8) = 18.95, Δ = 0.011 [0.010, 0.013].

**Figure S19.**
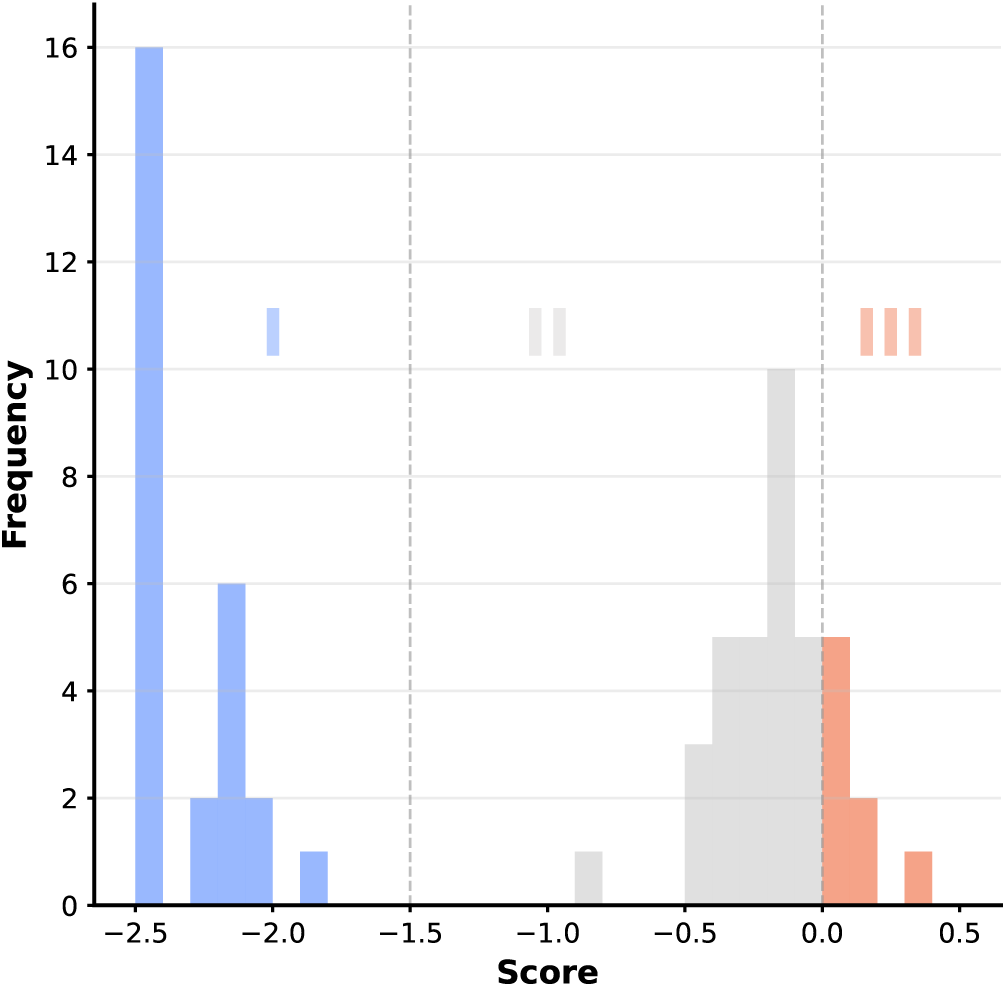
Score distribution of the 64 GFP variants. The 64 GFP variants randomly selected for METL-L-GFP training were split by score into three bins for visualization purposes. The bins were manually defined based on one threshold separating the two main modes of the distribution and another at the wild type score of 0. There are only eight variants in the bin with positive scores. The score represents the variant’s brightness.

**Figure S20.**
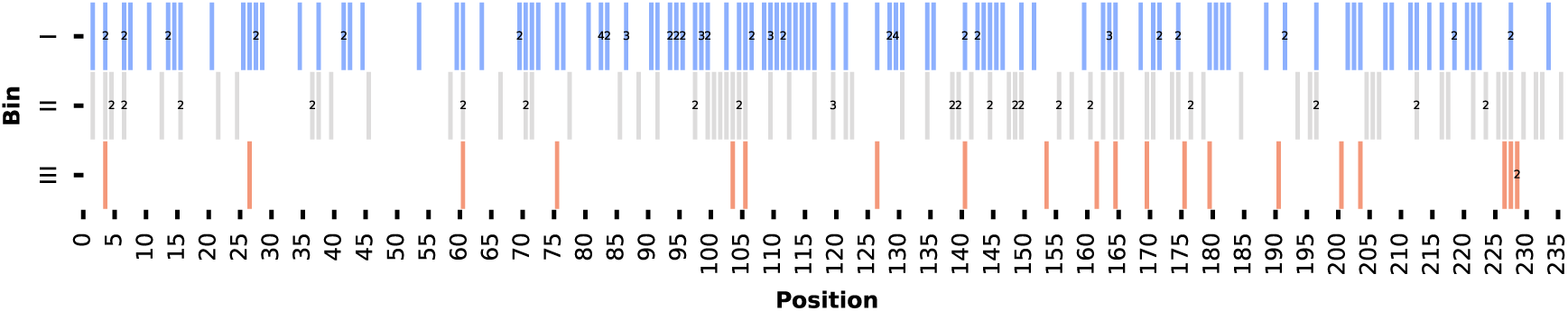
The positions of all 209 unique mutations for the 64 GFP variants. The mutations are divided using the same binning procedure and bin labels as Fig. S19. If there were multiple mutations at a position within a bin, it is marked with a numeric label. White indicates that no mutation is present at that position. A bar with no numeric label indicates there is only one mutation present at that position.

**Figure S21.**
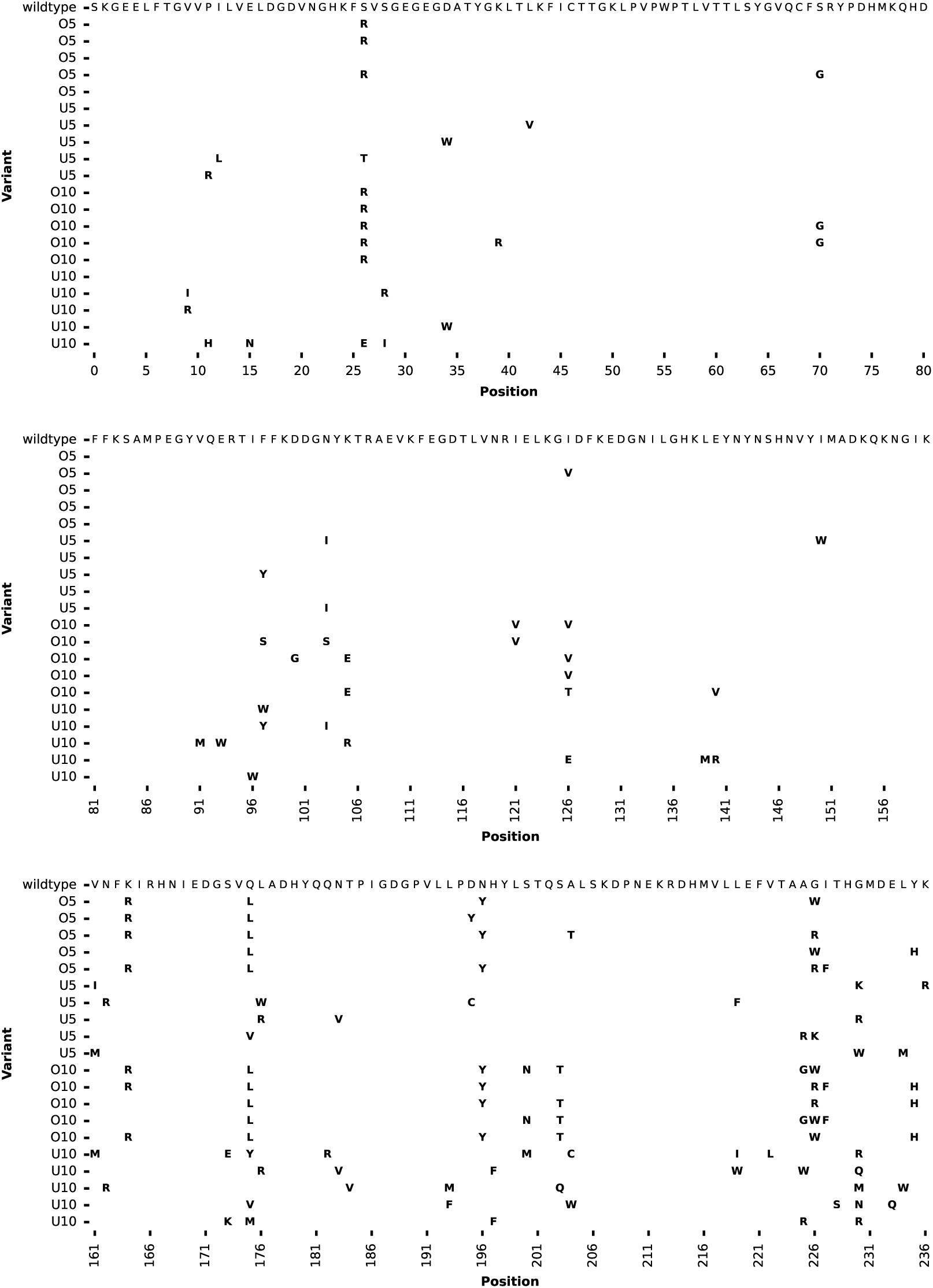
The 20 engineered GFP variants. Each row corresponds to a variant, where if a mutation was made it is shown in bold at the corresponding sequence position. O5 corresponds to Observed 5 mutant designs, U5 Unobserved 5 mutants, O10 Observed 10 mutants, and U10 Unobserved 10 mutants. The wild-type GFP sequence is shown in the top row.

**Figure S22.**
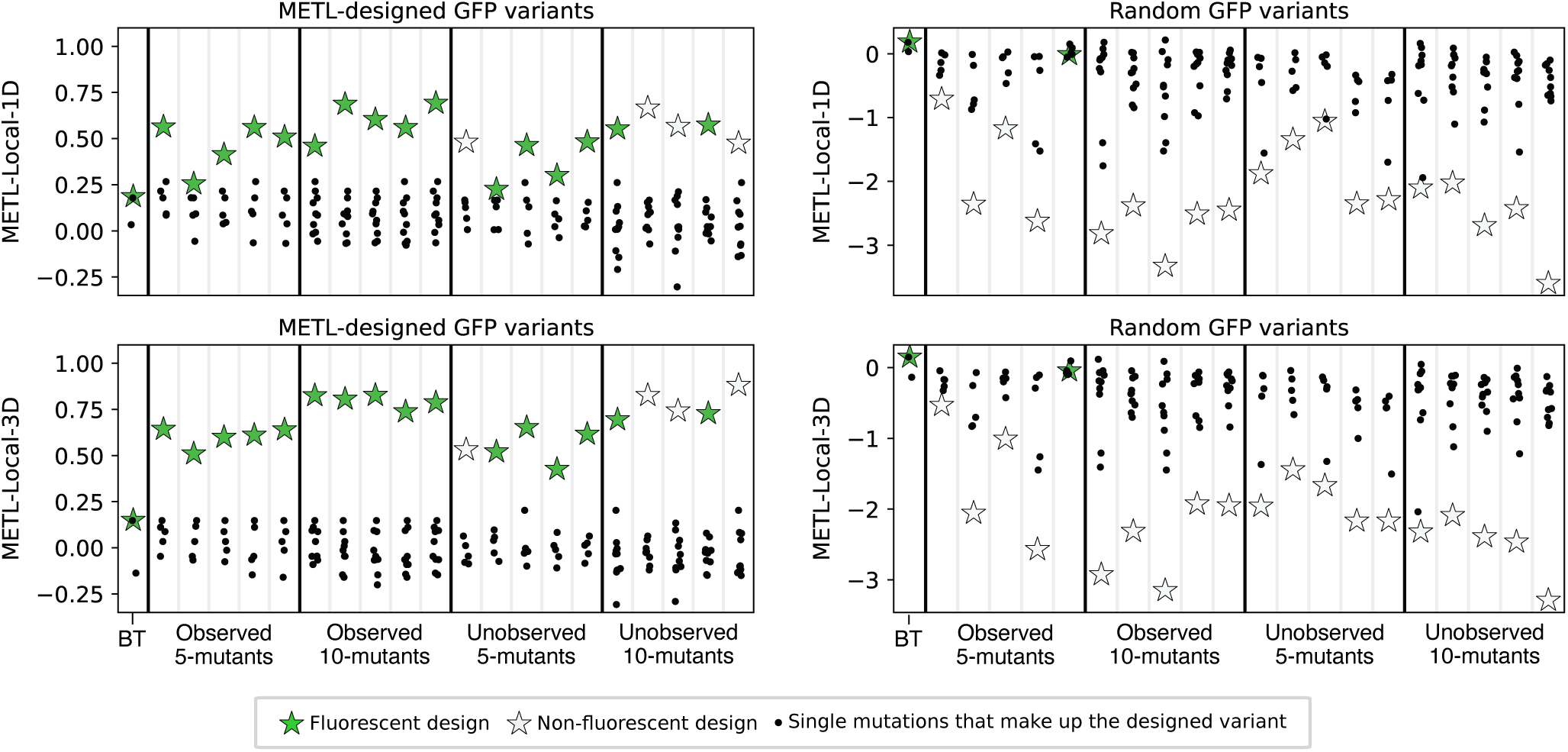
METL predictions for experimentally characterized GFP variants. METL-Local score predictions are shown for both METL-derived and random GFP variants that we experimentally characterized. Predictions from METL-Local models using 1D and 3D relative position embeddings are included. Stars represent predicted scores for the full 5- or 10-mutation variants, whereas black dots indicate the predicted scores for the single-mutation variants that compose these multi-mutation designs. BT represents the METL-Local training set variant with the highest assay score from the DMS dataset.

**Figure S23.**
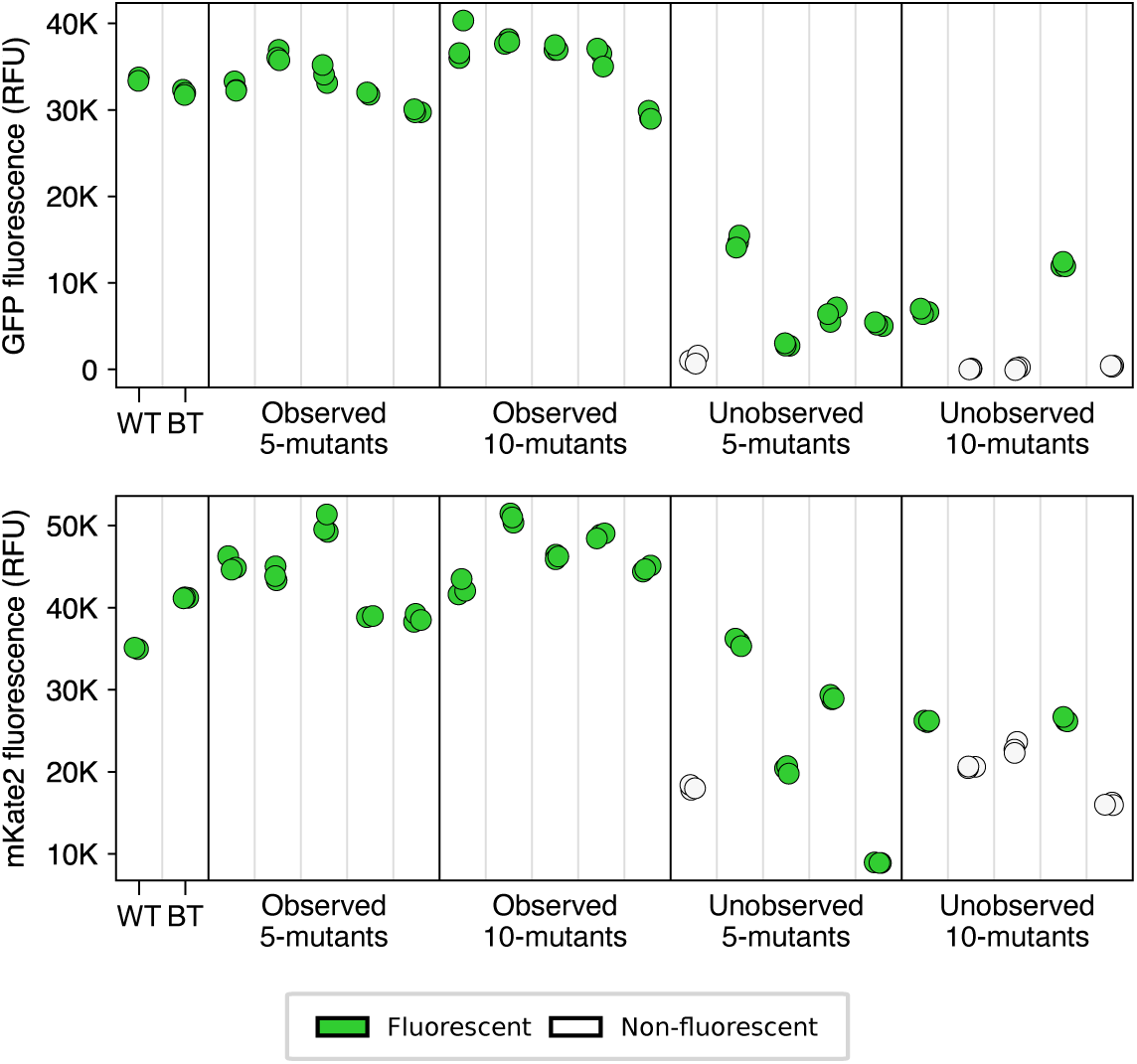
Experimental GFP and mKate2 fluorescence for engineered GFP variants. We expressed the GFP variants as fusion proteins with mKate2. The mKate2 sequence remained constant across the different GFP variants. These plots show GFP and mKate2 fluorescence normalized to optical density and with background fluorescence in negative control subtracted out. The best training set sequence (BT) and the wild-type sequence (WT) are included. Variants are colored according to whether they exhibited GFP fluorescence. Multiple replicates are shown. Because the mKate2 sequence remained constant, variation in mKate2 fluorescence may be due to changes in GFP stability.

**Figure S24.**
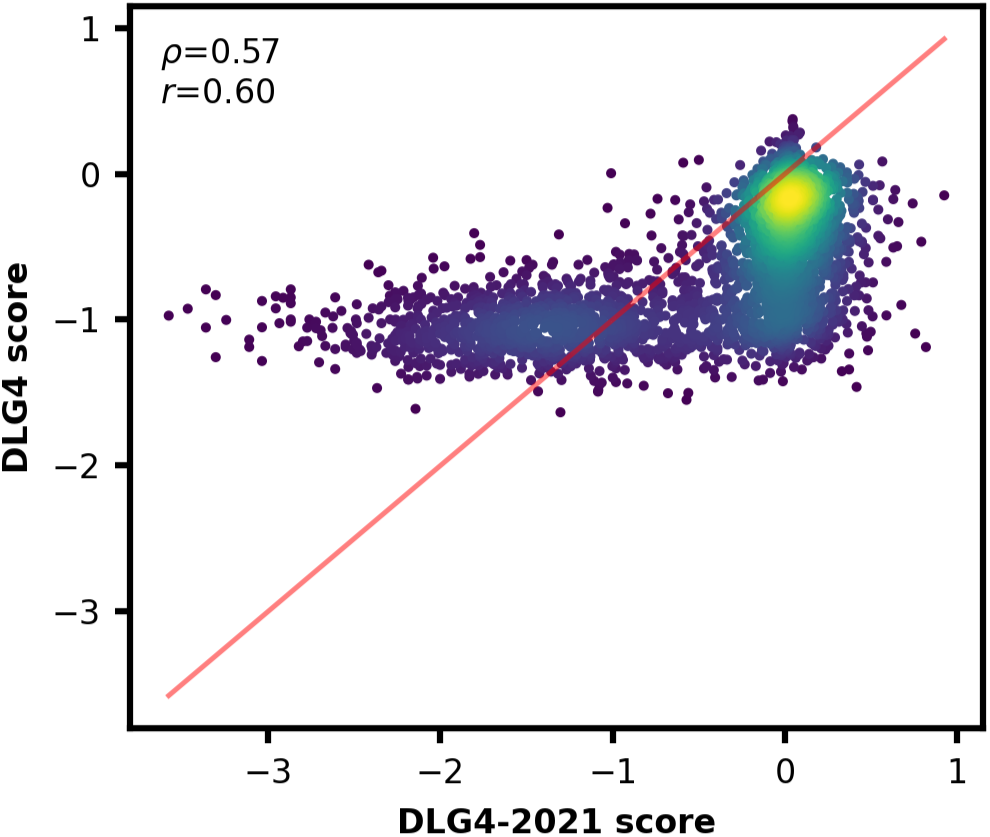
Correlation between DLG4 and DLG4-2021 dataset scores for 3,825 intersecting variants. These datasets both assayed PSD-95 PDZ3 binding to CRIPT, yet they disagree on scores, suggesting differences in methodology. We used DLG4 in our main analysis.

**Figure S25.**
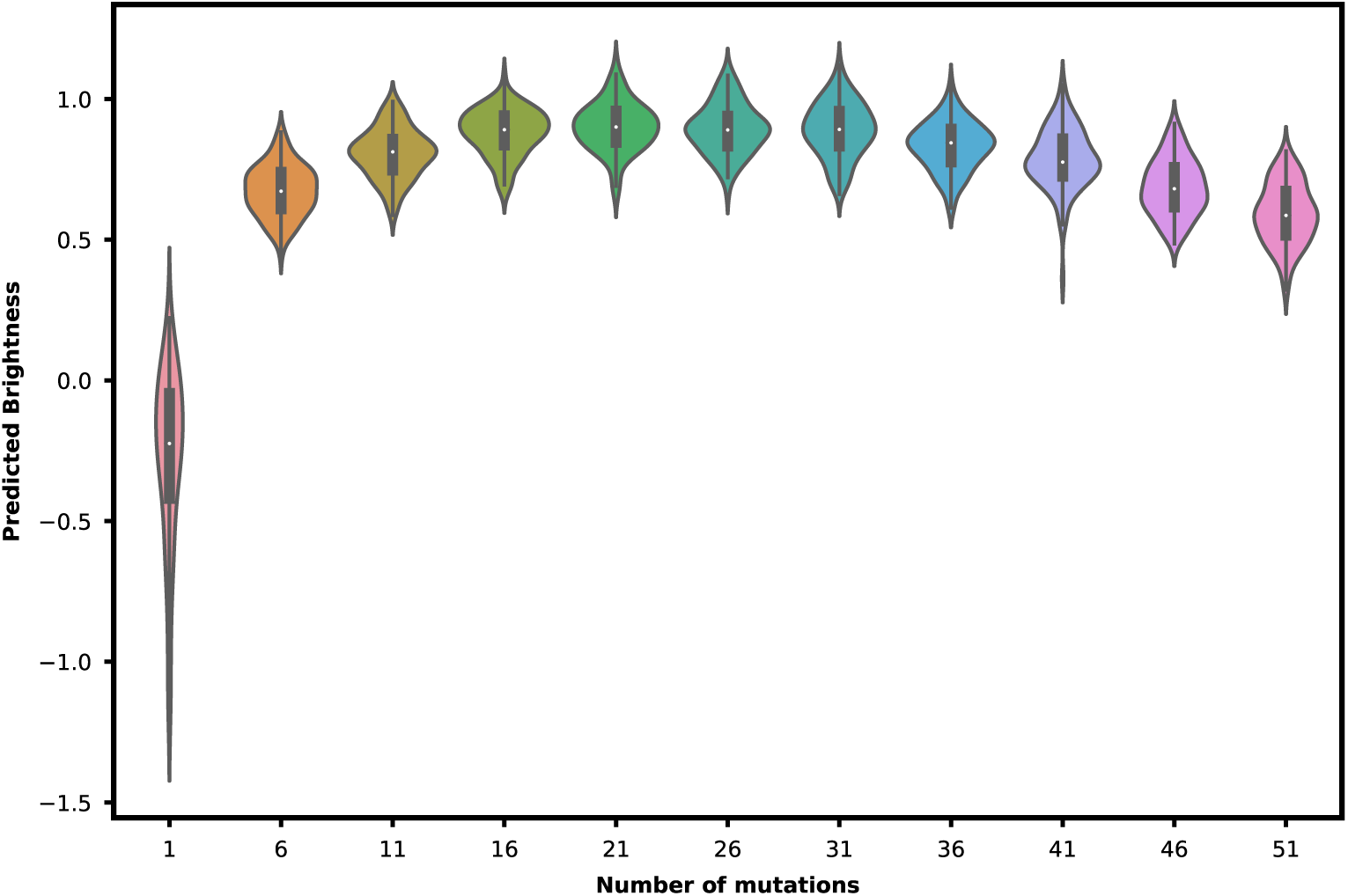
Distribution of METL-L-GFP predicted brightness of best variant found when increasing number of optimized mutations. We ran simulated annealing at different mutational distances from wild type using the same procedure used to design the 20 GFP variants with one exception. Instead of running simulated annealing 10,000 times for each mutation distance, we only ran it 100 times. The distribution of METL-L-GFP predicted brightness scores does not continue to increase as the number of mutations increases.

**Figure S26.**
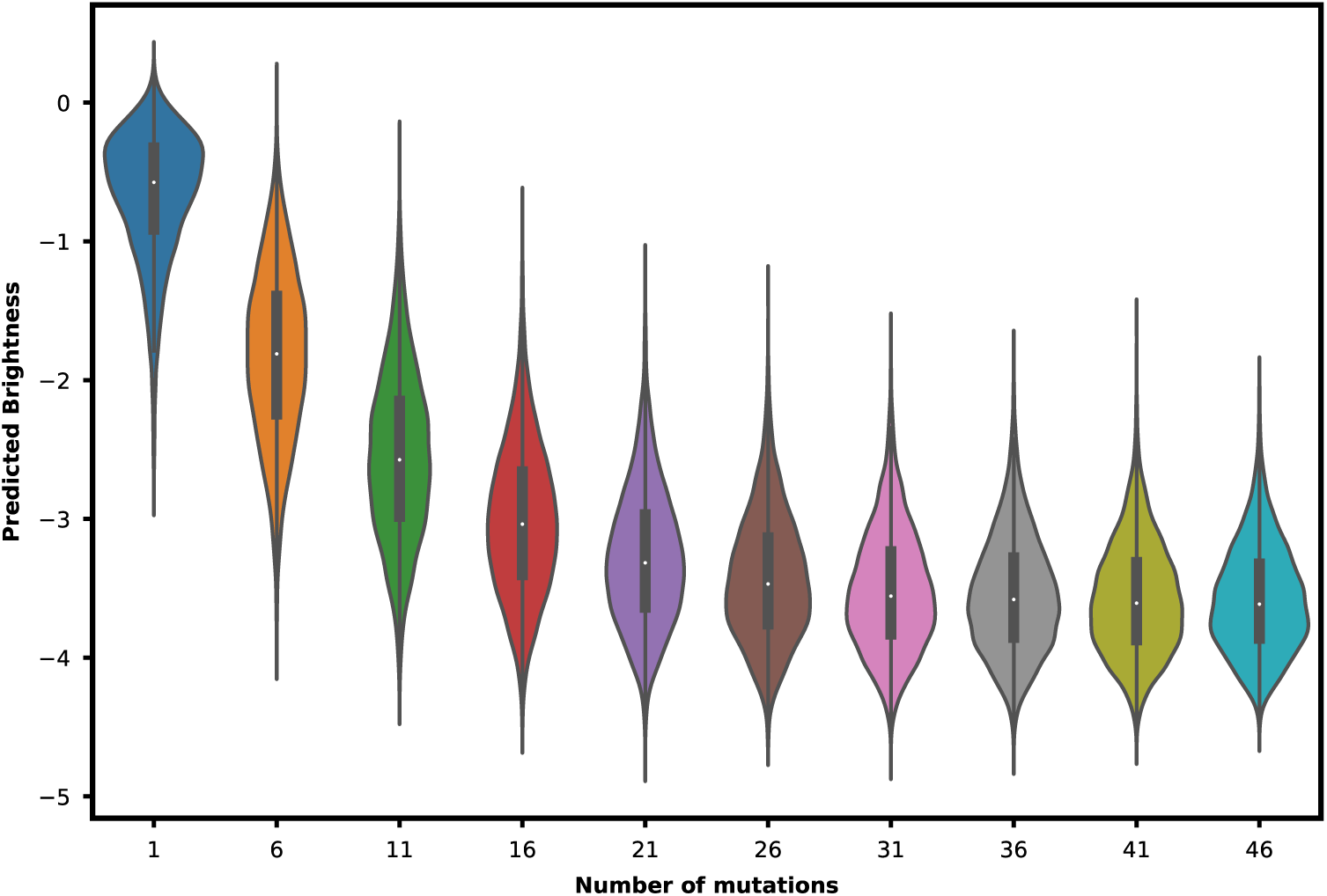
Distribution of METL-L-GFP predicted brightness for increasing number of random mutations. For each mutational distance from wild type, we randomly selected 10,000 variants and calculated their predicted brightness scores. At higher mutational distances, METL-L-GFP predicts lower brightness scores. However, the predicted brightness scores stabilize and do not continue decreasing.

**Table S1.** Rosetta score terms. The Rosetta score terms used to train METL.

| Rosetta score term | Description | Rosetta score term | Description |
| --- | --- | --- | --- |
| total_score | REF15 | exposed_np_AFIMLWVY | Custom |
| fa_atr | REF15 | exposed_polars | Custom |
| fa_dun | REF15 | exposed_total | Custom |
| fa_elec | REF15 | one_core_each | Custom |
| fa_intra_rep | REF15 | pack | Custom |
| fa_intra_sol_xover4 | REF15 | res_count_buried_core | Custom |
| fa_rep | REF15 | res_count_buried_core_boundary | Custom |
| fa_sol | REF15 | res_count_buried_np_core | Custom |
| hbond_bb_sc | REF15 | res_count_buried_np_core_boundary | Custom |
| hbond_lr_bb | REF15 | ss_contributes_core | Custom |
| hbond_sc | REF15 | ss_mis | Custom |
| hbond_sr_bb | REF15 | total_hydrophobic | Custom |
| lk_ball_wtd | REF15 | total_hydrophobic_AFILMVWY | Custom |
| omega | REF15 | total_sasa | Custom |
| p_aa_pp | REF15 | two_core_each | Custom |
| pro_close | REF15 | unsat_hbond | Custom |
| rama_prepro | REF15 | centroid_total_score | Centroid |
| ref | REF15 | cbeta | Centroid |
| yhh_planarity | REF15 | cenpack | Centroid |
| buried_all | Custom | env | Centroid |
| buried_np | Custom | hs_pair | Centroid |
| contact_all | Custom | pair | Centroid |
| contact_buried_core | Custom | rg | Centroid |
| contact_buried_core_boundary | Custom | rsigma | Centroid |
| degree | Custom | sheet | Centroid |
| degree_core | Custom | ss_pair | Centroid |
| degree_core_boundary | Custom | vdw | Centroid |
| exposed_hydrophobics | Custom |  |  |

**Table S2.**
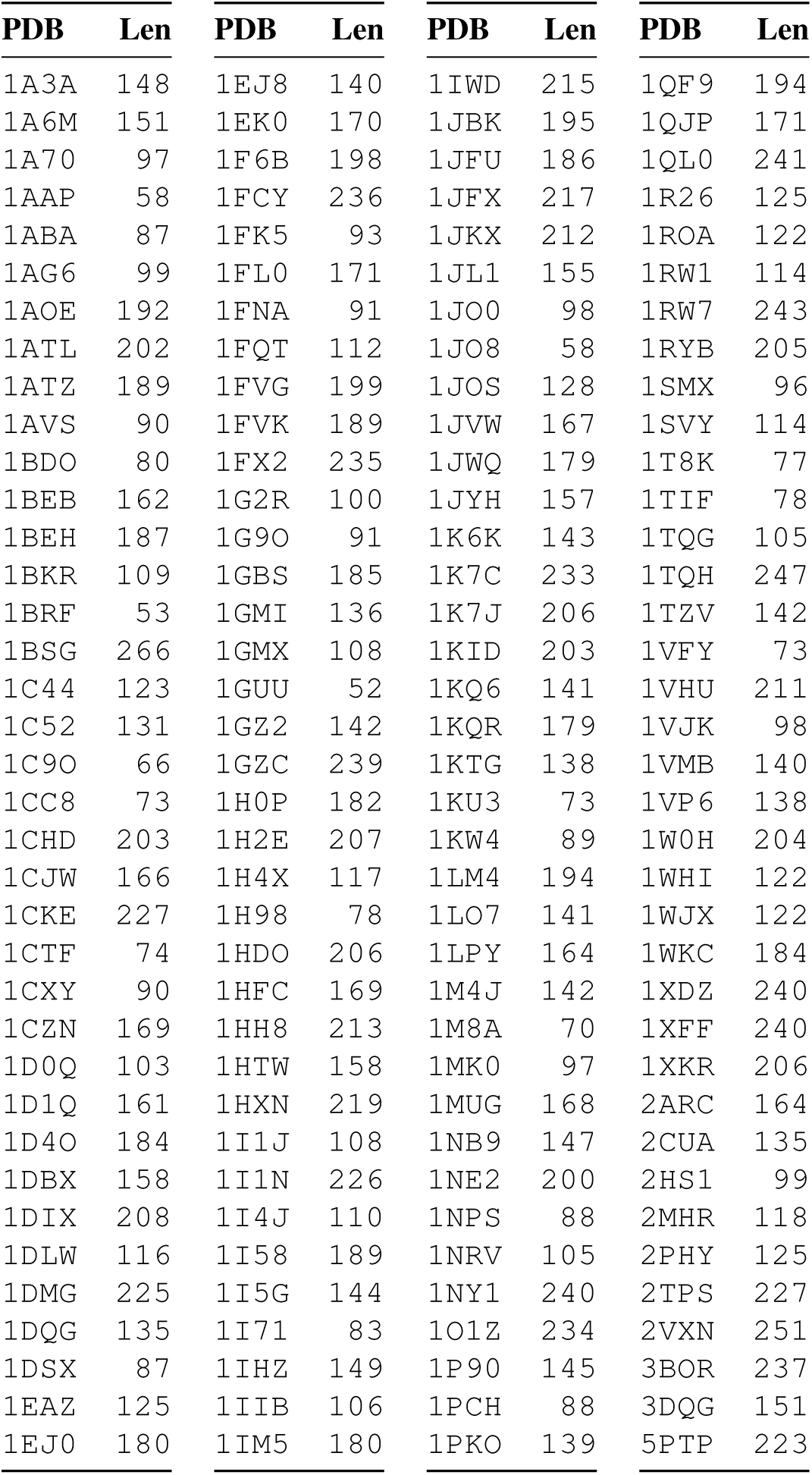
METL-Global training PDBs. The 148 base PDBs used for the METL-Global simulated pretraining data and their sequence lengths.

**Table S3.**
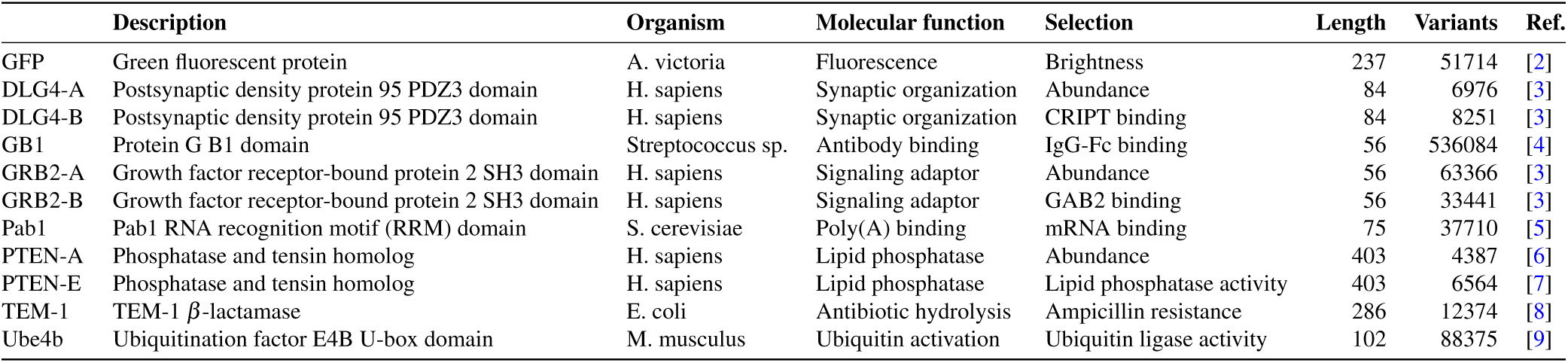
Experimental datasets. We evaluated METL on experimental datasets representing proteins of varying sizes, folds, and functions.

**Table S4.** Similarity between METL-Global pretraining structures and the downstream evaluation proteins. We clustered protein sequences and structures and used the RCSB PDB pairwise structure alignment tool to compare the METL-Global pretraining structures with the downstream evaluation proteins. Root-Mean-Square-Deviation (RMSD) is reported in Ångstroms. Template modeling score (TM-score) ranges from 0 to 1, with higher scores indicating stronger similarity. Identity is the percent sequence identity.

| Protein | Eval structure | Pretrain struct. | RMSD | TM-score | Identity | Aligned residues | Eval len | Pretrain len |
| --- | --- | --- | --- | --- | --- | --- | --- | --- |
| DLG4 | 6qji_p_trunc_2022.pdb | 1g9o_A_p.pdb | 1.79 | 0.87 | 27% | 80 | 84 | 91 |
| GRB2 | AF-P62993-F1-model_v4_trunc_p.pdb | 1jo8_A_p.pdb | 1.16 | 0.90 | 39% | 55 | 56 | 58 |
| GRB2 | AF-P62993-F1-model_v4_trunc_p.pdb | 1i1j_remod_p.pdb | 1.34 | 0.86 | 23% | 56 | 56 | 108 |
| TEM-1 | AF-Q6SJ61-F1-model_v4_p.pdb | 1bsg_p.pdb | 1.90 | 0.85 | 39% | 246 | 286 | 266 |

**Table S5.**
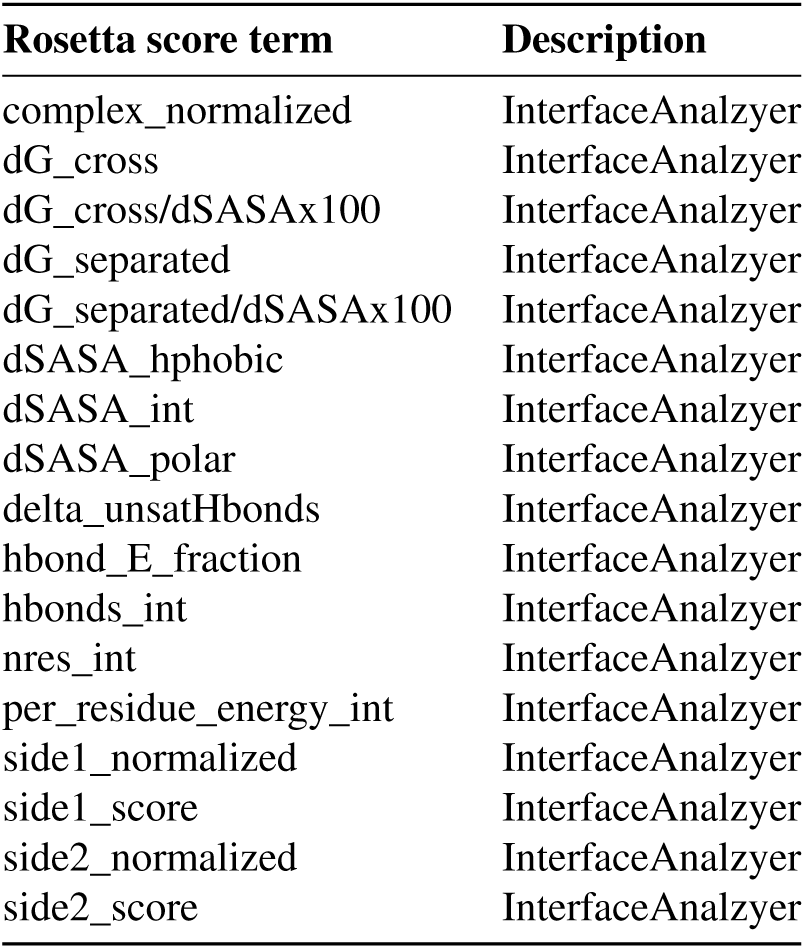
Binding score terms. The Rosetta binding score terms, calculated on the GB1-IgG complex structure and used in addition to the standard score terms to train METL-Bind.

| Rosetta score term | Description |
| --- | --- |
| complex_normalized | InterfaceAnalyzer |
| dG_cross | InterfaceAnalyzer |
| dG_cross/dSASAx100 | InterfaceAnalyzer |
| dG_separated | InterfaceAnalyzer |
| dG_separated/dSASAx100 | InterfaceAnalyzer |
| dSASA_hphobic | InterfaceAnalyzer |
| dSASA_int | InterfaceAnalyzer |
| dSASA_polar | InterfaceAnalyzer |
| delta_unsatHbonds | InterfaceAnalyzer |
| hbond_E_fraction | InterfaceAnalyzer |
| hbonds_int | InterfaceAnalyzer |
| nres_int | InterfaceAnalyzer |
| per_residue_energy_int | InterfaceAnalyzer |
| side1_normalized | InterfaceAnalyzer |
| side1_score | InterfaceAnalyzer |
| side2_normalized | InterfaceAnalyzer |
| side2_score | InterfaceAnalyzer |

**Table S6.** METL-designed GFP sequences. The METL-designed sequences in the GFP low-N design experiment.

| ID | Constraint | # Muts | Mutations |
| --- | --- | --- | --- |
| 1 | Observed | 5 | S26R, K164R, Q175L, N196Y, G226W |
| 2 | Observed | 5 | S26R, I126V, K164R, Q175L, D195Y |
| 3 | Observed | 5 | K164R, Q175L, N196Y, A204T, G226R |
| 4 | Observed | 5 | S26R, S70G, Q175L, G226W, Y235H |
| 5 | Observed | 5 | K164R, Q175L, N196Y, G226R, I227F |
| 6 | Unobserved | 5 | N103I, I150W, V161I, G230K, K236R |
| 7 | Unobserved | 5 | L42V, N162R, L176W, D195C, L219F |
| 8 | Unobserved | 5 | D34W, F97Y, L176R, N183V, G230R |
| 9 | Unobserved | 5 | I12L, S26T, Q175V, A225R, G226K |
| 10 | Unobserved | 5 | P11R, N103I, V161M, G230W, L234M |
| 11 | Observed | 10 | S26R, I121V, I126V, K164R, Q175L, N196Y, S200N, S203T, A225G, G226W |
| 12 | Observed | 10 | S26R, F97S, N103S, I121V, K164R, Q175L, N196Y, G226R, I227F, Y235H |
| 13 | Observed | 10 | S26R, S70G, D100G, K105E, I126V, Q175L, N196Y, S203T, G226R, Y235H |
| 14 | Observed | 10 | S26R, K39R, S70G, I126V, Q175L, S200N, S203T, A225G, G226W, I227F |
| 15 | Observed | 10 | S26R, K105E, I126T, E140V, K164R, Q175L, N196Y, S203T, G226W, Y235H |
| 16 | Unobserved | 10 | F97W, V161M, S173E, Q175Y, Q182R, S200M, A204C, L219I, V222L, G230R |
| 17 | Unobserved | 10 | V9I, S28R, F97Y, N103I, L176R, N183V, H197F, L219W, A225W, G230Q |
| 18 | Unobserved | 10 | V9R, V91M, E93W, K105R, N162R, T184V, L193M, S203Q, G230M, L234W |
| 19 | Unobserved | 10 | D34W, I126E, L139M, E140R, Q175V, L193F, A204W, T228S, G230N, E233Q |
| 20 | Unobserved | 10 | P11H, E15N, S26E, S28I, I96W, S173K, Q175M, H197F, A225R, G230R |

**Table S7.**
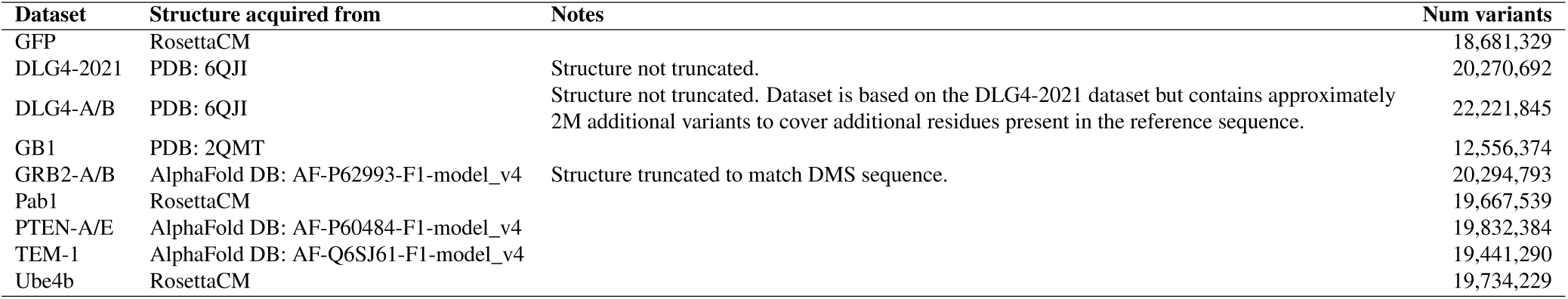
Rosetta datasets for METL-Local. Information about the Rosetta datasets used to train the METL-Local source models, including PDB origin and the final number of variants in each dataset. The DLG4-2021 dataset was not used in the main analysis.

| Dataset | Structure acquired from | Notes | Num variants |
| --- | --- | --- | --- |
| GFP | RosettaCM |  | 18,681,329 |
| DLG4-2021 | PDB: 6QJI | Structure not truncated. | 20,270,692 |
| DLG4-A/B | PDB: 6QJI | Structure not truncated. Dataset is based on the DLG4-2021 dataset but contains approximately 2M additional variants to cover additional residues present in the reference sequence. | 22,221,845 |
| GB1 | PDB: 2QMT |  | 12,556,374 |
| GRB2-A/B | AlphaFold DB: AF-P62993-F1-model_v4 | Structure truncated to match DMS sequence. | 20,294,793 |
| Pab1 | RosettaCM |  | 19,667,539 |
| PTEN-A/E | AlphaFold DB: AF-P60484-F1-model_v4 |  | 19,832,384 |
| TEM-1 | AlphaFold DB: AF-Q6SJ61-F1-model_v4 |  | 19,441,290 |
| Ube4b | RosettaCM |  | 19,734,229 |

**Table S8.** Experimental dataset preprocessing. This table specifies the experimental datasets used in this study, where we acquired them from, and any filtering or transformations we applied to standardize the dataset format. The DLG4-2021 dataset was not used in the main analysis.

|  | Acquired from | Files / URN / Accession | Variant filtering | Score transformation | Ref. |
| --- | --- | --- | --- | --- | --- |
| GFP | Paper supplement | amino_acid_genotypes_to_brightness.tsv | Drop variants with mutations to stop codons | Normalized to WT by subtracting WT score | [2] |
| DLG4-2021 | MaveDB | urn:mavedb:00000053-a | Keep if (inp >= 200) or (inp > 10 and sel >= 1) | N/A | [10] |
| DLG4-Abundance | NCBI GEO | GSE184042 | N/A | Normalized to WT by subtracting WT score | [3] |
| DLG4-Binding | NCBI GEO | GSE184042 | N/A | Normalized to WT by subtracting WT score | [3] |
| GB1 | Paper supplement | mmc2.xlsx | Keep if input_count + sel_count >= 5 | Computed from read counts w/ Enrich2 [11] | [4] |
| GRB2-Abundance | NCBI GEO | GSE184042 | N/A | Normalized to WT by subtracting WT score | [3] |
| GRB2-Binding | NCBI GEO | GSE184042 | N/A | Normalized to WT by subtracting WT score | [3] |
| Pab1 | Paper supplement | Supplementary tables 2 and 5 | N/A | Converted to log scores by taking log base 2 | [5] |
| PTEN-Abundance | Paper supplement | Supplementary table 3 | Keep only missense variants | Normalized to WT by subtracting WT score | [6] |
| PTEN-Activity | Paper supplement | Supplementary table 2 | Keep high-confidence missense; drop NaN scores | N/A | [7] |
| TEM-1 | Paper supplement | mmc2.xlsx | N/A | Converted to log scores by taking log base 2 | [8] |
| Ube4b | MaveDB | urn:mavedb:00000004-a-3 | Drop variants with mutations to stop codons | N/A | [9] |

